# Biogeochemical variability in the California Current System

**DOI:** 10.1101/2020.02.10.942565

**Authors:** Curtis Deutsch, Hartmut Frenzel, James C. McWilliams, Lionel Renault, Faycal Kessouri, Evan Howard, Jun-Hong Liang, Daniele Bianchi, Simon Yang

## Abstract

The biological productivity and diversity of the California Current System (CCS) is at the leading edge of major emerging climate trends, including hypoxia and acidification. We present results from a hindcast simulation (reanalysis) of an eddy-resolving oceanic physical-biogeochemical model of the CCS, to characterize its mean state and its patterns and drivers of variability in marine biogeochemical and ecosystem processes from 1995-2010. This is a companion paper to a physical analysis in Renault et al. (2021). The model reproduces long-term mean distributions of key ecosystem metrics, including surface nutrients and productivity and subsurface *O_2_* and carbonate undersaturation. The spatial patterns of Net Primary Productivity (NPP) are broadly consistent with measured and remotely sensed rates, and they reflect a predominant limitation by nitrogen, with seasonal and episodic limitation by *Fe* nearshore in the central CCS, and in the open ocean northern CCS. The vertical distribution of NPP is governed by the trade-off between nutrient and light limitation, a balance that reproduces and explains the observed spatial variations in the depth of the deep *Chl* maximum. The seasonal to interannual variability of biogeochemical properties and rates is also well captured by model simulations. Because of the prevailing nutrient limitation, fluctuations in the depth of the pycnocline and associated nutricline are the leading single factor explaining interannual variability in the interior biogeochemical state, and the relationships between density and biogeochemical rates and tracers are consistent between model and observations. The magnitude and relationship between density structure and biogeochemical processes is illustrated by the 1997-98 El Niño event, which faithfully reproduces the single largest deviation from the mean state in the simulated period. A slower decadal shoaling of the pycnocline also accounts for the concomitant trends in hypoxic and corrosive conditions on the shelf. The resulting variability is key to understanding the vulnerability of marine species to oceanic change, and to the detection of such changes, soon projected to exceed the range of conditions in the past century.

## 1 Introduction

Coastal upwelling systems along the eastern boundaries of the subtropical oceanic basins are some of the most climatically and biologically dynamic regions of the world’s oceans (Carr and Kearns,2003; Kudela et al., 2008). In the California Current System (CCS) of the Eastern North Pacific, the seasonal cycle of alongshore winds and offshore surface currents yields a direct conduit for nutrient-rich water to rise from the deep ocean to the sunlit surface (Chavez and Messie, 2009). The upwelling of cold water near the coast is evidenced in satellite remote sensing of sea surface temperature, expansive offshore marine stratocumulus clouds, and a nearshore ribbon of high chlorophyll from phytoplankton. The resulting cascade of phytoplankton biomass up the food chain supports high biodiversity and productive fisheries (FAO, 2009).

The physical state of the Northeastern Pacific varies on time scales of days to decades. This variability includes mesoscale eddies, seasonal cycle, interannual El Niño Southern Oscillation (ENSO) variability, and lower frequency climate fluctuations characterized by the Pacific Decadal Oscillation (PDO, Mantua et al. (1997)) and the North Pacific Gyre Oscillation (NPGO, Di Lorenzo et al. (2008)). Ecosystems throughout the Eastern Pacific respond strongly to physical forcing at each of these timescales, through the physical influence of winds, light, and heat, and their effects on the supply of nutrients to phytoplankton and oxygen to marine animals. Interannual climate cycles associated with ENSO in particular are major perturbations to these parameters and thus to plankton productivity (Chavez et al., 2002; Bograd and Lynn, 2001; Turi et al., 2018). Because thermocline waters entering the CCS are, like other subtropical Eastern Boundary Upwelling Systems, far from the sites of atmospheric ventilation in the west, the rising waters are “old” and bear the signature of decades of biogeochemical process that yield low *O*_2_, high nutrients, and low pH. The upwelling in eastern boundary systems also generates energetic mesoscale eddy fields (Capet et al., 2008). These eddies can transport the extreme properties of CCS thermocline waters far off-shore and down into the subtropical interior, connecting the biogeochemistry of the coastal CCS with the adjacent oceanic gyres (Nagai et al., 2015; Gruber et al., 2011; Renault et al., 2016a; Frenger et al., 2018).

Oceanic acidification and deoxygenation are also emerging trends in the CCS ecosystem (Gruber et al., 2012; Chan et al., 2008). Anthropogenic *CO_2_* has been detected in coastal subsurface waters off Northern California (Feely et al., 2008). Decadal trends in oxygen have also been observed in the California Current (McClatchie et al., 2010; Pierce et al., 2012; Dussin et al., 2019) and have altered the proportions of biologically important nutrients (Deutsch et al., 2011). In the Northern California Current, massive benthic die-offs have been attributed to episodes of extreme hypoxia along the Oregon shelf (Chan et al., 2008). Fluctuating abundance of species in upper trophic levels observed over decadal and longer time scales arise from climate variability, but the specific mechanisms remain obscure (Chavez et al., 2003; Rykaczewski and Checkley, 2008). In addition to climate forcing, the coastal ocean is subjected to anthropogenic pollution that could locally exacerbate hypoxia and acidification (Doney et al., 2007), and may contribute to the increasing frequency and toxicity of harmful algal blooms along California’s coast (Andersson et al.,2008).

The continental margin of western North America has a rich endowment of historical and ongoing observational programs aiding evaluation of climate-ecosystem interactions, including one of the most extensive time series programs (the California Cooperative Ocean Fisheries Investigations, CalCOFI) anywhere in the world’s oceans (Ducklow et al., 2009). The age of shallow thermocline waters also make them prone to large amplitudes of low-frequency variability (Ito and Deutsch, 2010) that confound the detection of climate and other anthropogenic changes impinging through the oceanic surface. Thus, even in this relatively well-studied region of the ocean, it is difficult to distinguish long-term trends associated with anthropogenic climate change from the low-frequency variability that pervades oceanic properties in regions of close contact with old, deep waters. For example, it remains unclear whether the changes in hypoxia in the California Current are driven by internal climate variability, or are an early sign of long-term climate warming (McClatchie et al., 2010; Pierce et al., 2012; Long et al., 2016).

Models of this dynamic region often exhibit substantial biases in the mean state and unknown fidelity in representing historical variability and its causal mechanisms. The purpose of this paper is to report a system-wide validation of eddy-resolving, regional model fields and property relationships through comparison to a variety of hydrographic, experimental, and remote sensing observations. Here we focus on biogeochemical aspects of the model solution; the physical dynamics and its model validation are discussed in a companion paper (Renault et al., 2021). In this work, we demonstrate the fidelity of modeled spatial patterns and seasonal to interannual variability to observational datasets. We further probe the mechanisms underlying this variability-better understanding these drivers is critical to attributing and projecting the biogeochemical responses of the CCS to natural climate fluctuations and anthropogenic change. Section 2 provides a description of the model, its boundary conditions, the simulations performed, and the datasets used for model evaluation. These constitute the core results described in Section 3 and summarized in the final Section 4.

## 2 Methods

### 2.1 Model description

The ecosystem and biogeochemical cycles are simulated in the Regional Ocean Modeling System (ROMS, Shchepetkin and McWilliams (2005)). As in Renault et al. (2016b), the primary domain extends from 144.7°W to 112.5°W and from 22.7°N to 51.1°N. Its grid is 437 x 662 points with a horizontal resolution of *dx* = 4 km and 60 vertical levels. Initial and horizontal boundary data for temperature, salinity, surface elevation, and horizontal velocity are taken from the quarter-degree, daily-averaged Mercator Glorys2V3 data-assimilating ocean reanalysis (accessible via http://www.myocean.eu; described further at https://www.mercator-ocean.fr/en/science-publications/glorys/). In order to maintain a realistic representation of the variability in water mass properties and transport into the model domain over time, monthly anomalies from the Mercator data are added to the mean monthly climatology from the World Ocean Atlas (WOA, Locarnini et al. (2013); Zweng et al. (2013)) over the period 1995-2004. The freshwater, turbulent heat, and momentum fluxes are estimated using bulk formulae (Large, 2006) and the atmospheric fields derived from an uncoupled simulation with the Weather Research and Forecasting model (WRF). Heat and momentum fluxes are computed from bulk formulae, as detailed in Renault et al. (2021). The freshwater flux from river runoff is included as surface precipitation and is spread using a Gaussian distribution over the grid cells that fall within the range from the coast to 150 km off-shore; this excludes a detailed representation of river plumes. The river-runoff forcing dataset we use is a monthly climatology from Dai et al. (2009). The river inputs are assumed to carry negligible chemical concentrations, except the outflow from the Juan de Fuca Strait (see below). This statistically equilibrated solution, named USW4, is integrated over the period 1995-2010 after a spin up of 1 year (with initial conditions derived from the World Ocean Atlas).

Further details are described in the companion paper (Renault et al., 2021).

The coastal biogeochemical dynamics are simulated using an ecosystem model (the Biogeochemical Elemental Cycling (BEC) model, Fig. 1 and Appendix). This model includes both phytoplankton and zooplankton, and dissolved, suspended, and sinking particulates (Moore et al.,2004). The model includes four phytoplankton functional groups (picoplankton, diatoms, coccolithophores, and diazotrophs) characterized by distinct biogeochemical functions (nutrient recycling, silicification, calcification, and N_2_ fixation, respectively). Four nutrient cycles (nitrogen, silicic acid, phosphate, and iron) are simulated and are coupled through a fixed phytoplankton stoichiometry, except for iron (Fe), which varies in proportion to the other nutrients (see Equations A100-109; Moore et al. (2002, 2004)). The ecosystem is linked to an oceanic biogeochemistry module that includes total inorganic carbon (DIC), alkalinity, iron, and dissolved *O*_2_. Remineralization of sinking organic matter is parameterized according to the mineral ballast model of Armstrong et al. (2001). Gas exchange fluxes for *O*_2_ and *CO*_2_ are based on Wanninkhof (1992). The BEC equations are listed in the Appendix, and the model code, including parameter settings, are available through the GitHub repository (see the remark at the end of the paper).

**Figure 1:**
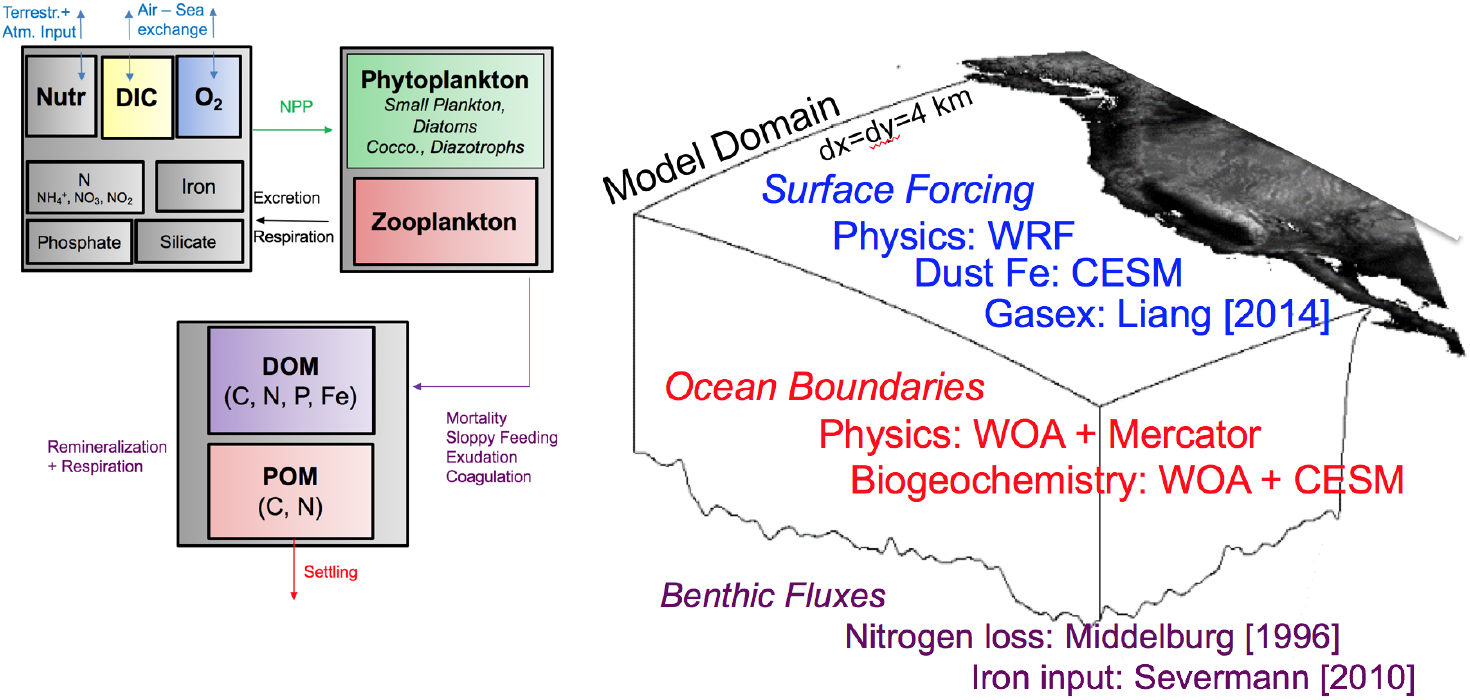
Schematic structure and physical configuration of ROMS-BEC biogeochemical model. (a) The main ecosystem state variables and fluxes. (b) Geographic scale of simulation, and sources of surface forcing, open boundary condition data and representations of benthic nutrient fluxes.

The iron *Fe* cycle includes dissolved iron, scavenged iron, and iron associated with organic matter pools and dust particles, but only dissolved iron and organically bound iron are explicitly modeled as state variables. For dissolved iron, four processes are considered: atmospheric deposition, biological uptake and remineralization, scavenging by sinking particles, and release by sediments. Atmospheric iron deposition is based on the dust climatology of Mahowald et al.(2006). We implemented a sedimentary iron source based on benthic flux chamber measurements in the California margin. An equation relating sediment *Fe* release as a function of bottom water *O*_2_(log_10_[*F_fe_*] = 2.5 - 0.0165 · *O*_2_, where *O*_2_ is in mmol m^−3^ and the efflux units are *μ* mol m^−2^ d^−1^) is derived from data compiled by Severmann et al. (2010). The resulting rates of *Fe* supply from sediments (Fig. 2) exceed those from atmospheric dust deposition throughout the model domain.

**Figure 2:**
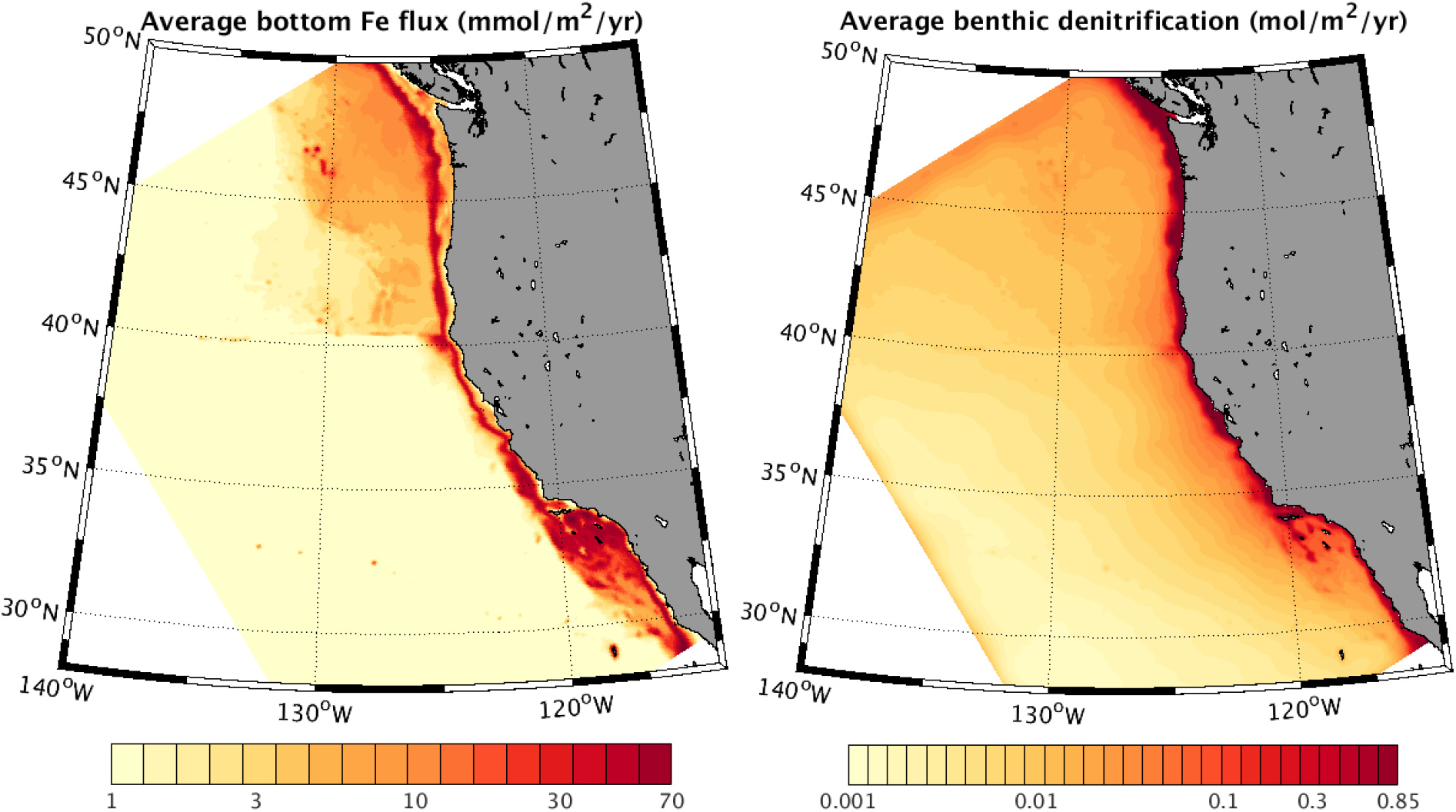
Parameterized fluxes of iron and nitrate between the water column and sediments. (a) The *Fe* efflux (in mmol m^−2^ yr^−1^) to the water column from the sediments. (b) The 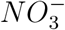 flux from the water column to the sediments due to net denitrification in sedimentary pore waters. Both fluxes are parameterized as a function of bottom water *O*_2_, and denitrification is additionally parameterized as a function of organic matter flux to the seafloor. These maps are therefore part of the model solutions, and not prescribed forcings.

We also added an anoxic nitrogen cycle, with losses to the sediments and water column. Bottom water nitrate is removed using a statistical description of sediment denitrification proposed by Middelburg et al. (1996), based on a vertically resolved diagenetic model that predicts the primary dependence of benthic denitrification to be on organic carbon sedimentation rate, with a secondary sensitivity to bottom water oxygen concentration. This statistical description of the complete diagenetic model reproduces basic controls on observed sediment fluxes, without the considerable computational cost of a sedimentary submodel. The predicted rates of 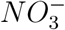 loss from this sedimentary sink (Fig. 2) amount to a small loss of ≈ 3 ×10^12^ gN yr^−1^. Denitrification in the water column is also modeled, but its integrated removal rate is an order of magnitude smaller than sedimentary losses, and has negligible impact on the results because *O*_2_ in the model domain rarely falls below the threshold (5 mmol m^−3^) assumed for this process. The higher *O*_2_thresholds associated with anaerobic particle micro-environments could increase the importance of anaerobic processes in the CCS, but they are not represented in this model (Bianchi et al., 2018). The removal of 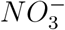by denitrification also acts as a sink for alkalinity.

### 2.2 Biogeochemical forcing and validation data

The biogeochemical model components and the physical (Renault et al., 2021) and biogeochemical forcings are schematically represented in Fig. 1. The biogeochemical boundary conditions for nutrients 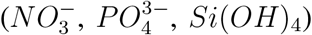 and *O*_2_ are taken from monthly climatological observations in the 2013 World Ocean Atlas (WOA) (Garcia-Reyes et al., 2014). Additionally, lateral 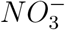 fluxes derived from prior model simulations (Davis et al., 2014) of nutrient exchange between the Strait of Juan de Fuca and the coastal ocean were imposed as landward boundary conditions in the Northern CCS. Non-nitrogenous nutrients were not available for inclusion in fluxes from the Strait of Juan de Fuca. However, we imposed an *Fe* concentration at this boundary by scaling it to nitrate. The scaling factor 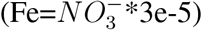 was chosen to be similar to that of the surrounding coastal water. This ensures that Juan de Fuca nutrient inputs will not alter the locally limiting nutrient, absent data to support such an alteration. Boundary condition data for *Fe* is taken from global simulations with the Community Earth System Model (CESM) that used an earlier version of the same BEC ecosystem model. The 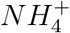 boundary concentrations, being small in nature, are set to zero, but adjust rapidly to the ecosystem processes in the interior of the domain. Time-dependent carbon cycle parameters, DIC and Alk, are taken from GLODAP (Key et al., 2004), with a reference year of 1995. An imposed trend at the boundary scales the anthropogenic component of DIC in proportion to the rise of atmospheric *CO_2_* since 1995. Time-dependent atmospheric *pCO_2_* is also used as a surface boundary condition for air-sea gas exchange. Aside from the boundary carbonate system parameters, the only non-stationary forcing of the model solution comes from the physical boundary conditions and surface forcing (Renault et al., 2021).

In order to ensure the integrity of tracer relationships along isopycnal surfaces, we map the biogeochemical boundary conditions from source data to the model grid using density rather than depth as the vertical coordinate, while retaining the mean-seasonal values of *T* and S (hence density) along the boundary as specified in the physical conditions (Renault et al., 2021). This prevents any errors in the depth of isopycnal surfaces inherited from the physical boundary data (Mercator) from biasing the biogeochemical properties along that surface. Concurrently, this results in biogeochemical boundary conditions responding to interannual variability in isopycnal depth, despite being climatologically fixed along isopycnal surfaces at the boundary. Thus interannual biogeochemical variability is propagated into the model domain because of the covariance between property isopleths and isopycnal displacements rather than changing biogeochemical water mass properties on density surfaces. However, biogeochemical variability along isopycnal surfaces in the interior domain can still arise from varying the proportions (mixture) of water masses entering from different boundaries, or from time variable rates of biogeochemical transformation on isopycnals in the model interior.

The CCS is among the best-sampled regions of the world’s oceans. Hydrographic sampling and biological rate measurements have been conducted repeatedly if not routinely along several sections off the West coast, most notably in the CalCOFI sampling area in the Southern California Bight, off Monterey Bay, California, off Newport, Oregon, and Line P off Victoria, British Columbia at the northern edge of the 4 km model domain. Despite the abundant datasets from this region, data density is still sparse for much of the central California coast and for many biogeochemical properties of interest (*e.g., Fe*). The total number of profiles in the 2013 World Ocean Database (WOD; downloaded from https://www.nodc.noaa.gov/OC5/WOD13/ and including the hydrographic line data) are plotted for 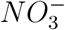 and *O*_2_ over the entire historical data period (1955-2013; Fig. 3).

**Figure 3:**
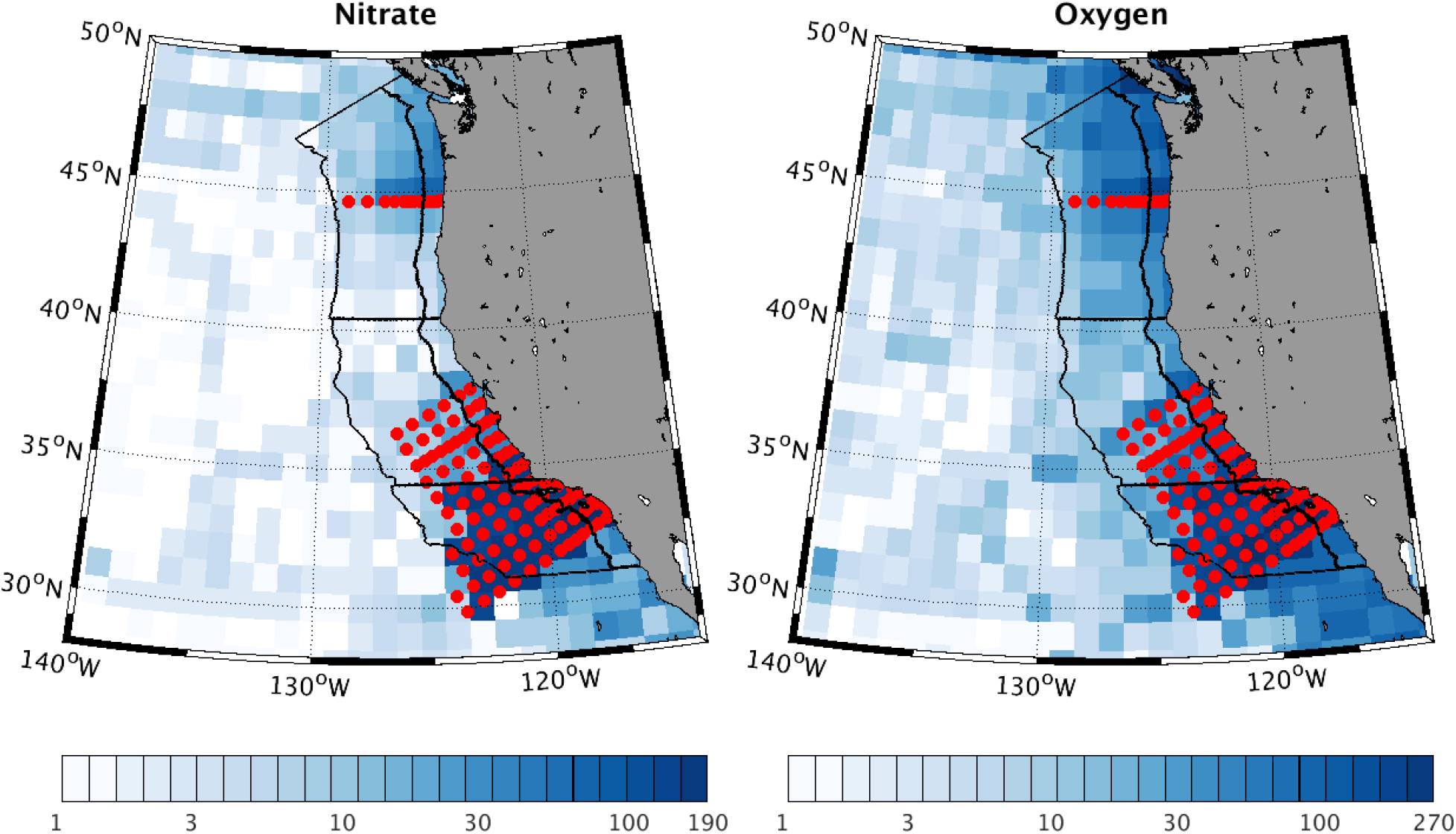
Hydrographic data density used for model validation. Observations in the World Ocean Database are binned in a regular 1° latitude/longitude grid for each month over the entire historical data period (1955-2013). Total number of months with a profile are shown for 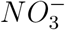 (left) and *O*_2_ (right). Nominal station locations for major repeat hydrographic lines used for validation (see Fig. 16) are shown (red circles), along with the boundaries (black lines) used for regional time series comparisons in Figs. 6,7, 14, and 21.

To facilitate comparison of model outputs to data, and particularly seasonal cycles, we defined 6 regions (Fig. 3), dividing the CCS by distance from the coast into nearshore (0-100 km) and offshore (100-500 km) regions, and by latitude into the Northern, Central and Southern CCS. These designations are somewhat arbitrary, but are based on a combination of topographic delineations and to ensure adequate data coverage in each region. We focus our validation efforts on broad-scale measures that can be evaluated from climatological databases, namely WOD and its objectively-mapped climatological representation, the World Ocean Atlas (WOA; Garcia-Reyes et al. (2014)). We further evaluate the vertical and cross-shelf structure of biogeochemical variables at greater resolution at the hydrographic line locations. Higher frequency biogeochemical measurements from moorings are generally available only for more recent periods, and primarily from nearshore environments. Model comparison to mooring data is left for planned downscaling of these simulations better suited to examining high-frequency variability.

## 3 Results

We describe the spatial patterns and temporal variability of model biogeochemical solutions and their fidelity to observational datasets. Of the numerous properties and rates of the biogeochemical system that are predicted by the model, we focus on those that are most important to ecosystem primary productivity and the overall elemental cycling of carbon, nitrogen, and oxygen, and are best observed over scales captured by the model. This analysis begins with the photic zone, with particular emphasis on the factors driving NPP variability at multiple scales of space and time. We also evaluate the export of this productivity to depth. Second, we present results from the ocean thermocline, where the respiration of exported surface productivity contributes to hypoxic and corrosive conditions. Aspects of the model solution that are not presented include nutrients that are not limiting, rates that lack large-scale and climatological datasets, and variability that is poorly resolved by a *dx* = 4 km model (*e.g.,* submesoscale and nearshore phenomena).

### 3.1 Photic Zone

#### 3.1.1 Chlorophyll and Net Primary Production

We begin with an evaluation of model distributions of Chlorophyll-a (hereafter *Chl*), and Net Primary Productivity (NPP), both of which can be estimated from remote sensing of ocean color. While NPP is of greater biological significance, its estimation is less direct than for *Chl*.

The model *Chl* concentrations are governed by the product of biomass and the *C : Chl* ratios. Biomass is subject to advection and to ecosystem transformations (see Appendix). The *C : Chl* ratio is determined by photoacclimation, or the amounts of light-harvesting pigments and photo-protective compounds produced by phytoplankton in response to their growth environment. This process is included in the ROMS-BEC representation of phytoplankton physiology following the model of Geider et al. (1998), which relates changes in chlorophyll synthesis and nutrient uptake in response to changing PAR. The dominant patterns of *Chl* are also found in biomass (see below), indicating that photoacclimation is not the leading factor in *Chl* variability. While biomass may be a more ecologically meaningful comparison, we validated model solutions using *Chl* because it is more directly estimated from ocean color sensing. The frequency distribution of *Chl* in both ROMS and SeaWIFS remote sensing data is approximately log-normal and is mapped after logarithmic transformation.

The annual mean concentrations of *Chl* vary most strongly in the cross-shore direction, with relatively weak alongshore gradients, a well-known pattern in observations (Banas and Hickey,2008) that is well represented by the model (Fig. 4a,b). The offshore drop-off in *Chl* is somewhat weaker in model simulations, resulting in a wider band of high coastal *Chl*, a tendency that is not reflected in NPP (discussed immediately below). Kessouri et al. (2020) shows that there is some sensitivity in these distributions to model resolution, with higher resolution increasing the nearshore biomass and productivity. The leading pattern of variability in climatological *Chl* is characterized by a seasonal cycle that is also largely synchronous along the coast.

**Figure 4:**
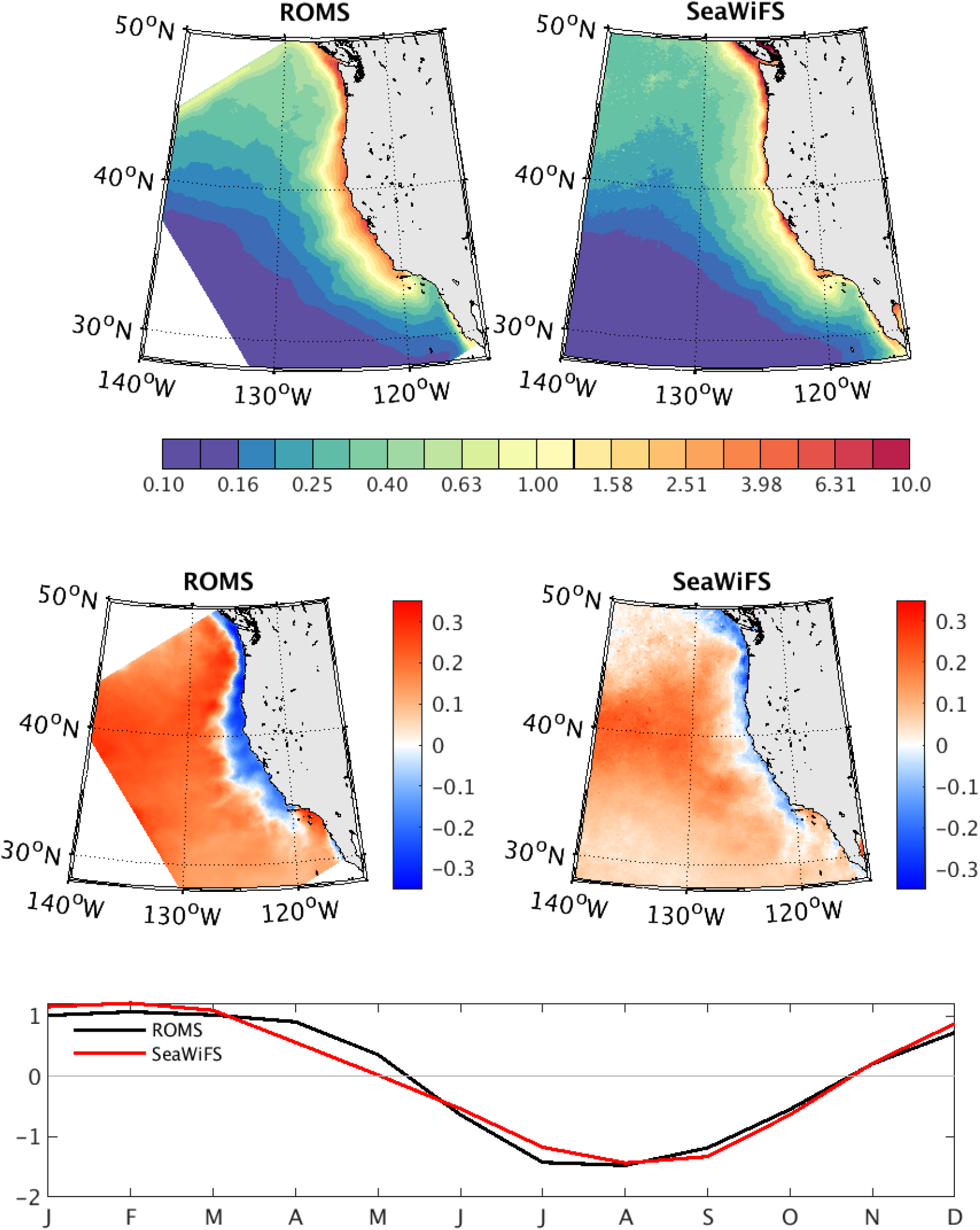
Mean annual Chlorophyll (*Chl*) and its seasonal cycle. Mean annual concentrations (upper panels) are shown in mg m^−3^ averaged over the simulation period in both model output and SeaWiFS remote sensing level 3 product. The seasonal cycle is shown as the spatial loading patterns (middle panels) and time series (bottom panel) of the first EOF of climatological values of log10[*Chl*]. In both the mean annual and seasonal variations of *Chl*, the dominant variations are cross-shore and at Point Conception, which separates the central CCS from the SCB. High coastal *Chl* extends further offshore in model solutions than in observations, a bias that is not found in productivity (Fig. 5).

The leading Empirical Orthogonal Function (EOF) (Fig. 4c,d,e), has *Chl* reaching peak values in late summer, and it accounts for the large majority of the climatological variance in both observations (EOF1=64% variance) and model solutions (EOF1 = 69% variance). The second EOF (not shown) also has a similar loading pattern, but with more meridional structure offshore and a minimum in spring-this accounts for only ≈13% and 19% of the variance in observations and the model, respectively. The seasonality of *Chl* reveals anti-phased cycles between nearshore high-*Chl* band and the lower *Chl* offshore. While near-shore *Chl* peaks in late summer, the offshore surface *Chl* has a minimum. This pattern is not found in the EOFs of depth-integrated NPP, indicating that the dipole structure of the *Chl* pattern results from a vertical redistribution of *Chl* to greater depths in offshore waters as they become more oligotrophic due to nutrient uptake during summer months. This interpretation is confirmed in the analysis of the vertical *Chl* maximum (Sec.3.1.2).

In BEC, NPP depends on the sum of *j* model phytoplankton biomasses (*B_j_*), their maximum growth rates (*μ_max_*(*T*) = *μ*_0_*T*^1.06^), and the limitation of those rates by light (0 ≤ *γ_j_*(*I*) ≤ 1) and the minimum Michaelis-Menten function λ_i,j_(0 ≤ λ_i,j_(*N_i_*) ≤ 1) among the *i* nutrients with half-saturation *K_i,j_* (0 ≤ *λ_i,j_* (*N_i_*) ≤ 1), written as:

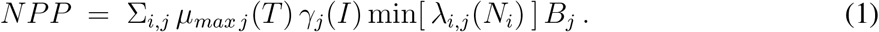

The spatial patterns of modeled NPP fall within the range of satellite-derived estimates (Fig. 5). The two commonly used satellite-based algorithms are the vertically generalized production model (VGPM) (Behrenfeld and Falkowski, 1997) and the carbon-based productivity model (CbPM) (Westberry et al., 2008). The VGPM estimates productivity on the basis of light and chlorophyll concentrations, calibrated to a predominantly coastal radiocarbon incubation dataset. The CbPM additionally incorporates phytoplankton backscattering and growth rate relationships in order to estimate productivity as a function of phytoplankton biomass, and it is calibrated to subtropical gyre radiocarbon incubations. The two algorithms exhibit a relatively wide range for the CCS region, reflecting the considerable uncertainty in “empirical” NPP estimates as well as differences in the measurements underlying each. The VGPM algorithm has a larger offshore gradient, with higher coastal values, and lower values in the open ocean, compared to the carbon-based CbPM. NPP rates from ROMS-BEC fall between the two remote sensing products, but are generally closer to the values of the VGPM algorithm, supporting higher near-shore rates, lower offshore rates, and increased seasonality relative to the CbPM. The VGPM has been explicitly calibrated against radiocarbon bottle incubations from the CalCOFI program, and it is therefore likely to be more accurate in this region (Kahru et al., 2009). Indeed, we find that ROMS-BEC rates and distributions of productivity are also consistent with direct estimates from ship-based data both from CalCOFI (Fig. 5; Munro et al. (2013)) and the broader subtropical Northeast Pacific (Palevsky et al., 2016).

**Figure 5:**
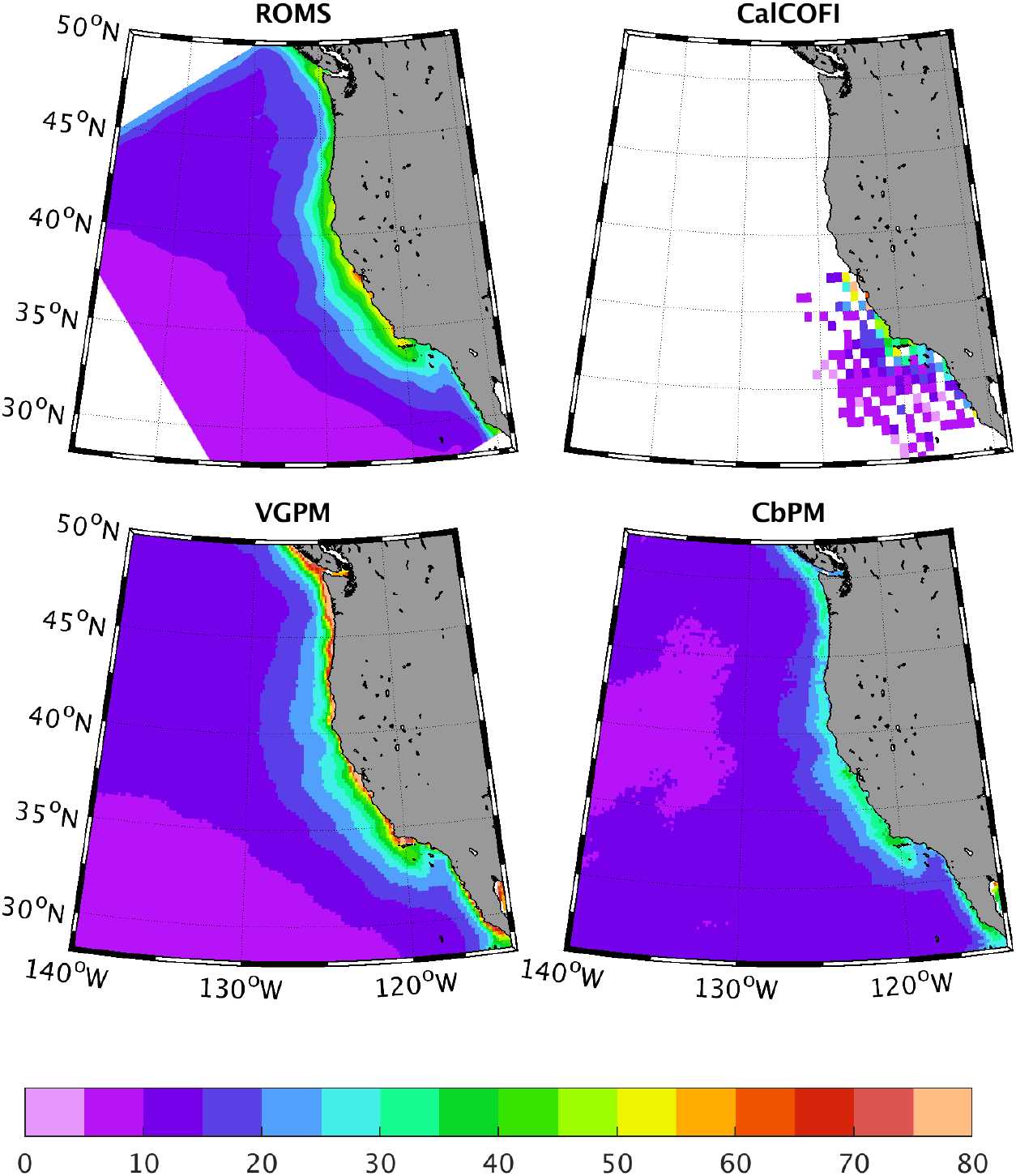
Spatial distribution of annual Net Primary Productivity (mol *C* m^−2^ yr^−1^) integrated over the depth of the photic zone, from (top left) ROMS, (top right) CalCOFI, (bottom left) VGPM, and (bottom right) CbPM.

Productivity in the northern CCS has been consistently biased in regional models (Banas and Hickey, 2008), including in our initial simulations. We conducted simulations with and without lateral nitrogen fluxes at the Strait of Juan de Fuca imposed as boundary condition from model simulations by Davis et al. (2014). Consistent with that study, without the nutrient inputs from the Salish Sea, NPP was biased low by > 50%. The inclusion of the effects of nitrogen inputs at the Strait of Juan de Fuca brought the model much closer to satellite-based empirical models. The inclusion of these inputs is consistent with the study by Davis et al. (2014), and it is used in all results reported here.

The seasonal cycle of NPP is also well captured by the model (Fig. 6). In all 6 regions of the CCS, the climatological NPP, integrated over the depth of the photic zone and averaged over the regional mask, exhibits an amplitude and phasing that is within the range of satellite-based empirical models. The most notable exception is the offshore Northern domain, where a spring bloom is predicted to be of stronger magnitude than estimated by either satellite product. This model result is consistent with measured geochemical tracers, which also indicate the spring bloom in the offshore Northeast Pacific is greater than estimated from the satellite algorithms (Palevsky et al., 2016). A smaller discrepancy occurs in the southern nearshore region, where ROMS-BEC generates greater summer production than either of the satellite algorithms. Model NPP in the oligotrophic part of the domain is lower than satellite estimates, but is more consistent with the most offshore values in the depth-integrated rates based on radiocarbon bottle incubations from CalCOFI. Overall, where and when ROMS-BEC and satellite algorithms for NPP disagree, ROMS-BEC output is generally closer to the available observational data.

**Figure 6:**
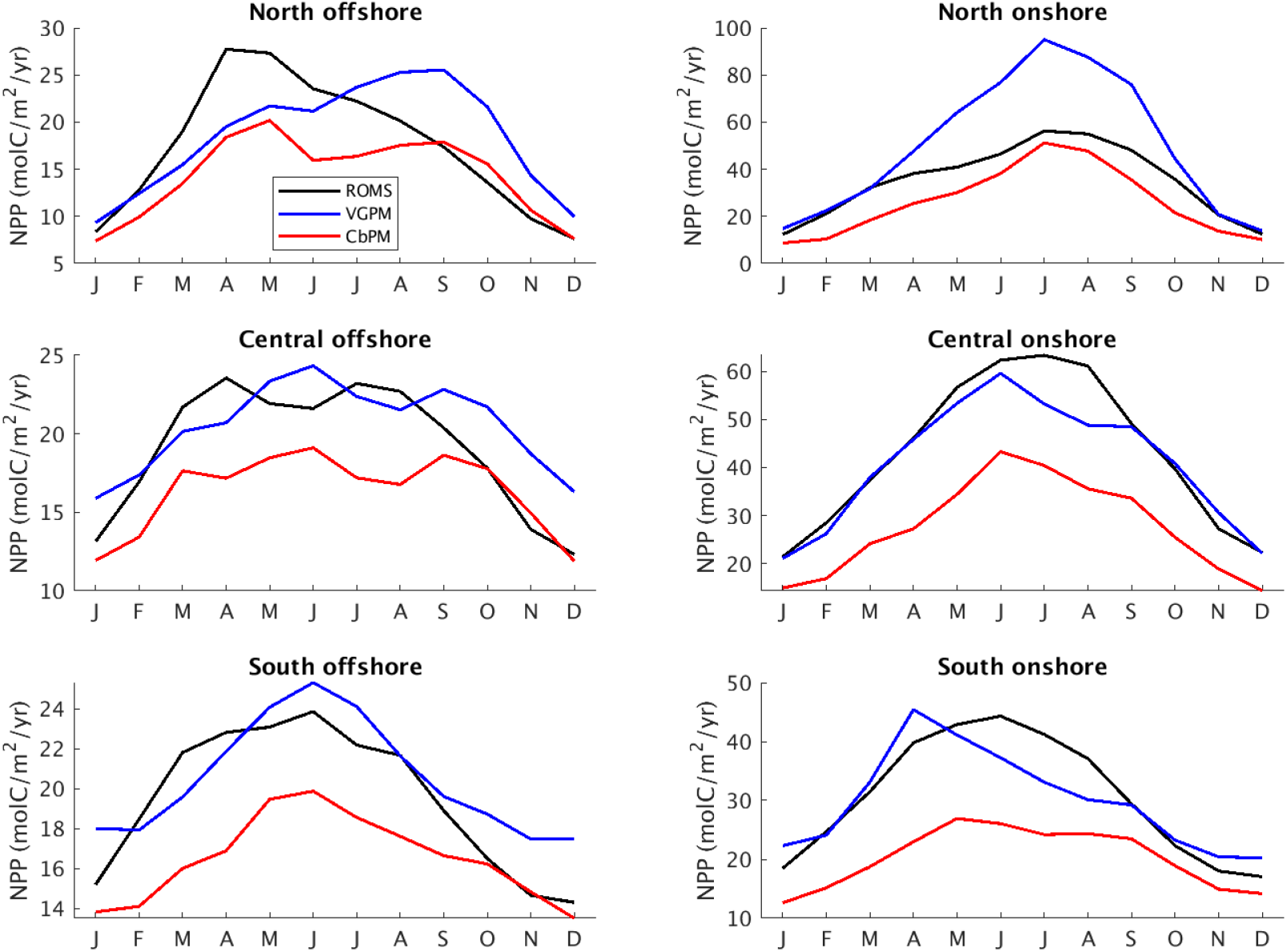
Seasonal cycle of annual Net Primary Productivity. The NPP rate (mol *C* m^−2^ yr^−1^) from ROMS-BEC (black), and two satellite algorithms (VGPM, blue; CbPM, red) are integrated over the depth of the photic zone, and averaged over 6 regions (see Fig. 3) from northern (top row), central (middle row), and southern (bottom row) of the CCS, and divided by distance from the coast into a nearshore (0-100 km; right column) and offshore (100-500 km; left column) region.

#### 3.1.2 Seasonal limitation of productivity by light and nutrients

To evaluate the role of environmental factors shaping the seasonal cycle and regional differences in rates of productivity, we computed monthly mean limitation factors for each of the environmental variables that modulate the maximum growth rates, including macronutrients 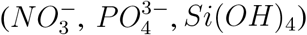, *Fe*, and light; see (1). By construction, growth rates are limited by only one nutrient at a time (Liebig’s Law of the Minimum), such that only the lowest value has an influence on rates. Light operates as a multiplicative factor on nutrient limitation, reducing growth relative to the light-saturated photosynthesis rate (Geider et al., 1998). Temperature influences maximum growth rate, but is not considered a limiting factor in the upper ocean, so is not analyzed here.

Over a climatological seasonal cycle, small plankton growth rates are almost always reduced by light more so than by nutrients, regardless of season or location (Fig. 7). The small plankton are assumed to have a lower half-saturation constant for nutrients, and the resulting higher affinity makes them less prone to nutrient limitation than large plankton are. Thus, in the inshore regions where nutrients are high, light is always limiting. Offshore, nutrients can limit small phytoplank-ton to a similar degree as light in the summer. However, large phytoplankton make up ≈ 90% of modeled NPP on average, and large phytoplankton (“diatoms”) are primarily limited by nutrient availability. In the model, this results from a nutrient supply by combined wind-induced upwelling and eddy-induced subduction (Gruber et al., 2011; Nagai et al., 2015; Renault et al., 2016a) that is unable to saturate the potential net uptake of nitrogen by phytoplankton at prevailing light levels. Significant eddy fluxes also occur due to submesoscale eddies and fronts in the CCS (Kessouri et al., 2020), but are not resolved in this model. Large phytoplankton only experience light limitation in the northern CCS, where the seasonal cycle alternates between winter light limitation and nutrient limitation for the rest of the year.

**Figure 7:**
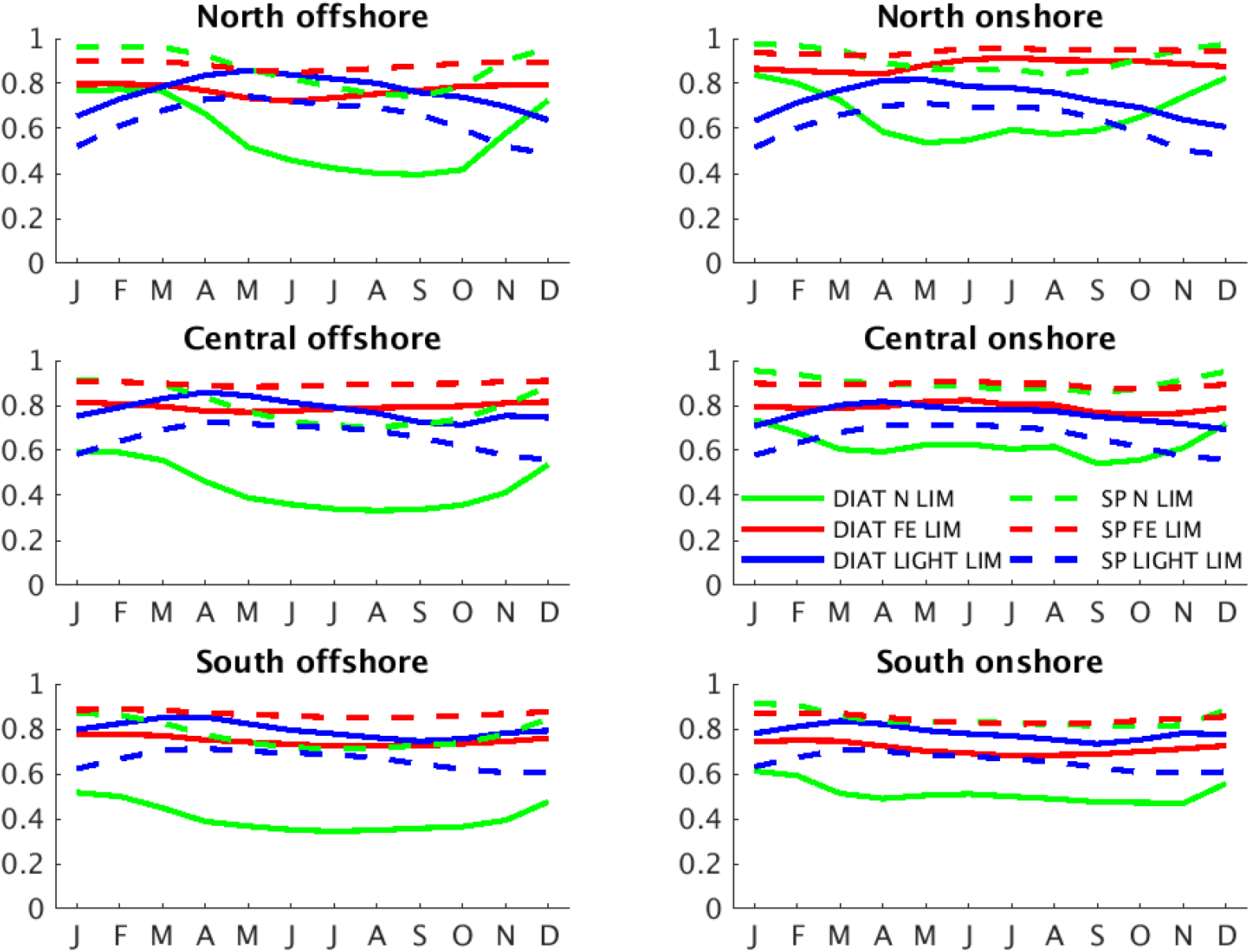
Seasonal cycle of growth limitation factors for light (γ term in (1), blue) and for nutrients (λ term in (1): 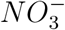 (green) and *Fe* (red) for diatoms (solid) and small phytoplankton (dashed)). Factors are NPP-weighted and averaged over the photic zone for each region shown in Fig. 3. Limitation factors close to 1 mean no limitation; values close to 0 mean complete limitation.

While on the regional scales used for this analysis nitrogen limitation appears more stringent than *Fe* limitation throughout the CCS, significant *Fe* limitation occurs on smaller scales and shorter durations (see below). The seasonal amplitude of plankton growth rates is relatively small (≈ 10%), indicating that the amplitude of seasonal production (≈ 100%) is governed by seasonal controls on biomass rather than growth rates. The trade-off between light and nutrient limitation spatially and seasonally is a ubiquitous feature of phytoplankton distributions and phenology. Moving deeper in the water column, light becomes more limiting as photosynthetically active radiation (PAR, light of 400 - 700 nm wavelengths) is absorbed and scattered, while nutrient concentrations are greatest below the surface mixed layer and rapidly decrease upward across the seasonally variable pycnocline; *i.e.,* there is a well-defined nutricline.

The competing influences of nutrient and light limitation on depth of optimal plankton growth (i.e., highest growth rates) are reflected in the depth of the deep chlorophyll maximum (DCM). In the CCS, the depth of the DCM deepens from the coast to the open ocean, suggesting that the growth-maximizing combination of light and nutrients is found deeper offshore, consistent with a deepening nutricline that intensifies nutrient limitation at the surface and light limitation where nutrients are abundant, for both small and large phytoplankton. We use the observed pattern of the DCM depth as an indicator of whether the model achieves a realistic trade-off between these two countervailing growth condition gradients (Fig. 8).

**Figure 8:**
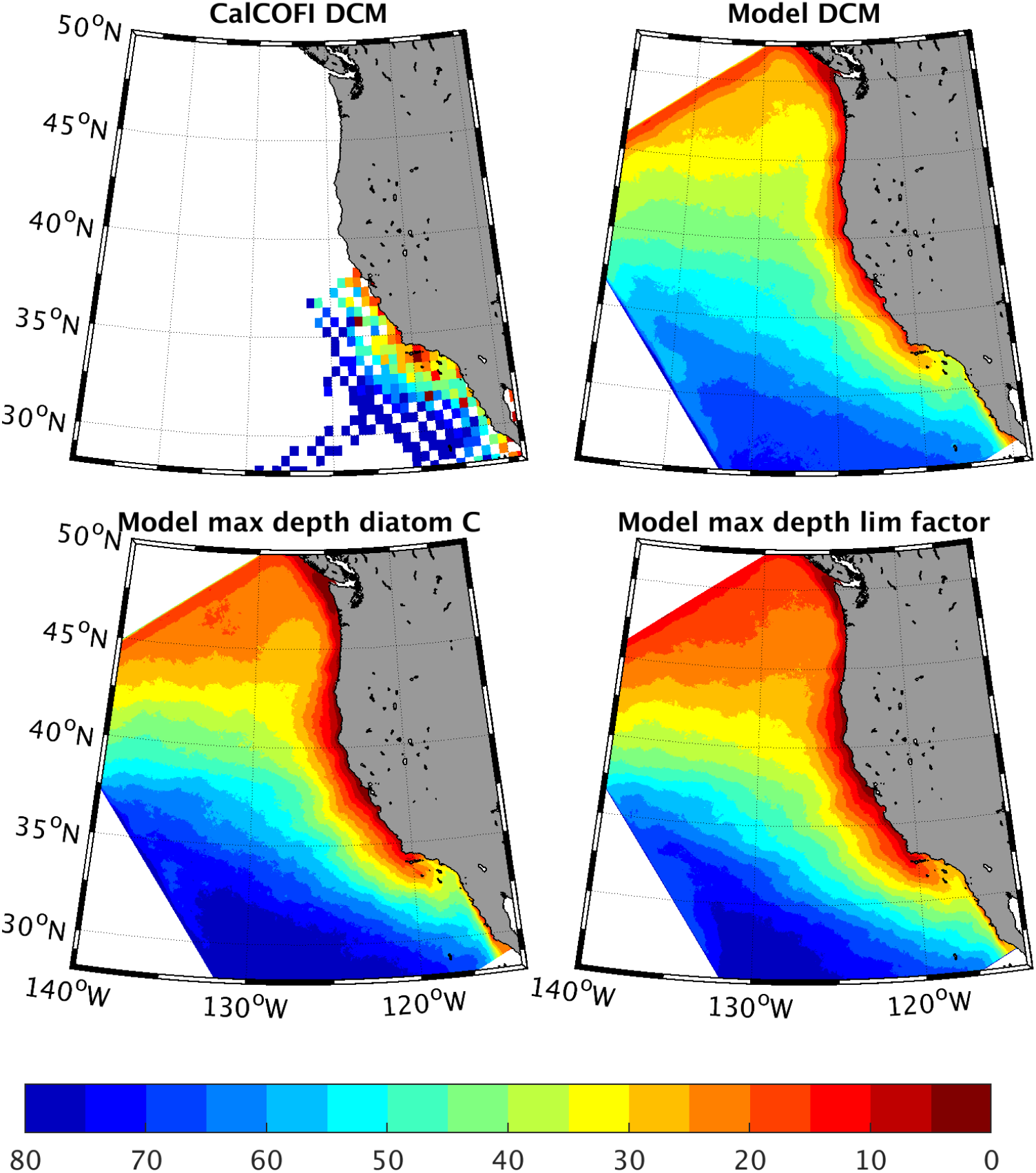
Depth of the vertical maximum of Chlorophyll in CalCOFI (top left) and ROMS (top right), and of model diatom biomass (bottom left) and nutrient limitation factor (bottom right). The correspondence between model fields demonstrates that the model DCM tracks nutrient limitation, while the fidelity to observations implies the model captures a realistic trade-off between nutrient and light limitation.

However, for DCM depth to be a reliable indicator of phytoplankton growth rate trade-offs, we must rule out two alternative interpretations relating to isopycnal advection and photoacclimation. First, the offshore deepening of the DCM can have a physical origin. Because it closely follows the plunging of isopycnal surfaces offshore, the vertical peak in biomass and associated chlorophyll could be caused by eddy subduction carrying high surface chlorophyll away from the coast along deepening isopycnals (Gruber et al., 2011; Nagai et al., 2015; Renault et al., 2016a). To evaluate this possibility, we compared the depths of maximum chlorophyll concentration and phytoplankton biomass to the depth at which the product of light and nutrient limitation factors are maximized (Fig. 8). These maps are virtually indistinguishable, suggesting the DCM follows growth rates rather than advection. As a more stringent test, we performed a short (5-year) simulation in which surface PAR was reduced by 10%. The results revealed a significant shoaling of both the biomass and *Chl* peaks, but no detectable change in isopycnal depths, confirming that these depths do in fact reflect a nutrient-light trade-off rather than advection along density surfaces. Second, the peak depth of *Chl* may also be decoupled from that of biomass and growth rates due to photoacclimation, or a shift in the amounts of light-harvesting pigments and photoprotective compounds produced by phytoplankton in response to the light-environment. This process is included in the ROMS-BEC representation of phytoplankton physiology following the model of Geider et al. (1998), which relates changes in chlorophyll synthesis and nutrient uptake in response to changing PAR. Indeed, we find that the depth of the DCM is slightly deeper than that of the maximum plankton biomass. However, the offshore and latitude gradients of the depth of peak biomass and chlorophyll are very similar. DCM deepening offshore is consistent with optimized growth conditions in the model, and reproduces the pattern observed in the available CalCOFI data (Fig. 8).

In summary, the Chl maximum does not primarily reflect photoacclimation or isopycnal transport, although it is affected by those processes. Instead, it is found at approximately the same depth as that of peak biomass (panel c), which in turn is found at the depth that maximizes growth rate through the combined impacts of nutrients and light (panel d). This is the same depth at which NPP, the product of biomass and growth rate, is also maximum. Thus, these correspondences indicate that the model DCM is a reflection of the essential trade-off between light and nutrient limitation, and the fidelity to the observed DCM implies this trade-off is adequately captured.

Growth rates are modulated by a complex and evolving pattern of nutrient limitation by reactive nitrogen 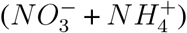 and soluble dissolved *Fe*, with no appreciable limitation by *Si*(*OH*)_4_ and 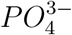 in the CCS. The limitation factors are mapped as a biomass-weighted fraction of time that each of the nutrients is most limiting (Fig. 9). The spatial pattern among nutrients largely reflects the areas where 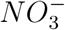 supply routinely exceeds maximum potential uptake seasonally. Thus, the waters entering the CCS from the subarctic High Nutrient - Low Chlorophyl (HNLC) region are most frequently *Fe* limited. Along the coast, the seasonal upwelling of excess nitrogen leads to significant periods of *Fe* limitation as well. In the coastal zone off Monterey Bay, *Fe* limitation has been diagnosed via incubation experiments, in a band of water slightly offshore, with nitrogen limitation both in more shoreward and open coastal zones (Firme et al., 2003). This pattern is consistent with that predicted by the model (Fig. 9b inset). Most of the rest of the model domain is perpetually nitrogen limited.

**Figure 9:**
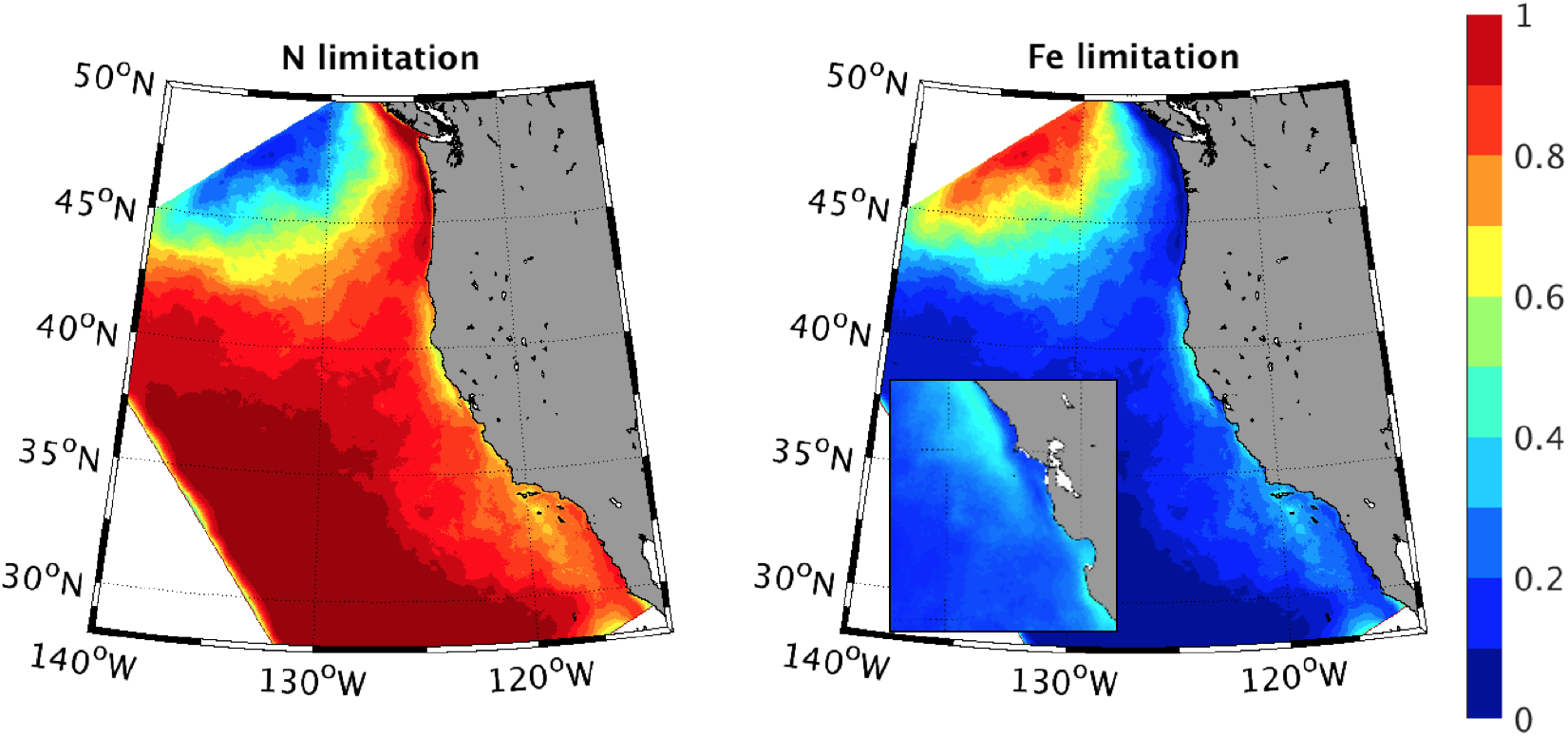
Frequency of limitation by 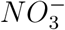 (left) and *Fe* (right) for the model’s dominant primary producer, diatoms. The limitation factors are weighted by biomass (as in Fig. 7), using 5-day output, and plotted as the fraction of time that each nutrient limitation factor is the lowest among nutrients. The inset shows offshore band of relatively frequent *Fe* limitation along the central CCS, similar to that observed by (Firme et al., 2003).

#### 3.1.3 Nutrient concentrations

Surface nutrient concentrations provide an important measurable test of system behavior. For nutrients that limit phytoplankton growth, accurately simulating their distributions is a necessary condition for a mechanistic prediction of NPP. Moreover, they provide an integrative measure of net community production (equal to NPP less community heterotrophic respiration) and export of organic matter to the thermocline. We therefore compared model predicted distributions of the two primary limiting nutrients, 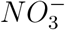 and *Fe*, to available observations.

Coastal measurements of dissolved *Fe* reveal a spatially patchy distribution, reflecting its short residence time with respect to removal by plankton uptake and particle-active scavenging. Existing data are too sparse to yield a clear climatological pattern for model validation. However, the primary coastal region where the model predicts most frequent *Fe* limitation, in the central CCS, has been relatively well sampled, including on two cruises off Monterey Bay that also tested for *Fe* limitation (Firme et al. (2003); see Fig. 9, inset). Given the lack of a clear large-scale pattern of surface *Fe* levels, we used a more statistically-based validation metric, focusing on the relative frequency of *Fe* measurements versus concentration and distance from the shore (Fig. 10). On average, the data and model both show a decline in the mean and median *Fe* levels with offshore distance. This reduction is driven largely by the decreasing frequency of high concentrations, while the most commonly observed *Fe* levels remain consistent at ≈ 0.5 · 10^−6^ mol m^−3^ regardless of distance from shore. The thinning tail of high concentrations in *Fe* distribution with cross-shelf distance occurs in both modeled and observed fields, but is more pronounced in the measurements.

**Figure 10:**
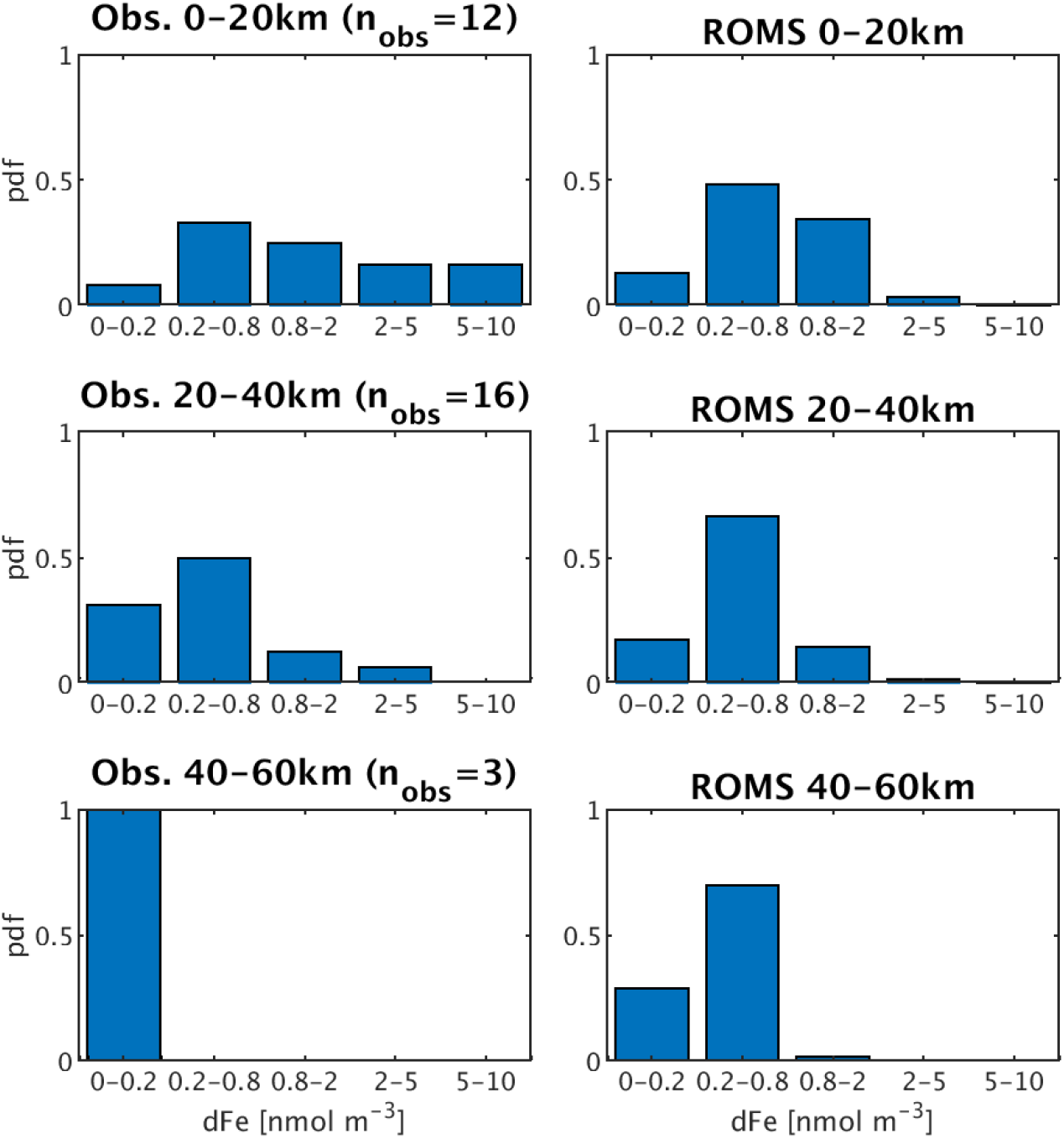
Histogram of surface *Fe* concentrations (10^−6^ mol m^−3^) from observations (left) and ROMS (right). Because *Fe* concentrations are patchy in nature and sparsely observed, values are binned by distance from shore (0-20 km, upper row; 20-40 km middle row; 40-60 km lower row) to reveal a cross-shelf gradient, in the same latitude band off Monterey Bay for which observations were made on summer cruises (Firme et al., 2003).

In the CCS, the nutrient most often limiting NPP is reactive nitrogen, of which by far the largest and most commonly measured pool is 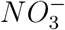. We therefore compare the model simulated patterns of 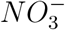 to climatological values from the World Ocean Database (Fig. 11). A depth of 50 m is chosen because it approximates the average depth of maximum biomass and NPP (see Fig. 8) and is generally near the base of the photic zone on the continental shelf. We included all historical measurements for this analysis because the data density in the model period (1994-2010) was much sparser, and no significant differences were found between the average 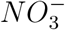 in this period relative to 1955-2013.

**Figure 11:**
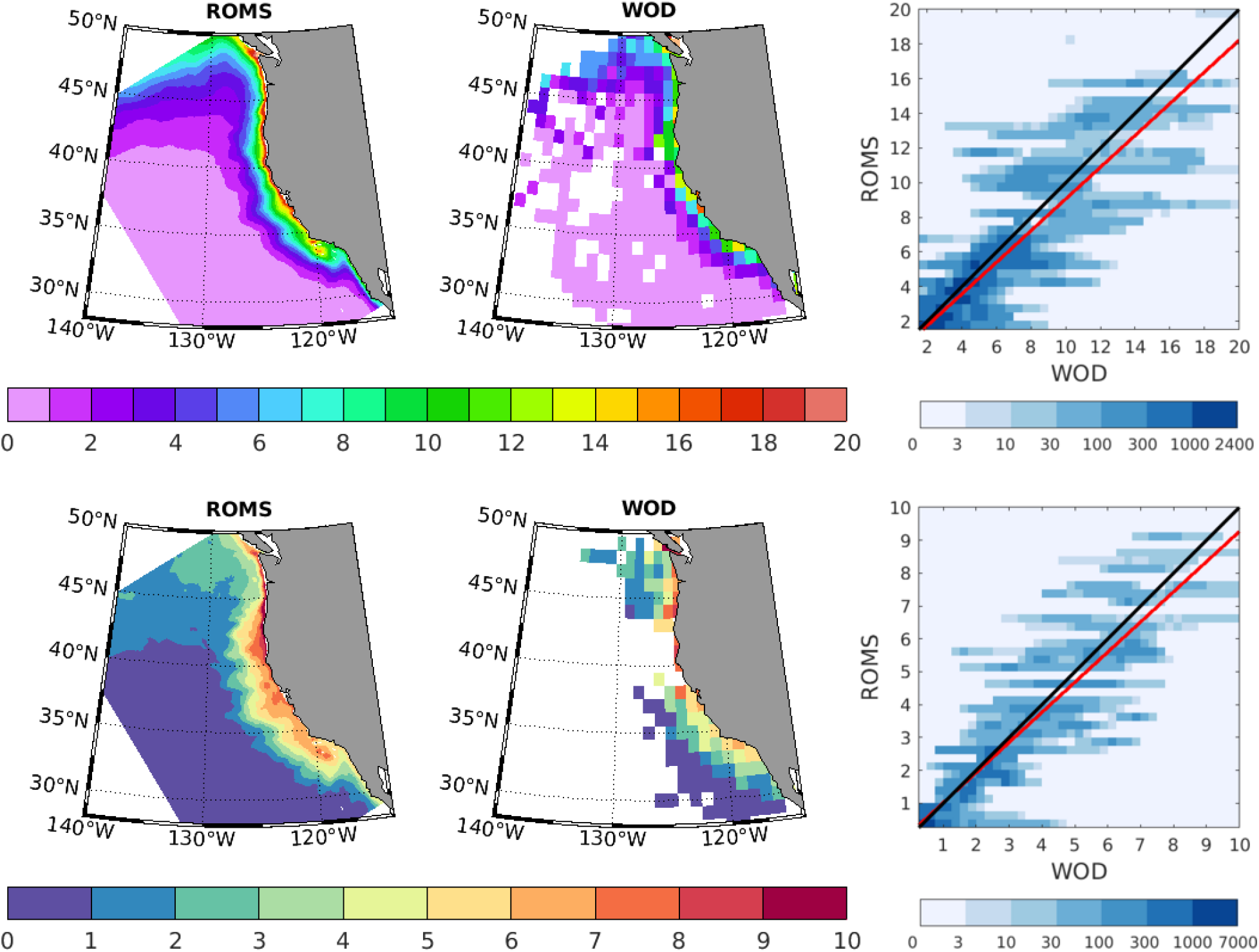
Long-term mean (upper row) and historical variability (bottom row) of nitrate (mmol m^−3^) near the base of the photic zone (50 m) in ROMS (left column), WOD (center column), and their correlations (right column). Variability is mapped as the standard deviation and consists of roughly equal contributions from seasonal and interannual variability (see text). The full period of the World Ocean Database (1955-2013) is used to yield the most robust estimate of variance. The correlations between ROMS and WOD are highly significant (p≪0.01) for both mean and variability, with squared Pearson correlation coefficients (R^2^) of 0.84 and 0.80, respectively.

ROMS-BEC captures regional patterns well for 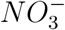 (Fig. 11). Annual mean concentrations of ≈ 15 mmol m^−3^ along the coast decline to values below the half-saturation level for model diatom growth 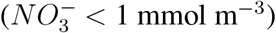 within 500 km from shore. The offshore gradient is similar throughout most of the CCS, except in the Southern California Bight (SCB), where coastal surface values are much lower. Similar model fidelity was found for other macronutrients 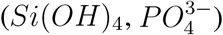, but not shown because they do not reach limiting concentrations. The coastal zone exhibits strong variability in 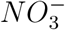 at 50 m, with standard deviations of 5-10 mmol m^−3^ throughout most of the coastal zone, but with a slight northward increase in variance (Fig. 11c). The variability of 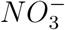 in the climatology (WOA) exhibits a similar spatial pattern, but with ≈ 50% of the magnitude. Thus, approximately half of the variation in surface nutrients most commonly limiting NPP is associated not with the seasonal cycle, but with interannual variability. The model also reproduces observed magnitudes and patterns of 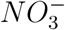 variability (Fig. 11d). We use the interannual anomaly fields in the model and in observations to test the importance of nutrient supply as a mechanism driving changes in NPP over time.

#### 3.1.4 Interannual variability in surface nutrients and NPP

Factors that limit NPP during the mean seasonal cycle may drive interannual and longer term productivity changes. We examined correlations between NPP and both light and nutrient concentrations in model simulations and observations, where available. Interannual anomalies in 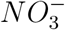 in ROMS-BEC are found to be well correlated (R^2^≈ 0.5) to the density of water at 100 m (Fig. 12). This reflects the role of pycnocline heave and vertical mixing in supplying macronutrients, and alleviating local nutrient limitation. Observations show a similar magnitude of correlation in the southern and central CCS, but a weaker correlation to the north. This may reflect the role of nutrient supply processes that are either missing or represented only climatologically in our model, and not connected to pycnocline heave. Because the weaker correlations are in the northern domain where nutrients can enter from subarctic surface waters, the climatological 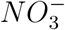 used for the boundary conditions is a likely culprit. However, the presence of river 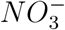 sources or a variable Juan de Fuca flux could also weaken the correlation in the observations relative to the model.

**Figure 12:**
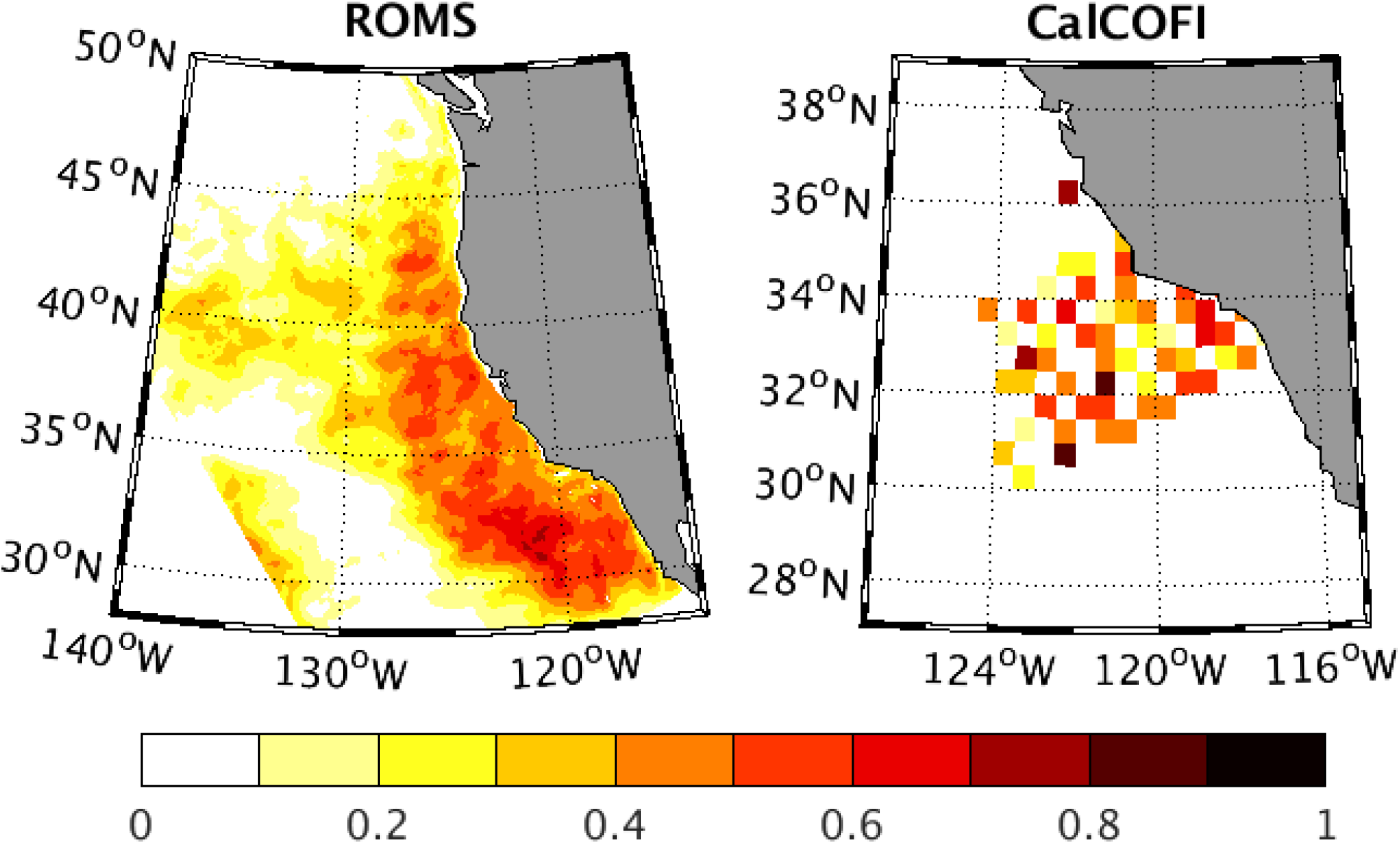
Correlation (*R*^2^) between NPP and density, from ROMS-BEC (left) and CalCOFI measurements. The NPP rate is integrated over depth from ^14^C bottle incubation data, and density is interpolated to 100 m as an index of nutrient supply (see Fig. 13). In both variables, the mean seasonal cycle is removed leaving interannual variations. Relationships have similar strength in data and model, and indicate that ≈ 50% of interannual variability in NPP can be attributed to anomalous nutrient supply due to pycnocline heaving.

Predicted correlations between NPP and density can be tested directly by combining bottle incubations and hydrographic observations in the southern CCS (Fig. 13). Relationships between nutrient and density anomalies (subtracting the mean seasonal cycle), are of similar strength, accounting for ≈ 50% of the variability in both CalCOFI observations and model simulations. Interannual anomalies in NPP in ROMS-BEC are also significantly correlated with surface PAR (not shown) due to variable cloudiness, though it accounts for a smaller fraction of the variance (≈ 20%). The role of light is confined closer to the coastal upwelling where surface 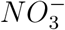 is high, and light availability thus limits phytoplankton growth.

**Figure 13:**
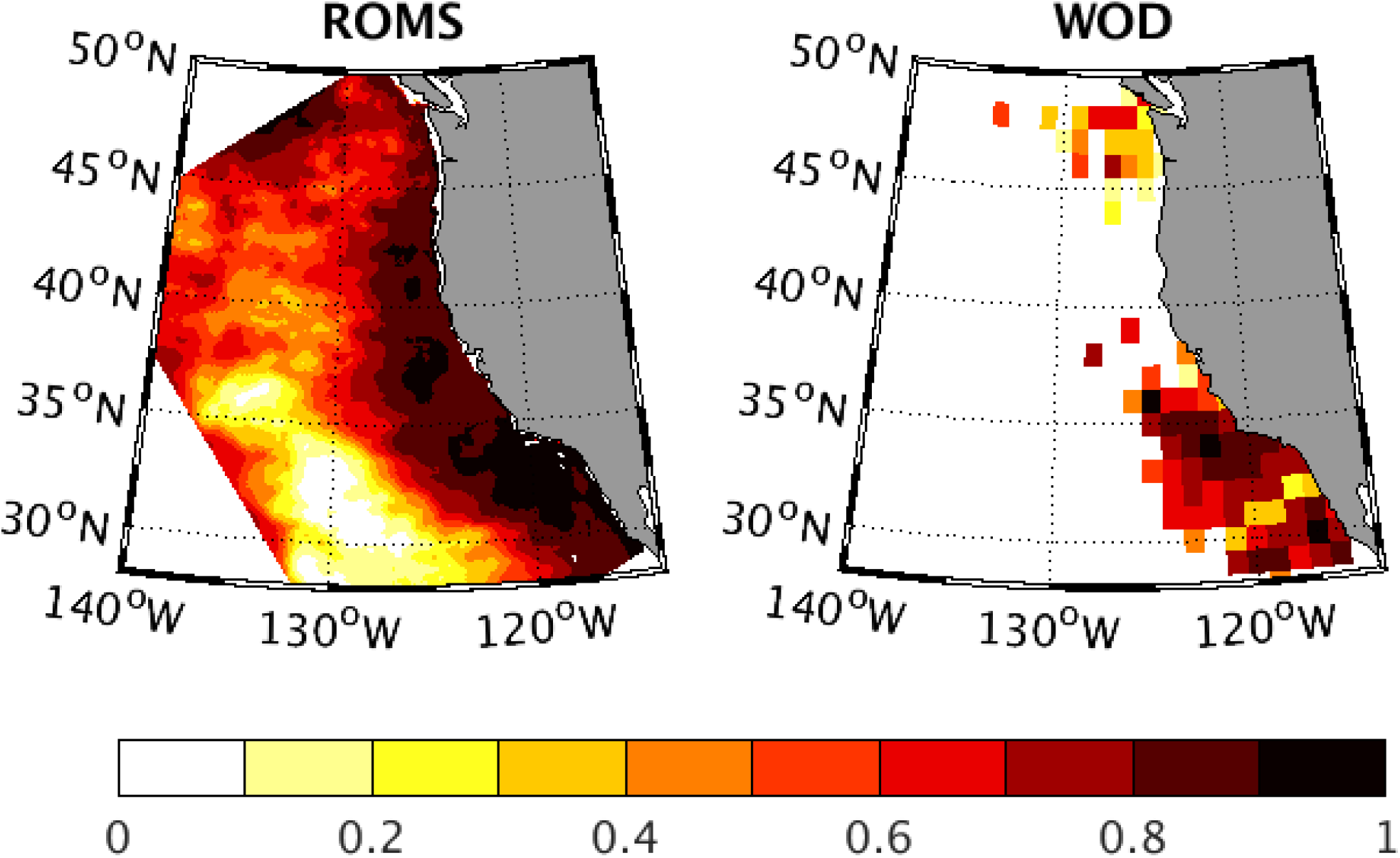
Correlation (*R*^2^) between 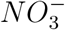 and density at 100 m depth in ROMS-BEC (left) and WOD (right). For both variables, the mean seasonal cycle is removed leaving interannual variations. Interannual variations in subsurface (100 m) 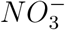 highly correlated with density (*R*^2^ ≥ 0.8) in most of CCS. Correlation is weaker in northern CCS in data than in model.

#### 3.1.5 Carbon fluxes from the photic zone

Of the net production of organic matter by phytoplankton, a substantial fraction can be respired by zooplankton and higher trophic levels. The remainder is available to be transported away, by particle sinking and transport of dissolved organic matter, e.g., via eddy subduction (Sec.3.1.2). The fraction of NPP that is regenerated within the surface ocean depends on food web processes, such as grazing rates. Although data is not available to evaluate large-scale patterns of grazing rates, an indirect comparison can be made through the export flux and the fraction of NPP that is exported in sinking particles rather than recycled (the so-called pe-ratio (Dunne et al., 2005; Murray et al., 1996)).

The fraction of NPP that is exported varies from 5-25%, consistent with the range of values inferred in field studies (Fig. 14). The model predicts highest pe-ratios in the coastal zone, where productivity is high and sea surface temperature relatively low. These dependencies are also consistent with those inferred from a global analysis of rate measurements for NPP and net community production (NCP, assumed equal to total export of both particulate and dissolved organic carbon) (Dunne et al., 2005). For a more quantitative comparison, we compared modeled pe-ratios to those predicted by a statistical model fit to global observations by Dunne et al. (2005), over the seasonal cycle in each of our 6 standard CCS regions (Fig. 14). In both the empirical model and in ROMS-BEC, the mean value, phasing, and seasonal amplitude of changes in pe-ratio are similar. Empirically based estimates generate a similar result that remains consistent with model simulations, even when sea surface temperature is held constant. This suggests that temperature and its impacts on the relative growth rates of phytoplankton and their grazers are not the essential cause of variable pe-ratios.

**Figure 14:**
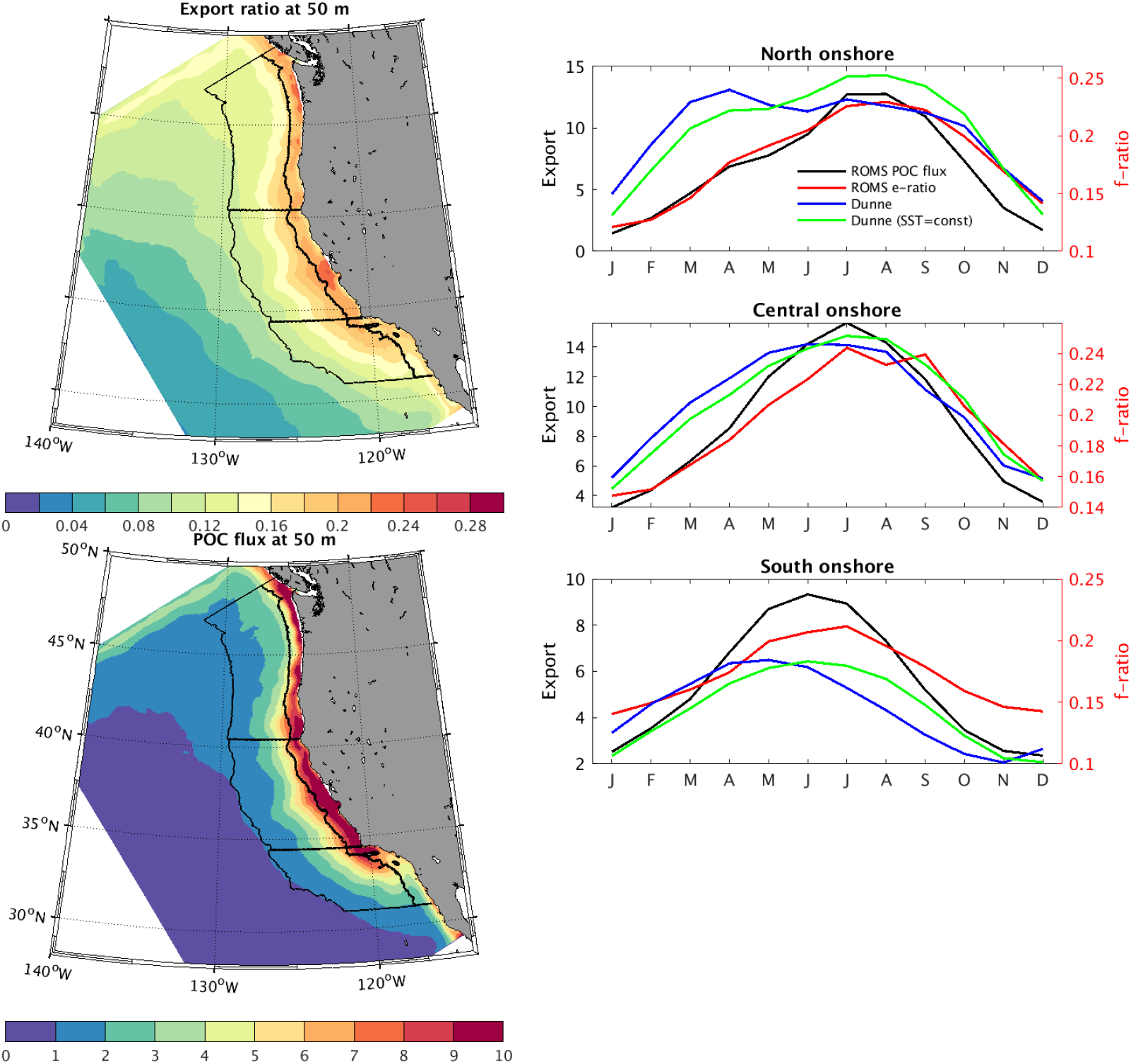
Annual mean and seasonal cycle of fraction of NPP that is exported in sinking particles (*i.e.,* pe-ratio; colored lines), and the export flux (mol *C* m^−2^ yr^−1^; black line). Values diagnosed from model simulations (red line) are compared to an empirically derived algorithm (Dunne et al.,2005) based on *Chl* and SST (blue line), and from the same algorithm applied with constant SST (green line). Observed annual net community production, which should approximate export on an annual basis are measured to 5-10 mol *C* m^−2^ yr^−1^(Munro et al., 2013).

Annual mean export flux represents the transfer of biogenic material from the surface to depth, and thus the influence of CCS productivity on air-sea *CO*_2_ flux and thermocline properties. Model simulated export production ranges from ≈ 1 – 10 mol *C* m^−2^ y^−1^(Fig. 14b). Few measurements of export or net community production are available to evaluate the overall pattern of this flux. In the SCB region, measurements of *O*_2_ have been used to estimate NCP rates of 3 - 17 (mean 6.5) mol *C* m^−2^ y^−1^(Munro et al., 2013), in line with modeled rates and with similar spatial patterns of export (greatest along the Northern coast of the SCB). Munro et al. (2013) also combine information from ^15^N uptake experiments and nitrate based new production ratios (Dugdale et al.,1992; Eppley et al., 1992) with radiocarbon incubations to generate very similar estimates of the magnitude and spatial variability of production in the SCB. Particle-based estimates of export (neglecting the role of dissolved carbon) are lower than observed NCP as well as ROMS-BEC export, 2 - 4 mol *C* m^−2^ y^−1^(Collins et al., 2011; Stukel et al., 2011), though enhanced particle export associated with mesoscale fronts (2 - 3 times greater rates over smaller spatial and temporal scales (Stukel et al., 2017)) highlights the potentially important role of subduction by eddies and fronts in explaining mismatches between observations. Importantly, ROMS-BEC generates such mesoscale features which contribute to model export estimates. While the model assumes particulate matter is redistributed vertically without being transported by the lateral circulation, the magnitudes and pattern of export are not substantially different from models that include explicit 3-dimensional particle transport (Frischknecht et al., 2018).

Model export production is also similar to regional nutrient budget analyses which suggest NCP averaging 7 - 9 mol *C* m^−2^ y^−1^ over the broader CalCOFI region (Roemmich and McCallister,1989; Bograd and Lynn, 2001). Similar nutrient budget analysis indicates that annual NCP should be ≈ 17 mol *C* m^−2^ y^−1^ off Monterrey Bay, again in line with ROMS-BEC estimates of export for that region (Fig. 14b). As noted above, the fidelity of modeled nitrate distributions across the model domain provides a critical broad-scale measure of net biological drawdown, and thus of net community and export production.

Surface ocean *CO*_2_ partial pressure and sea-to-air flux (Fig. 15) is reduced by net community production, increased by surface heat flux, reduced by freshwater fluxes, and modulated by upwelling and lateral circulation. It thus represents another important metric of overall system function. The role of the CCS and its sub-regions in the atmosphere-ocean balance of *CO*_2_ has previously been investigated in several studies, both empirically (e.g.,Hales et al. (2012)) and in models (e.g.,Fiechter et al. (2014); Turi et al. (2014) and references therein). We evaluated the patterns of annual mean and summertime surface *pCO*_2_ in the model hindcast simulation against observations in the SOCATv6 database from 1995-2010 (Bakker and coauthors, 2016), and the associated air-sea fluxes over the modeled coastal region. Similar to observations over the simulated period, strong *CO*_2_ supersaturations are simulated in a narrow band of coastal water within about 100 km of the shore (Fig. 15). The values are highest along the central coast (35N-43N), lowest in the northern CCS, and intermediate in the southern domain. The highest values are associated with major topographic features, as previously noted by Fiechter et al. (2014).

**Figure 15:**
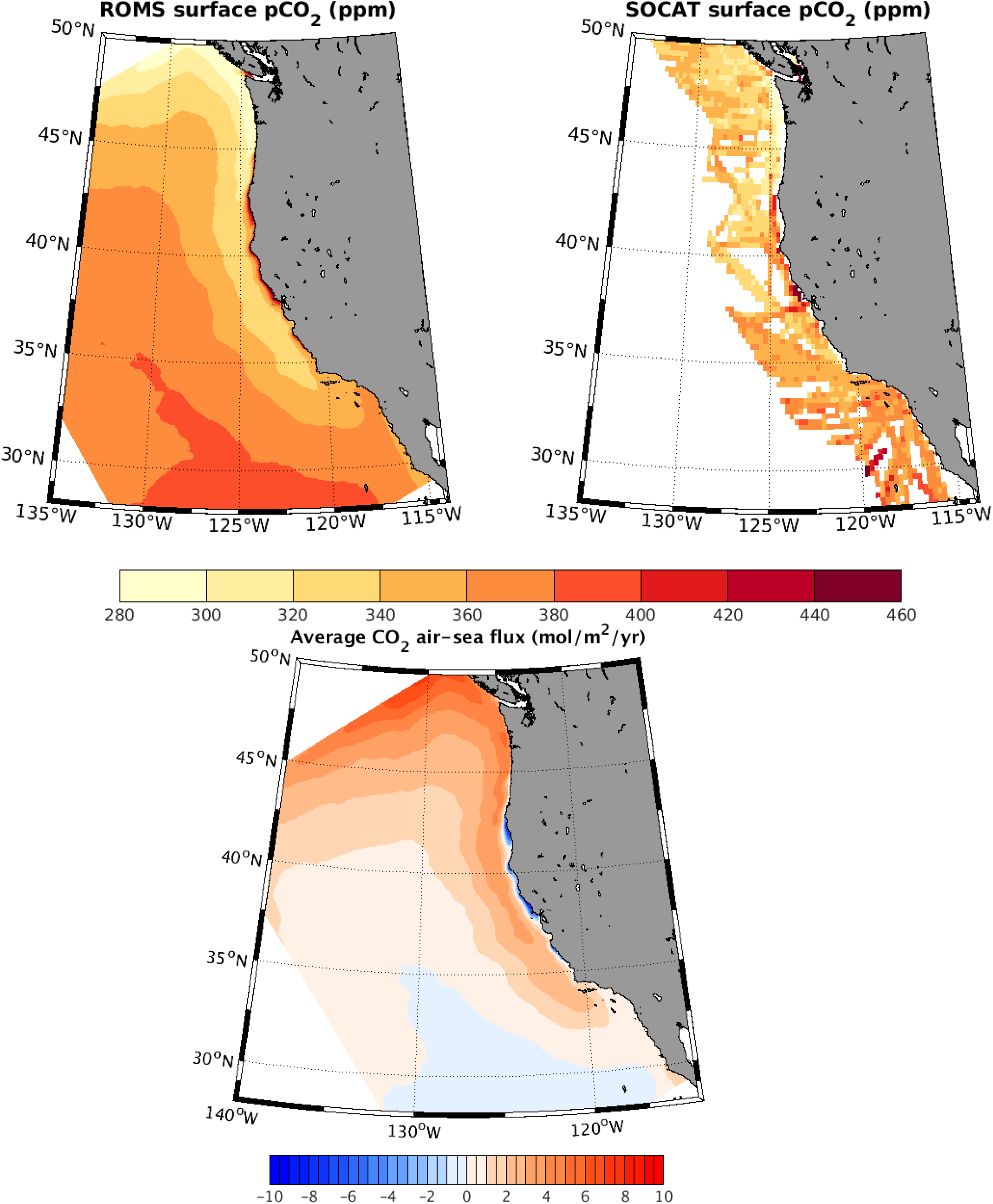
Surface ocean *CO*_2_ partial pressure (ppm) and air-sea flux (mol m^−2^ yr^−1^). (a) Annual surface *pCO*_2_ from ROMS. (b) Surface *pCO*_2_ from SOCATv6 gridded coastal dataset, averaged over all months from 1995-2010. (c) Annual airsea *CO*_2_ flux from ROMS. Negative values indicate outgassing to the atmosphere.

The lack of apparent regional or seasonal bias in the the surface *pCO*_2_, together with the good model-data agreement in surface buoyancy fluxes (Renault et al., 2021) and biological rates in these simulations, suggests that the balance of processes regulating the model’s surface *CO*_2_ fluxes is reliably captured. Consistent with previous studies, the net integrated *CO*_2_ flux to the atmosphere (an uptake of 1.4 Tg C yr-1) is found to be a relatively small residual of larger compensating outgassing and ingassing fluxes. A detailed accounting of factors driving *pCO*_2_ and air-sea flux variability has been described (Fiechter et al., 2014; Turi et al., 2014). Our model predicts a somewhat larger net uptake than previous studies, likely because our model domain extends farther into the northern CCS where the combination of heat loss and fresh water forcing suppresses surface *pCO*_2_. An extension of similarly detailed analysis of air-sea flux variability to the northern CCS domain is left to future work.

### 3.2 Thermocline

Here we test the model’s representation of biogeochemical properties below the mixed layer and photic zone, *i.e.,* in the thermocline. We focus on distributions and variability of *O*_2_ and aragonite saturation state (Ω*_A_*) as these properties influence habitability for calcification and aerobic respiration by marine animals. The values of Ω*_A_* are calculated from model dissolved inorganic carbon (DIC) and total alkalinity (Alk) using carbonate system equilibrium equations (*CO*_2_*SYS*) (van Heuven et al., 2011). The variability of *O*_2_ has been analyzed in greater detail using previous simulations of this model with different atmospheric and physical boundary conditions (Durski et al., 2017).

To evaluate the vertical structure of biogeochemical tracers, we turn to repeat hydrographic sections (Fig. 16). Transects through three cross-sections spanning the Southern California Bight (CalCOFI line 80), the central California coast (MBARI line 67), and the central Oregon coast (Newport line) show the typical vertical and cross-shelf gradients of 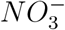 and *O*_2_. The downward enhancement of 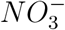 and depletion of *O*_2_ is a signal of the broad-scale vertical redistribution of these elements by the formation and degradation of organic matter within the CCS, as well as the gradients imported from the Pacific basin through the boundary conditions. The shoaling of the isopleths of both quantities follows that of the isopycnal surfaces by upwelling along the coast. The distributions of 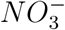 and *O*_2_ are generally well reproduced by ROMS-BEC. The model somewhat underestimates the slope of these isopleths very nearshore. This tendency is also reflected in, and likely derived from, the same underestimate in the zonal tilt of isopycnal surfaces (Renault et al.,2021).

**Figure 16:**
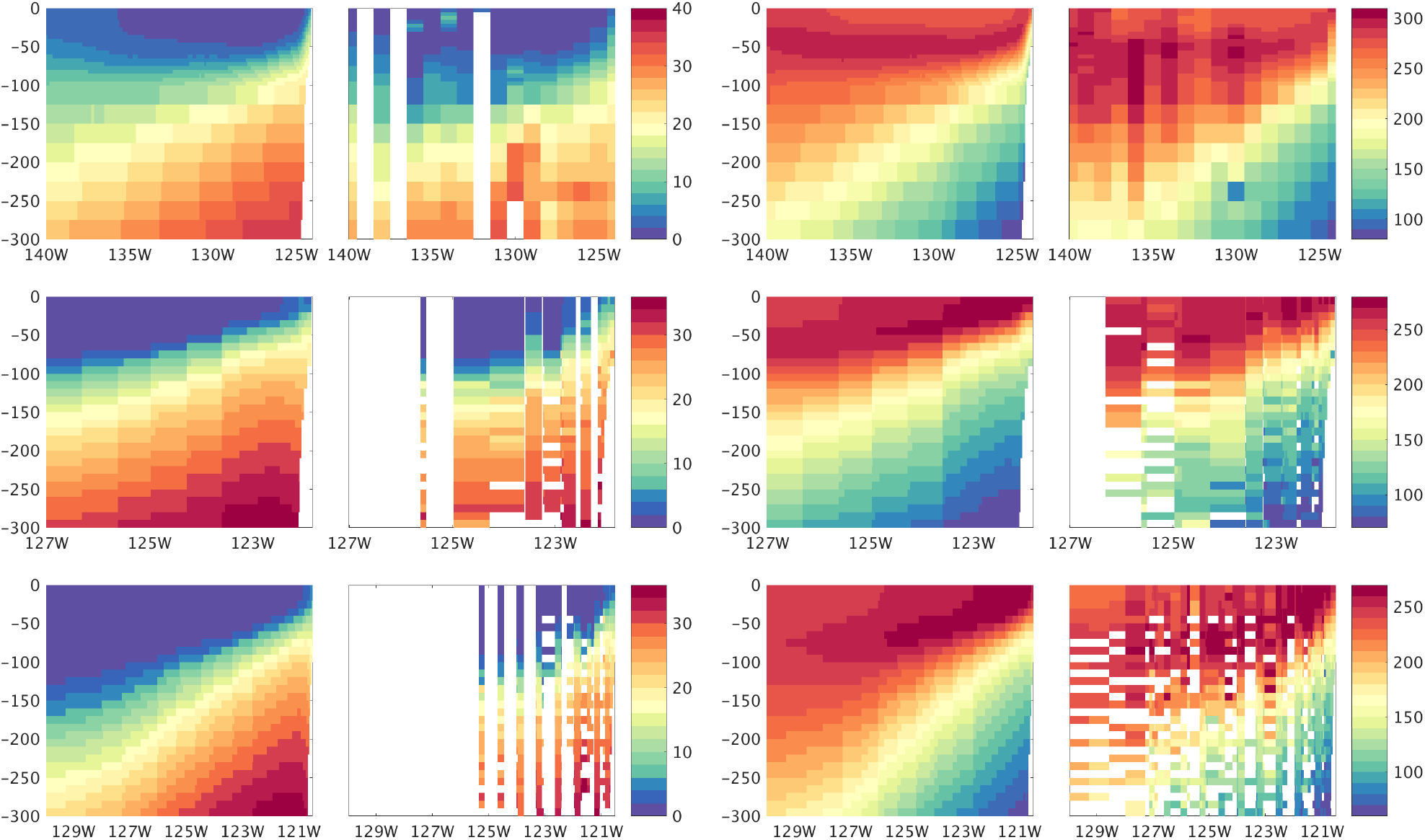
Vertical sections of annual mean 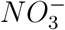(left) and *O*_2_(right) (mmol m^−3^), from ROMS (left) and WOD (right) at the latitudes with regular observations by repeat hydrographic surveys. The lines span the northern CCS (upper row, ≈ 44.5°N, nearest the Newport OR), the central CCS (middle row, MBARI line 67), and the southern CCS (bottom row, CalCOFI line 80). Locations of observations are shown in Fig. 3.

Along isopycnal surfaces, *O*_2_ generally increases with latitude and with distance from shore, reflecting the contrasting properties carried by the broad offshore California current from the *O*_2_-rich subarctic, and the narrow near-shore California Undercurrent that transports low-O_2_ waters of tropical origin northward along the slope. Both northern and southern end-member water types can be seen on the isopycnal surface 26.5 (Fig. 17), which also comprises the source of water upwelling onto the continental shelf along much of the US west coast (Pierce et al., 2012). On this and other density surfaces, the distribution of *O*_2_ in ROMS-BEC is consistent with climatological observations, suggesting that the balance of distinct water masses and the respiratory modifications they experience in the interior of the domain are relatively well represented in the model. Thermocline nutrient distributions exhibit a similar model skill (not shown).

**Figure 17:**
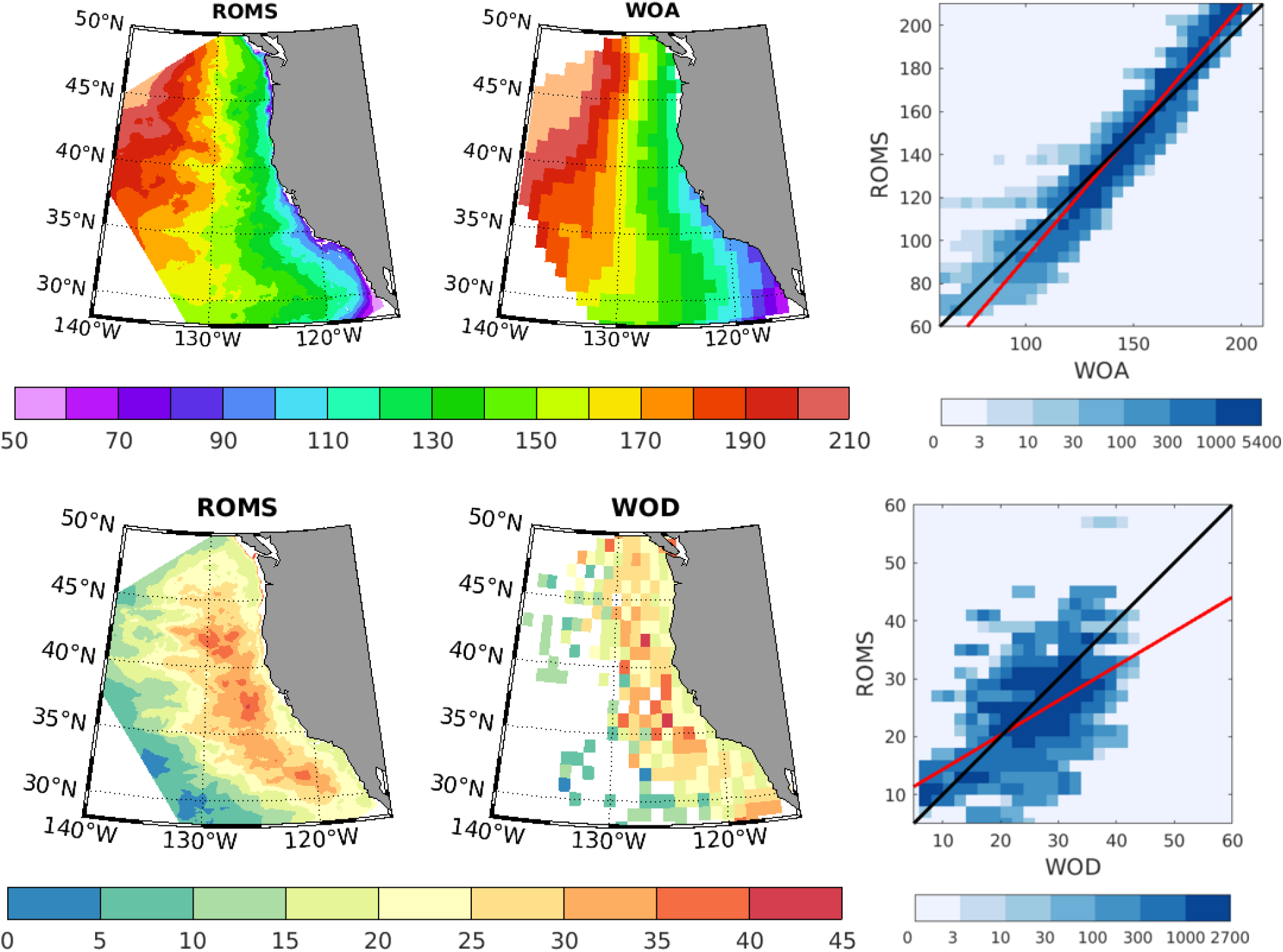
Thermocline *O*_2_ in model simulations and observations of the climatological mean (upper) and standard deviation (lower). All maps are interpolated to a potential density surface (26.5) surface chosen as the density class of waters upwelling onto the shelf in summer (a.k.a. “source water”). The mean *O*_2_ maps are from summer months (JJA), however other seasons reveal similar model fidelity. Observed mean summer *O*_2_ is from objectively mapped climatology (WOA). Variability is mapped as the standard deviation of monthly values from the World Ocean Database covering 1955-2013 (lower right), and it is predominantly due to interannual variability rather than the seasonal cycle (see text). The correlations between ROMS and WOD are highly significant (p≪0.01) for both mean *O*_2_ and its variability, with squared Pearson correlation coefficients (R^2^) of 0.95 for the mean state and 0.35 for the spatial pattern of temporal variability.

The *O*_2_ in the thermocline of the CCS is highly variable, with standard deviations of ≈ 20 mmol m^−3^ that are on average 15-20% of the mean *O*_2_ across the historical measurements (Fig. 17). The magnitudes and patterns of variance are well captured by the ROMS hindcast. In both the model hindcast and in observations, the standard deviation of monthly *O*_2_ is ≈ 5 times larger than that of the climatological seasonal cycle. Thus, a large majority of the *O*_2_ variability is explained by non-seasonal time-scales, including large eddy-driven fluctuations (Frenger et al., 2018) and low-frequency climate variability (Buil and Lorenzo, 2017; McClatchie et al., 2010). Both model and observations indicate that *O*_2_ variation peaks slightly offshore, in a pattern resembling that of eddy kinetic energy (Renault et al., 2021), and reflecting the role of eddies in transporting hypoxic waters offshore (Frenger et al., 2018). The high interannual to decadal *O*_2_ variability observed in the central North Pacific, which reaches its maximum on the isopycnal 26.5, is not included in the climatological boundary conditions, and thus likely accounts for the model bias toward low *O*_2_ variance in the north of the domain.

Variations in *O*_2_ on depth surfaces in the thermocline are also dominated by interannual anomalies. The fluctuating *O*_2_ at depths of 100-200 m are highly correlated with density (*R*^2^ ≥ 0.5) throughout the CCS, reflecting the importance of isopycnal heaving of the background *O*_2_ gradient (Ito et al., 2019). Similar correlations are observed in ROMS and the World Ocean Database (Fig. 18). In the model, the largest such anomaly is associated with the ENSO event in 1997-98, in which deepened isopycnals yield high *O*_2_ conditions that last for ≈ 1 year. The signal is recorded in the central CCS as well, although the magnitude of the anomalies is reduced by ≈ 50% relative to the better-sampled event in the SCB.

**Figure 18:**
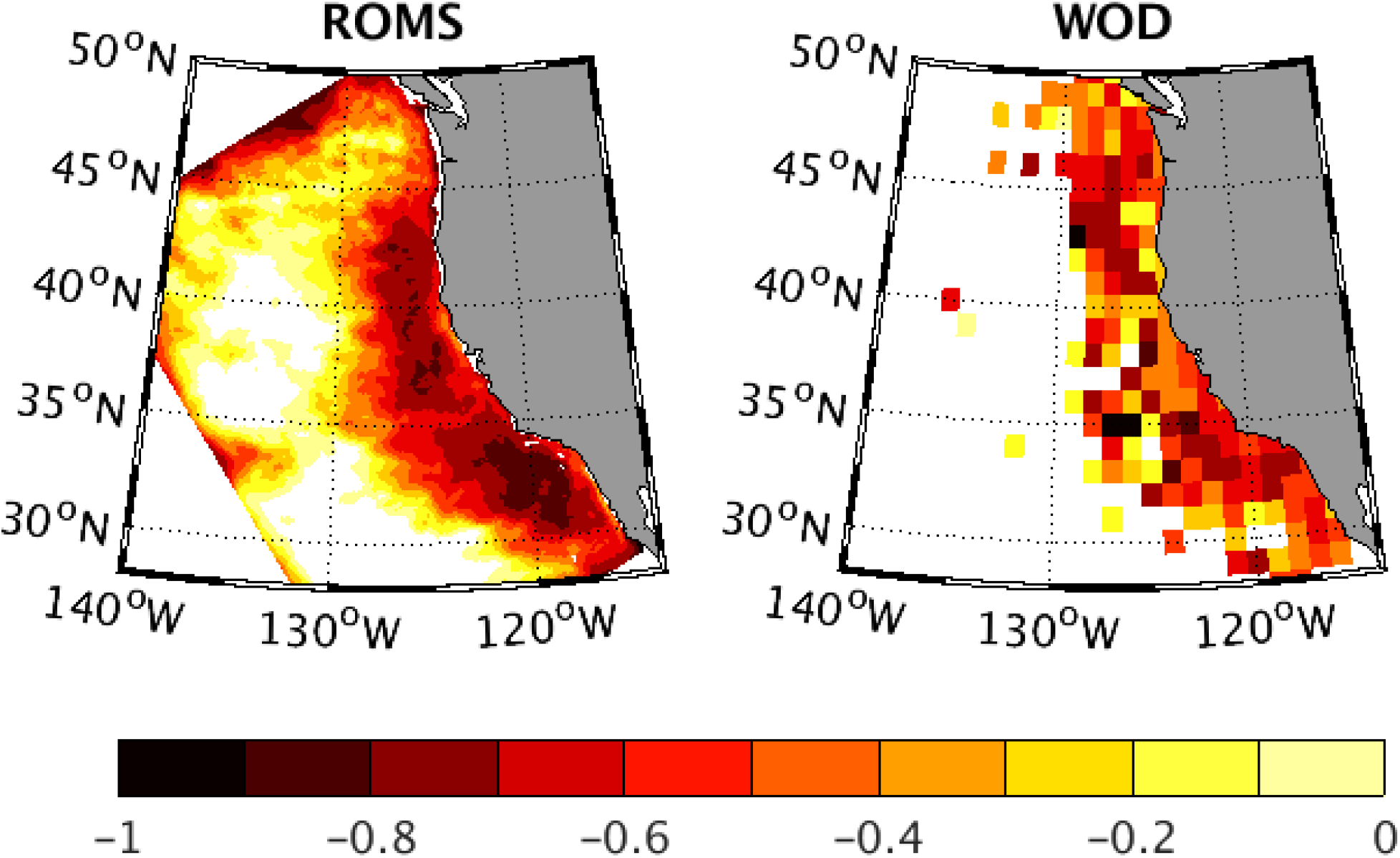
Correlation between *O*_2_ and density anomalies at 100 m, in ROMS-BEC (left) and WOD (right). Interannual variations in subsurface *O*_2_ are highly correlated with density (*R*^2^≥ 0.5) throughout the CCS, and the strength of the relationship is similar in model and observations.

The variability of *O*_2_ within an isopycnal surface can be used to account for this portion of variance, leaving only lateral circulation and respiration. We find that along *σ_θ_* = 26.5 kg m^−3^, a large fraction of *O*_2_ variability is correlated with salinity (Fig. 19), commonly used as a proxy for tropical low-oxygen and high-salinity water transported poleward in the coastal undercurrent (Meinvielle and Johnson, 2013). Interannual variability in respiration rates on this surface also accounts for ≈ 20% of isopycnal *O*_2_ variance, and is in turn correlated with the depth of the density layer (Deutsch et al., 2011). Historical observations show a declining strength of correlation between S and *O*_2_ with latitude, suggesting that variability from dynamics other than the CUC become an increasingly important source of *O*_2_ variability to the north. Indeed, the variability of *O*_2_ in central mode water from the open North Pacific is most pronounced on this density surface (**?**Buil and Lorenzo, 2017), and the North Pacific Current is thought to transport isopycnal *O*_2_ anomalies into the northern CCS and contribute to the significant interdecadal fluctuations observed in the CCS [insert citation in comment here]. However, this longer term variance along isopycnals is not represented in the climatological biogeochemical boundary conditions in the model. Thus the relatively constant correlation between S and *O*_2_ along isopycnal surfaces across latitude in the model may stem from the lack of *O*_2_ variability on isopycnal surfaces as those waters enter the domain from the open North Pacific. A complete attribution of the observed magnitudes of *O*_2_ variance in source waters to the CCS will require inclusion of anomalies entering from the broader North Pacific (Deutsch et al., 2006; Kwon et al., 2016) and is left for future work.

**Figure 19:**
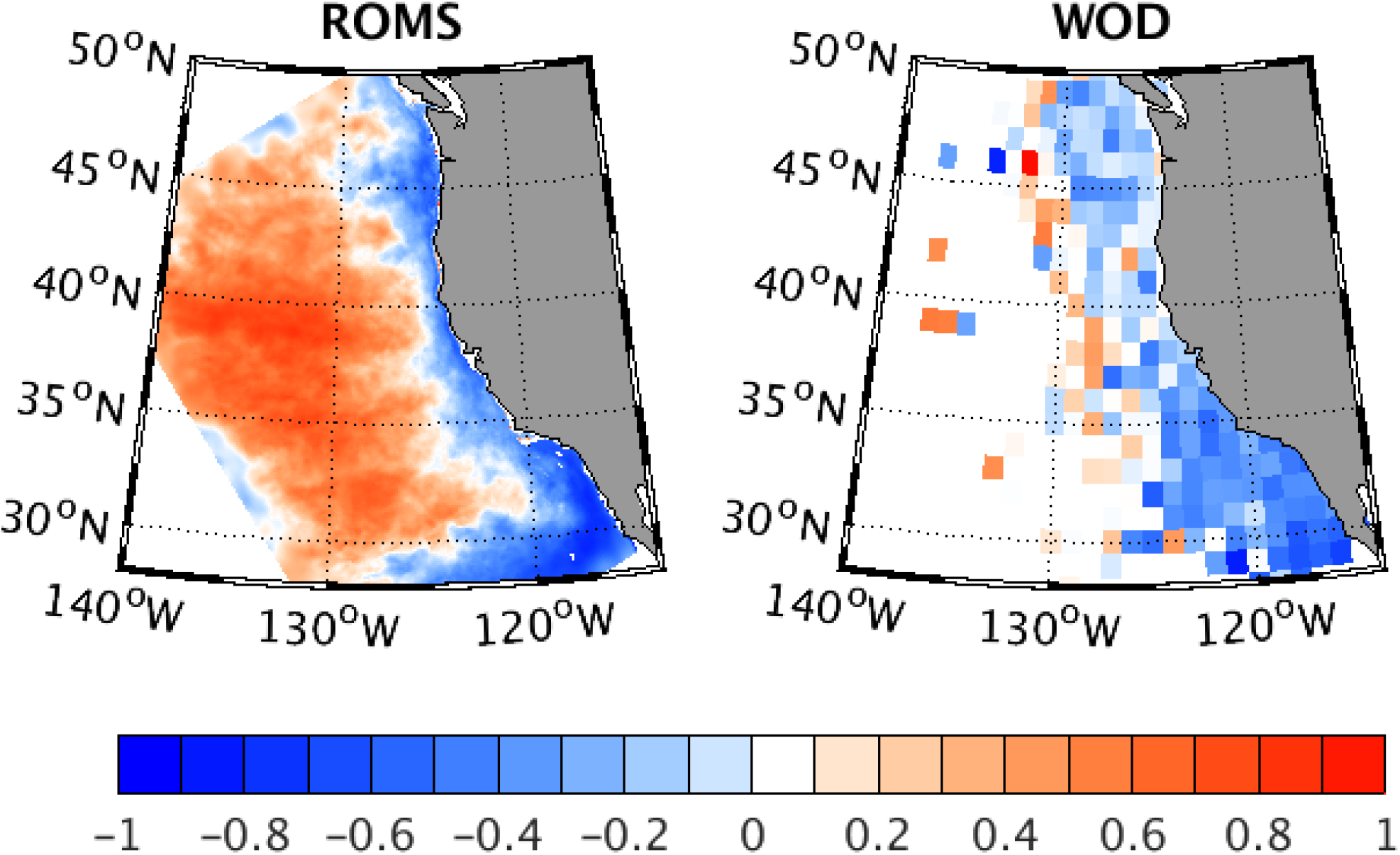
Correlation coefficient between *O*_2_ and salinity anomalies in (left) ROMS, and (right) WOD along the isopycnal surface *σ_θ_* = 26.5 kg m^−3^. Prevailing negative values indicate that high S occurs when *O*_2_ is low. To maximize data availability, the observational correlation is based on monthly anomalies in WOD from the period 1955-2013. The correlations are of similar magnitude when confined to the simulated period, 1995-2010, but are only available in the Southern California Bight.

In addition to low coastal *O*_2_, the CCS is characterized by shallow depth horizons for carbonate saturation (Fig. 20). Below ≈ 100 m depth, carbonate concentrations are commonly undersaturated with respect to aragonite mineral formation, and thus inhibit shell formation by calcifying organisms. The aragonite saturation state, Ω*_A_*, is predicted to fall below saturation (Ω*_A_* < 1) along most of the coast in summer, consistent with observations in NOAA coastal surveys (e.g.,Feely et al. (2008)). Coastal hydrographic surveys reveal a strong mesoscale patchiness to the carbonate saturation state, likely reflecting mesoscale eddies and submesoscale features. In the multi-annual mean distribution of Ω*_A_*, the model hindcast captures the scale and intensity of undersaturated conditions well.

**Figure 20:**
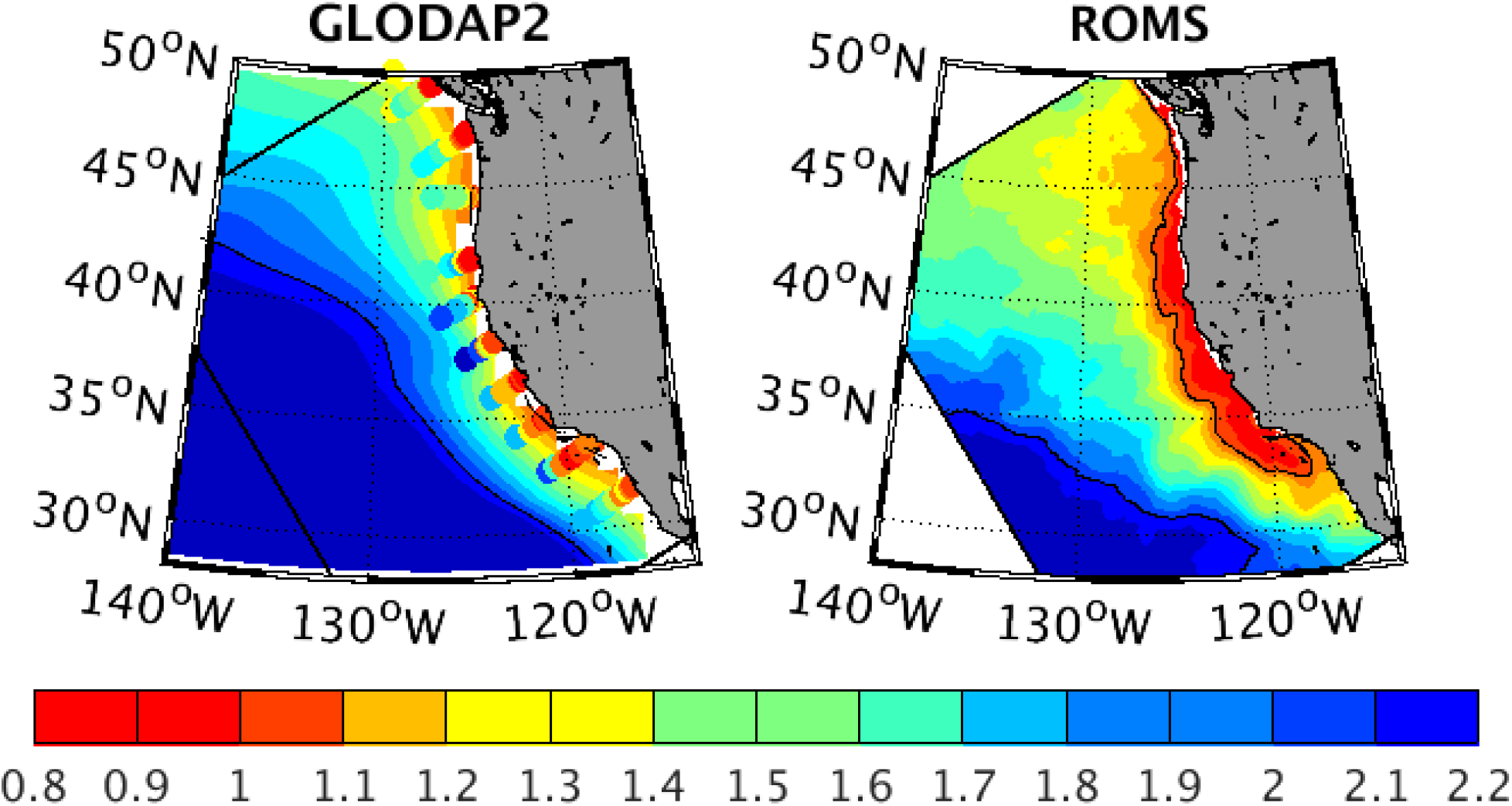
Aragonite saturation state (Ω*_A_*) at 100 m from observations (left) and ROMS-BEC (right). Observations are from large-scale objectively analyzed fields (color field, GLODAP2 (Lauvset and coauthors, 2016)), and from NOAA coastal surveys in the summer of 2007 (circles, Feelyet al. (2008)). Model distribution is averaged over summer (JJA) from 2004-2010 climatology.

Both low *O*_2_ and low Ω*_A_* have been implicated as primary factors mediating the influence of climate on organism fitness and species habitat in the CCS (Howard et al., 2020; Busch and McElhany, 2016). We compare decadal trends in both these properties from the hindcast simulations to the observed changes over time. For Ω*_A_*, the measurements are too sparse and the distribution too patchy to define a robust trend, even over the short model period. For each property, time series are shown for the regions with the most data coverage (Fig. 21): northern CCS for aragonite saturation, and southern CCS for *O*_2_. In both cases, the trend in the data is within the uncertainty in the measurements.

**Figure 21:**
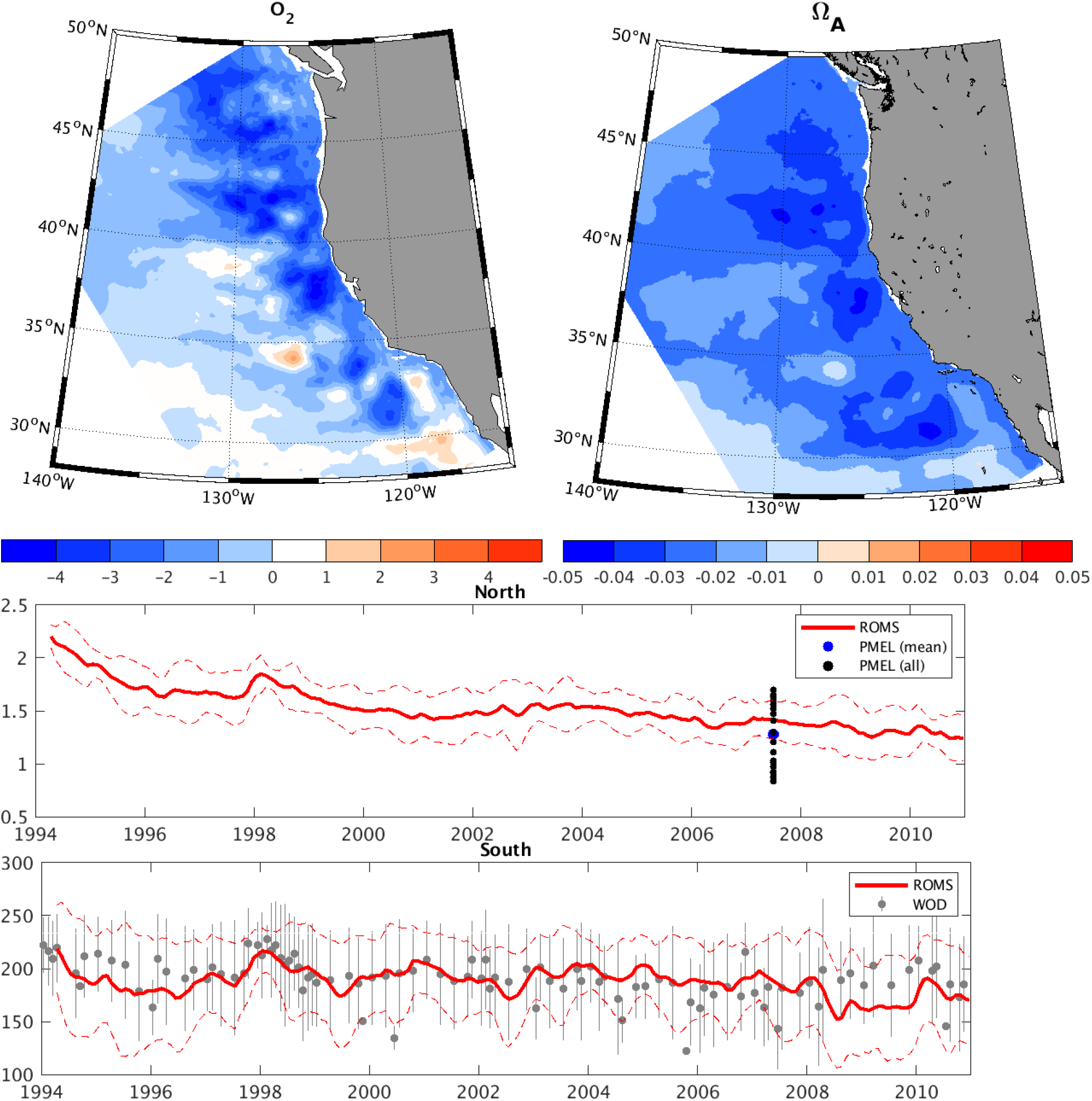
Trends in thermocline *O*_2_ and Ω*_A_* over the simulated period. Maps of the linear trend are shown in upper panels. Time series (lower panels) are shown for the regions with the most data coverage for each tracer: northern CCS for carbonate, and southern CCS for *O*_2_. For Ω*_A_* all available profiles are shown. For *O*_2_, the mean value and standard deviation are plotted as box-whisker for each month in the WOD.

As a metric of variability in these habitat constraints, we computed the volume of water subject to hypoxic or corrosive conditions. We use a constant *O*_2_ level of 100 mmol m^−3^ as a simple indicator of hypoxic constraints, recognizing that this value varies among species, and depends on other factors, including temperature (Deutsch et al., 2015). Corrosive conditions are defined by simple thermodynamic undersaturation (Ω*_A_* < 1), though biological sensitivities may begin at higher thresholds. Water volumes are computed as the sum of grid cell volumes with *O*_2_< 100 mmol m^−3^ or Ω*_A_* < 1 that are on the continental shelf (– *z* < 200 m).

The volume of hypoxic and corrosive water in the CCS varies strongly over latitude and time (Fig. 22). For both properties, restriction of putative habitat volume is stronger to the north of Pt. Conception, opposite the latitudinal gradient of *O*_2_ and Ω*_A_*. The corrosive volumes are much larger than hypoxic volumes, exceeding 90% of water volume in northern latitudes during the summer upwelling season, consistent with NOAA survey data. On an annual basis, waters with a more stringent criterion for calcification (Ω*_A_* < 2) are about twice as voluminous still, primarily because the length of the season with low carbonate is broadened. Hypoxic conditions occupy a smaller fraction of shelf waters, but reach 30% of shelf water volume over a broad latitude range. The fractional coverage by hypoxia peaks around 45 °N, on the Oregon coast, and it is quite small in the Southern California Bight, where *O*_2_ declines most sharply below 200 m rather than on the shelf.

**Figure 22:**
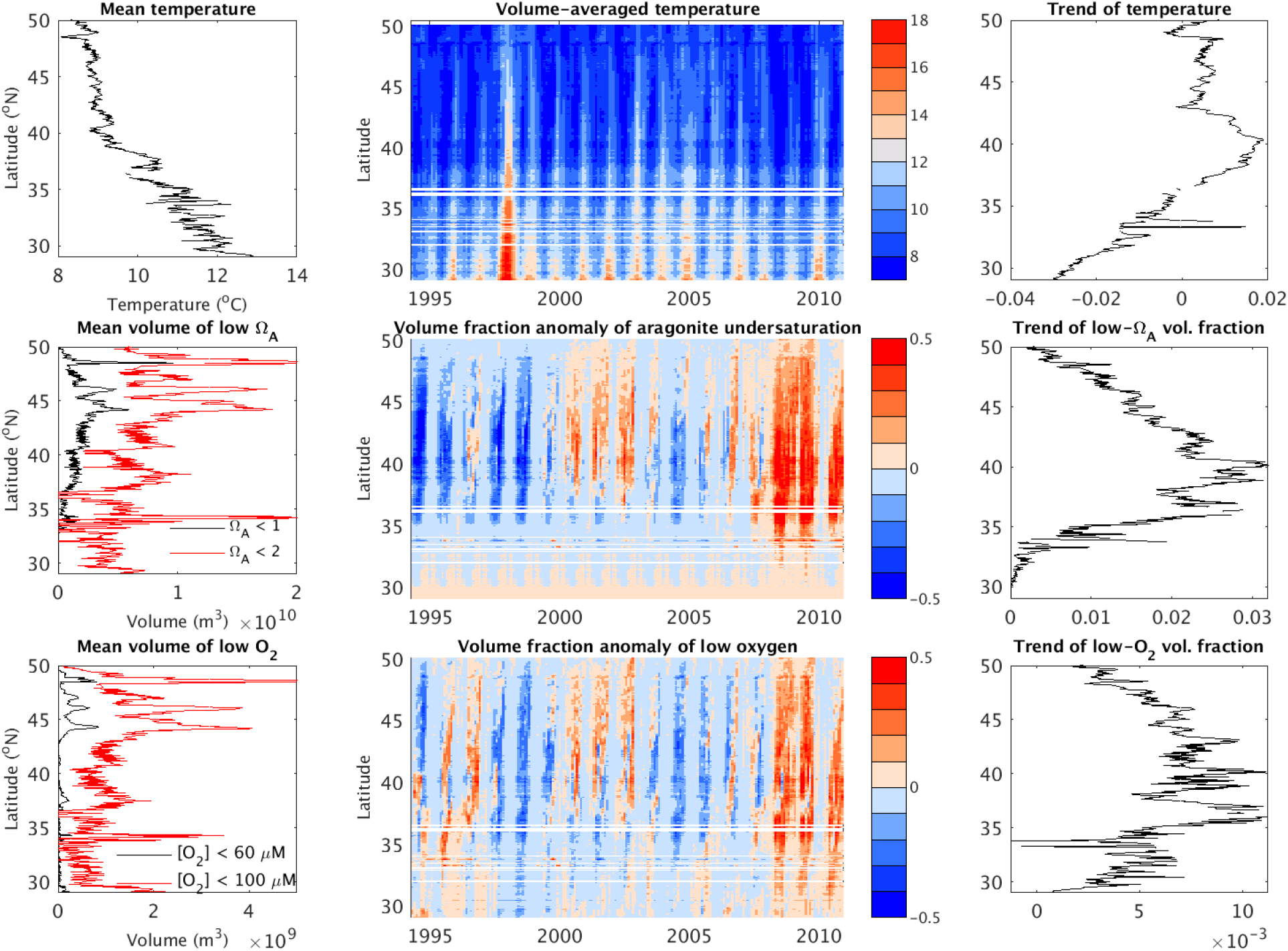
Top row: mean temperature and temporal trend (°C year^−1^) of water on the shelf (– *z* < 200 m). Middle and bottom rows: volume of corrosive and hypoxic water over time and latitude. Water volumes are computed as sum of grid cell volumes with *O*_2_ < 100 mmol m^−3^(bottom) and Ω*_A_* < 1 (middle) that are on the shelf (– *z* < 200 m). Mean values for each month are shown in the left column, and anomalies, computed as a fractional deviation from the climatological mean monthly volume at each latitude, in the middle column. Trends over time are shown in the right column.

An analogous figure is shown in Renault et al. (2021) for the along-coast and temporal variability of the sea-surface height and depth of the σ_θ_ = 26.5 kg m^−3^ depth anomalies. Relative to the quantities shown in Fig. 22, they exhibit more along-coast coherence and a more dominant seasonal cycle, with less evident interannual variability than shown here, apart from the 1997-98 ENSO event. This indicates somewhat smoother physical fields than biogeochemical ones, reflective of non-conservative biogeochemical processes acting on top of the broader patterns of physical circulation influence.

Variability in both habitat constraints is largely synchronous (Fig. 22c,d), reflecting the strong control on both *O*_2_ and Ω*_A_* by the effects of cumulative organic matter respiration. For both volumes, the fractional variation is similar and substantial, reaching ≈ 50% of the mean across much of the latitude range. Variability of corrosive volumes is greatly attenuated south of Pt. Conception. Simulated hypoxic volumes increased sharply during the early 2000s off Oregon, when major ecosystem die-offs were attributed to the onset of extreme hypoxia there (Chan et al.,2008).

## 4 Conclusions

We present model simulations of ecosystem and biogeochemical cycles in the CCS that reproduce the broad patterns of processes and states observed in this region over the past couple of decades. Our results demonstrate that productivity of the CCS reflects a complex interplay of factors. The limitation by the physical supply and removal of macronutrients (nitrate) provides the dominant seasonal and spatial pattern of NPP, but with significant constraints from light and *Fe* on a seasonal basis, especially in the northern CCS. Interannual variations in NPP are reasonably well predicted by fluctuations in pycnocline depth that modulate the rates of surface nutrient supply. Expanded datasets on near-surface *Fe* concentrations are needed to better establish its role as a limiting factor for growth in the CCS. A significant correlation between model NPP and surface irradiance suggests that changes in light are also influential. Together, these results suggest that an index of NPP that accounts for both regional pycnocline structure and cloud cover would be more skillful than one based only on coastal winds (e.g.,Bakun (1990); Jacox et al. (2018)). Our results highlight the value of continued measurements of the depth of the chlorophyll maximum.

Biogeochemical properties of subsurface waters in the CCS are also well reproduced by model simulations. The amplitude of interannual variability in 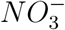 at the base of the photic zone and of *O*_2_ in the thermocline are also both strongly correlated to undulations of the pycnocline. The largest such anomalies in our simulation period were associated with the 1997-98 ENSO event, whose amplitude of density and *O*_2_ anomalies remains coherent over a wide latitude band, albeit with declining magnitude. For 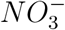, the overall variance is somewhat lower, and the strength of density correlations is somewhat higher in model output than in observations in the northern domain. This suggests an important role for anomalies entering the CCS from the subarctic North Pacific, an HNLC region. Basin-scale changes in biogeochemical properties are known to be exceptionally high at the gyre boundary ≈ 45°N (Mecking et al., 2008), and these remote anomalies are likely to play an important but uncertain role in the variability observed in the CCS. Similarly, inputs from terrestrial and riverine sources of nutrients and organic carbon could contribute to interannual variability in the system that is currently difficult to constrain from observational data. The impact of nearshore influences on the mean state suggests that its contribution to variability could also be substantial. Evaluating these remote influences from both the open ocean and from boundary inputs, using empirically-based time-dependent biogeochemical boundary conditions is an important avenue for future research.

The variability of biogeochemical properties leads to significant changes in the volume of waters characterized by biologically stressful conditions of hypoxia and carbonate undersaturation. In the volume anomalies for both habitat constraints, there is strong coherence across the CCS. Years with unusually large volumes of hypoxic or corrosive water offer few obvious latitudinal refuges. The onset of these conditions tends to propagate from the central CCS (≈ 40°N), arriving in the northern CCS with a 2-3 month delay. Thus, monitoring hypoxia and *CO*_2_ system parameters in the central CCS may offer some seasonal predictability for northern ecosystem impacts.

## Codes and Simulation Data

The physical and biogeochemical codes used for our simulations are at https://github.com/UCLA-ROMS/Code. Simulation model output archive data can be made available by email requests to the Corresponding Authors.

## Acknowledgments

This work was funded by grants from the National Science Foundation (OCE-1419323), the National Oceanic and Atmospheric Administration (DOC-NOAA NA15NOS4780186), and the Gordon and Betty Moore Foundation (GBMF #3775). Model simulations were carried out using the Extreme Science and Engineering Discovery Environment (XSEDE) and Yellowstone computers supported by the National Science Foundation at San Diego Supercomputer Center and NCAR, respectively. We gratefully acknowledge the PIs that supply high quality data to national centers, and to the programs that compile, manage and distribute these data for large-scale analyses, including the National Ocean Data Center (NOAA), the remote sensing products from NASA, and the California Cooperative Oceanic Fisheries Investigations (CalCOFI).

## 5 Appendix: Biogeochemical Model

Here, for completeness, we summarize the equations of the Biogeochemical Elemental Cycling (BEC) model in the implementation used for this work. This formulation is based on the original version presented in Moore et al. (2004).

### 5.1 Variables and parameters

#### 5.1.1 Prognostic variables

| Name | Description | Units |
| --- | --- | --- |
| $N_{NH_4}$ | ammonium | mmol $N/m^3$ |
| $N_{NO_2}$ | nitrite | mmol $N/m^3$ |
| $N_{NO_3}$ | nitrate | mmol $N/m^3$ |
| $N_{N_2O}$ | nitrous oxide | mmol $N/m^3$ |
| $N_{N_2}$ | nitrogen | mmol $N/m^3$ |
| $N_{don}$ | dissolved organic nitrogen | mmol $N/m^3$ |
| $P_{PO_4}$ | Phosphorus | mmol $P/m^3$ |
| $P_{dop}$ | dissolved organic phosphorus | mmol $P/m^3$ |
| $O_2$ | dissolved oxygen | mmol $O_2/m^3$ |
| $Fe$ | iron | nmol $Fe/m^3$ |
| $Fe_{sp}$ | small phytoplankton iron | mmol $Fe/m^3$ |
| $Fe_{diat}$ | diatom iron | mmol $Fe/m^3$ |
| $Fe_{diaz}$ | diazotroph iron | mmol $Fe/m^3$ |
| $Fe_{dofe}$ | dissolved organic iron | mmol $Fe/m^3$ |
| $Ca_{sp}$ | small phytoplankton calcium carbonate | mmol $CaCO_3/m^3$ |
| $Si_{diat}$ | diatom silicate | mmol $SiO_2/m^3$ |
| $Si_{SiO_2}$ | Silicate | mmol $SiO_2/m^3$ |
| $Alk$ | total alkalinity | mmol/$m^3$ |
| $C_{DIC}$ | dissolved inorganic carbon | mmol $C/m^3$ |
| $C_{sp}$ | small phytoplankton carbon | mmol $C/m^3$ |
| $C_{diat}$ | diatom carbon | mmol $C/m^3$ |
| $C_{diaz}$ | diazotroph carbon | mmol $C/m^3$ |
| $C_{doc}$ | dissolved organic carbon | mmol $C/m^3$ |
| $Chl_{sp}$ | small phytoplankton chlorophyll | mmol $Chl/m^3$ |
| $Chl_{diat}$ | diatom chlorophyll | mmol $Chl/m^3$ |
| $Chl_{diaz}$ | diazotroph chlorophyll | mmol $Chl/m^3$ |
| $C_{zoo}$ | zooplankton carbon | mmol $C/m^3$ |
| $POC_{sed}$ | organic carbon in sediment | mmol $C/m^2$ |
| $CaCO_{3,sed}$ | inorganic carbon in sediment | mmol $C/m^2$ |
| $Si_{sed}$ | Silicate in sediment | mmol $Si/m^2$ |
| $I_{SW}$ | Penetrative solar heat flux | $W/m^2$ |

#### 5.1.2 Local variables

| Name | Description | Units |
| --- | --- | --- |
| $J_{sp,no3}^u$ | nitrate uptake by small phytoplankton | mmol $N/m^3/sec$ |
| $J_{sp,nh4}^u$ | ammonium uptake by small phytoplankton | mmol $N/m^3/sec$ |
| $J_{diat,no3}^u$ | nitrate uptake by diatoms | mmol $N/m^3/sec$ |
| $J_{diat,nh4}^u$ | ammonium uptake by diatoms | mmol $N/m^3/sec$ |
| $J_{ammox}$ | rate of $NH_4$ oxidation to $NO_2$ | mmol $N/m^3/sec$ |
| $J_{nit}$ | rate of $NO_2$ oxidation to $NO_3$ | mmol $N/m^3/sec$ |
| $J_{no3,no2}^{denit}$ | amount of $NO_3$ converted to $NO_2$ by water column denitrif. | mmol $N/m^3/sec$ |
| $J_{sed}^{denit}$ | amount of $NO_3$ converted to $N_2$ by benthic denitrif. | mmol $N/m^3/sec$ |
| $J_{no2,n2o}^{denit}$ | amount of $NO_2$ converted to $N_2O$ by water column denitrif. | mmol $N/m^3/sec$ |
| $J_{no2,n2}^{denit}$ | amount of $NO_2$ converted to $N_2$ by water column denitrif. | mmol $N/m^3/sec$ |
| $J_{diaz}^{Nfix}$ | total $N_2$-fixation by diazotrophs | mmol $N/m^3/sec$ |
| $J_{diaz}^{Nexcrete}$ | $N$ excreted by diazotrophs | mmol $N/m^3/sec$ |
| $J_{diaz}^{photoN}$ | $N$ fixed by diazotroph (non-excreted) | mmol $N/m^3/sec$ |
| $J_{sp,po4}^u$ | phosphate uptake by small phytoplankton | mmol $P/m^3/sec$ |
| $J_{diat,po4}^u$ | phosphate uptake by diatoms | mmol $P/m^3/sec$ |
| $J_{diaz,po4}^u$ | phosphate uptake by diazotrophs | mmol $P/m^3/sec$ |
| $P_{diaz}^{remain}$ | remaining diazotrophs phosphate | mmol $P/m^3/sec$ |
| $J_{diaz,dop}^{loss}$ | non-grazing mortality of diazotrophs routed to POC | mmol $P/m^3/sec$ |
| $J_{diaz,dip}^{loss}$ | non-grazing mortality of diazotrophs routed to PIC | mmol $P/m^3/sec$ |
| $J_{sp,caco3}^{prod}$ | $CaCO_3$ production by small phytoplankton | mmol $CaCO_3/m^3/sec$ |
| $J_{sp,C}^{photo}$ | carbon uptake by photosynthesis in small phytoplankton | mmol $C/m^3/sec$ |
| $J_{diat,C}^{photo}$ | carbon uptake by photosynthesis in diatoms | mmol $C/m^3/sec$ |
| $J_{diaz,C}^{photo}$ | carbon uptake by photosynthesis in diazotrophs | mmol $C/m^3/sec$ |
| $J_{sp}^{grz}$ | grazing loss for small phytoplankton | mmol $C/m^3/sec$ |
| $J_{sp,dic}^{grz}$ | grazed small phytoplankton routed to DIC | mmol $C/m^3/sec$ |
| $J_{sp,doc}^{grz}$ | grazed small phytoplankton routed to DOC | mmol $C/m^3/sec$ |
| $J_{sp,poc}^{grz}$ | grazed small phytoplankton routed to POC | mmol $C/m^3/sec$ |
| $J_{sp,zoo}^{grz}$ | grazed small phytoplankton routed to new zooplankton biomass | mmol $C/m^3/sec$ |
| $J_{diat}^{grz}$ | grazing loss for diatoms | mmol $C/m^3/sec$ |
| $J_{diat,dic}^{grz}$ | grazed diatoms routed to DIC | mmol $C/m^3/sec$ |
| $J_{diat,doc}^{grz}$ | grazed diatoms routed to DOC | mmol $C/m^3/sec$ |
| $J_{diat,poc}^{grz}$ | grazed diatoms routed to POC | mmol $C/m^3/sec$ |
| $J_{diat,zoo}^{grz}$ | grazed diatoms routed to new zooplankton biomass | mmol $C/m^3/sec$ |
| $J_{diaz}^{grz}$ | grazing loss for diazotrophs | mmol $C/m^3/sec$ |
| $J_{diaz,dic}^{grz}$ | grazed diazotrophs routed to DIC | mmol $C/m^3/sec$ |
| $J_{diaz,doc}^{grz}$ | grazed diazotrophs routed to DOC | mmol $C/m^3/sec$ |
| $J_{diaz,poc}^{grz}$ | grazed diazotrophs routed to POC | mmol $C/m^3/sec$ |
| $J_{diaz,zoo}^{grz}$ | grazed diazotrophs routed to new zoo biomass | mmol $C/m^3/sec$ |
| $J_{sp}^{agg}$ | aggregation loss of small phytoplankton | mmol $C/m^3/sec$ |
| $J_{diat}^{agg}$ | aggregation loss of diatoms | mmol $C/m^3/sec$ |
| $J_{sp}^l$ | non-grazing mortality of small phytoplankton | mmol $C/m^3/sec$ |
| $J_{sp,dic}^l$ | non-grazing mortality of small phytoplankton routed to DIC | mmol $C/m^3/sec$ |
| $J_{sp,doc}^l$ | non-grazing mortality of small phytoplankton routed to DOC | mmol $C/m^3/sec$ |
| $J_{sp,poc}^l$ | non-grazing mortality of small phytoplankton routed to POC | mmol $C/m^3/sec$ |
| $J_{diat}^l$ | non-grazing mortality of diatoms | mmol $C/m^3/sec$ |
| $J_{diat,dic}^l$ | non-grazing mortality of diatoms routed to DIC | mmol $C/m^3/sec$ |
| $J_{diat,doc}^l$ | non-grazing mortality of diatoms routed to DOC | mmol $C/m^3/sec$ |
| $J_{diat,poc}^l$ | non-grazing mortality of diatoms routed to POC | mmol $C/m^3/sec$ |
| $J_{diaz}^l$ | non-grazing mortality of diazotrophs | mmol $C/m^3/sec$ |
| $J_{diaz,dic}^l$ | non-grazing mortality of diazotrophs routed to DIC | mmol $C/m^3/sec$ |
| $J_{diaz,doc}^l$ | non-grazing mortality of diazotrophs routed to DOC | mmol $C/m^3/sec$ |
| $J_{diaz,poc}^l$ | non-grazing mortality of diazotrophs routed to POC | mmol $C/m^3/sec$ |
| $J_{zoo}^l$ | zooplankton mortality | mmol $C/m^3/sec$ |
| $J_{zoo,dic}^l$ | zooplankton mortality routed to DIC | mmol $C/m^3/sec$ |
| $J_{zoo,doc}^l$ | zooplankton mortality routed to DOC | mmol $C/m^3/sec$ |
| $J_{zoo,poc}^l$ | zooplankton mortality routed to POC | mmol $C/m^3/sec$ |
| $f_{zoo}^d$ | fractional factor for routing of zoo losses | no units |
| $J_{sp}^{agg}$ | aggregation of small phytoplankton | mmol $C/m^3/sec$ |
| $J_{diat}^{agg}$ | aggregation of diatoms | mmol $C/m^3/sec$ |
| $J_{diaz}^{agg}$ | aggregation of diazotrophs | mmol $C/m^3/sec$ |
| $J_{don,nh4}$ | amount of dissolved organic $N$ remineralized | mmol $N/m^3/sec$ |
| $J_{doc,dic}$ | amount of dissolved organic $C$ remineralized | mmol $C/m^3/sec$ |
| $J_{dop,po4}$ | amount of dissolved organic $P$ remineralized | mmol $P/m^3/sec$ |
| $J_{dofe,fe}$ | amount of dissolved $Fe$ remineralized | mmol $Fe/m^3/sec$ |
| $J_{other}^{remin}$ | remineralization in the sediments by processes other than oxic remin. and denitrif. | mmol $C/m^3/sec$ |
| $J_{poc}^{prod}$ | amount of particulate organic $C$ produced | mmol $C/m^3/sec$ |
| $J_{pic}^{prod}$ | amount of particulate inorganic $C$ produced | mmol $C/m^3/sec$ |
| $J_{PSi}^{prod}$ | amount of particulate organic $Si$ produced | mmol $Si/m^3/sec$ |
| $J_{pfe}^{prod}$ | amount of particulate organic $Fe$ produced | mmol $Fe/m^3/sec$ |
| $J_{poc}^{remin}$ | remineralized particulate organic $C$ | mmol $C/m^3/sec$ |
| $J_{pic}^{remin}$ | remineralized particulate inorganic $C$ | mmol $C/m^3/sec$ |
| $J_{PSi}^{remin}$ | remineralized particulate organic $Si$ | mmol $SiO_2/m^3/sec$ |
| $J_{pfe}^{remin}$ | remineralized particulate organic $Fe$ | mmol $Fe/m^3/sec$ |
| $J_{Fe}^{scav}$ | scavenging of inorganic $Fe$ | mmol $Fe/m^3/sec$ |
| $J_{Fe}^{sed}$ | remineralized $Fe$ from sediment | mmol $Fe/m^3/sec$ |
| PAR | photosynthetically active radiation | W/m<sup>2</sup> |
| $\Phi_{poc}^{mineral}$ | incoming mineral associated particulate *C* flux | mmol *C* cm/m<sup>3</sup>/sec |
| $\Phi_{poc}^{out}$ | particulate *C*-flux buried in the sediments | mmol *C* /m<sup>2</sup>/sec |
| $\Phi_{poc}^{free}$ | free associated particulate *C* flux | mmol *C* /m<sup>2</sup>/sec |
| $\Phi_{pic}^{sol}$ | soluble and pic associated particulate *C* flux | mmol *C* /m<sup>2</sup>/sec |
| $\Phi_{pic}^{nonsol}$ | non-soluble and pic associated particulate *C* flux | mmol *C* /m<sup>2</sup>/sec |
| $\Phi_{psi}^{sol}$ | soluble and SiO<sub>2</sub> associated particulate *C* flux | mmol *SiO*<sub>2</sub> /m<sup>2</sup>/sec |
| $\Phi_{psi}^{nonsol}$ | non-soluble and SiO<sub>2</sub> associated particulate *C* flux | mmol *SiO*<sub>2</sub> /m<sup>2</sup>/sec |
| $\Phi_{dust}^{sol}$ | soluble and dust associated particulate *C* flux | mmol *C* /m<sup>2</sup>/sec |
| $\Phi_{dust}^{nonsol}$ | non-soluble and dust associated particulate *C* flux | mmol *C* /m<sup>2</sup>/sec |
| $\Phi_{pfe}^{sol}$ | soluble and dust associated particulate *C* flux | mmol *Fe* /m<sup>2</sup>/sec |
| $\Phi_{pfe}^{nonsol}$ | non-soluble and dust associated particulate *C* flux | mmol *Fe* /m<sup>2</sup>/sec |
| $\Phi_{poc}^{remin, sed}$ | remineralized organic *C* flux from sediment | mmol *C*/m<sup>2</sup>/sec |
| $\Phi_{pic}^{remin, sed}$ | remineralized inorganic *C* flux from sediment | mmol *C*/m<sup>2</sup>/sec |
| $\Phi_{Si}^{remin, sed}$ | remineralized Si flux from sediment | mmol *C*/m<sup>2</sup>/sec |
| $\Phi_{O_2}^{air}$ | *O*<sub>2</sub> air-sea flux | mmol *O*<sub>2</sub>/m<sup>2</sup>/sec |
| $\Phi_{N_2O}^{air}$ | *N*<sub>2</sub>*O* air-sea flux | mmol *N*/m<sup>2</sup>/sec |
| $\Phi_{N_2}^{air}$ | *N*<sub>2</sub> air-sea flux | mmol *N*/m<sup>2</sup>/sec |
| $\Phi_{CO_2}^{air}$ | *CO*<sub>2</sub> air-sea flux | mmol *C*/m<sup>2</sup>/sec |
| *Sc* | Schmidt Number | - |
| *W<sub>s</sub>* | Wind speed at 10 m | m/sec |
| *PV* | Piston Velocity | m/sec |

### 5.2 Ecosystem parameters

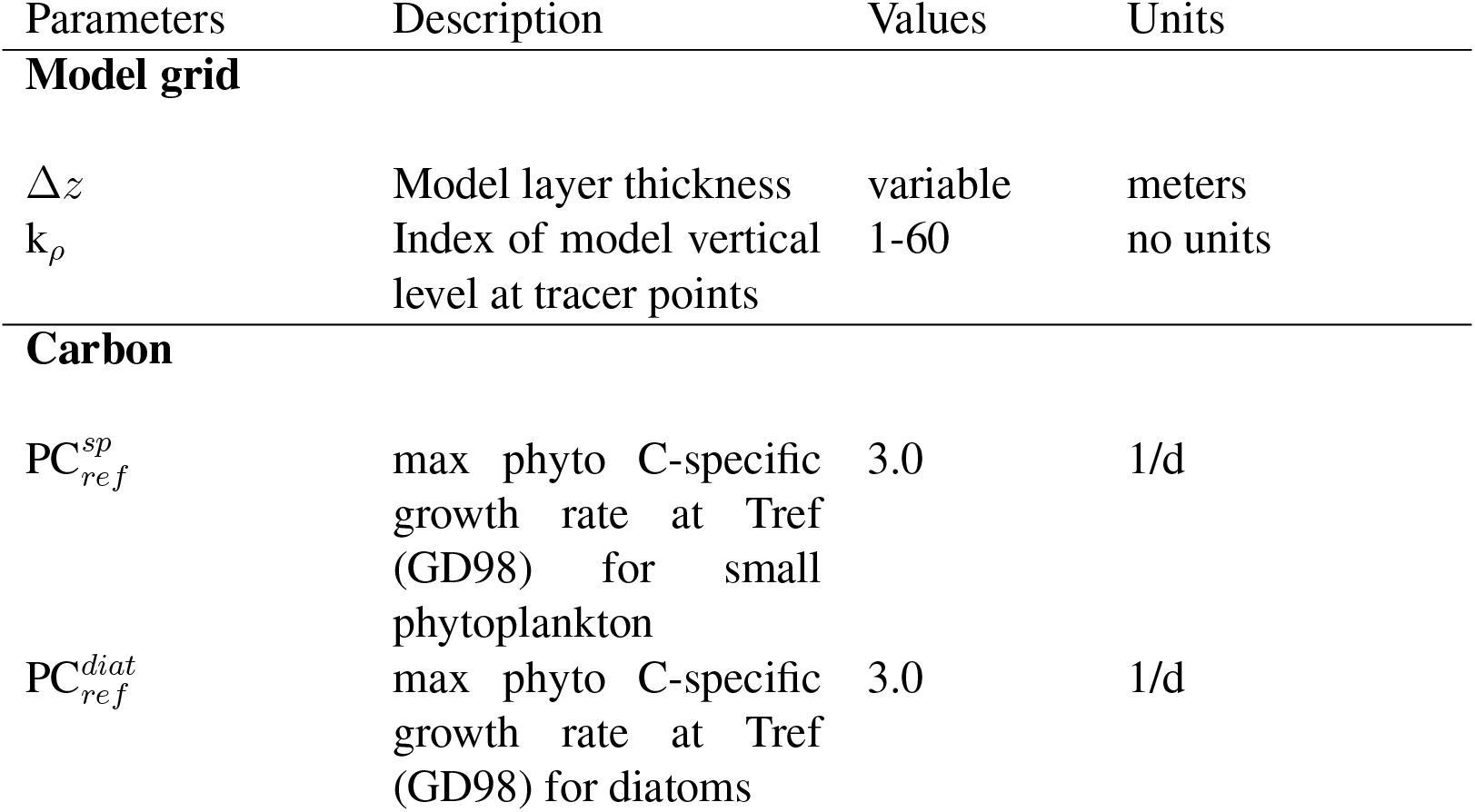
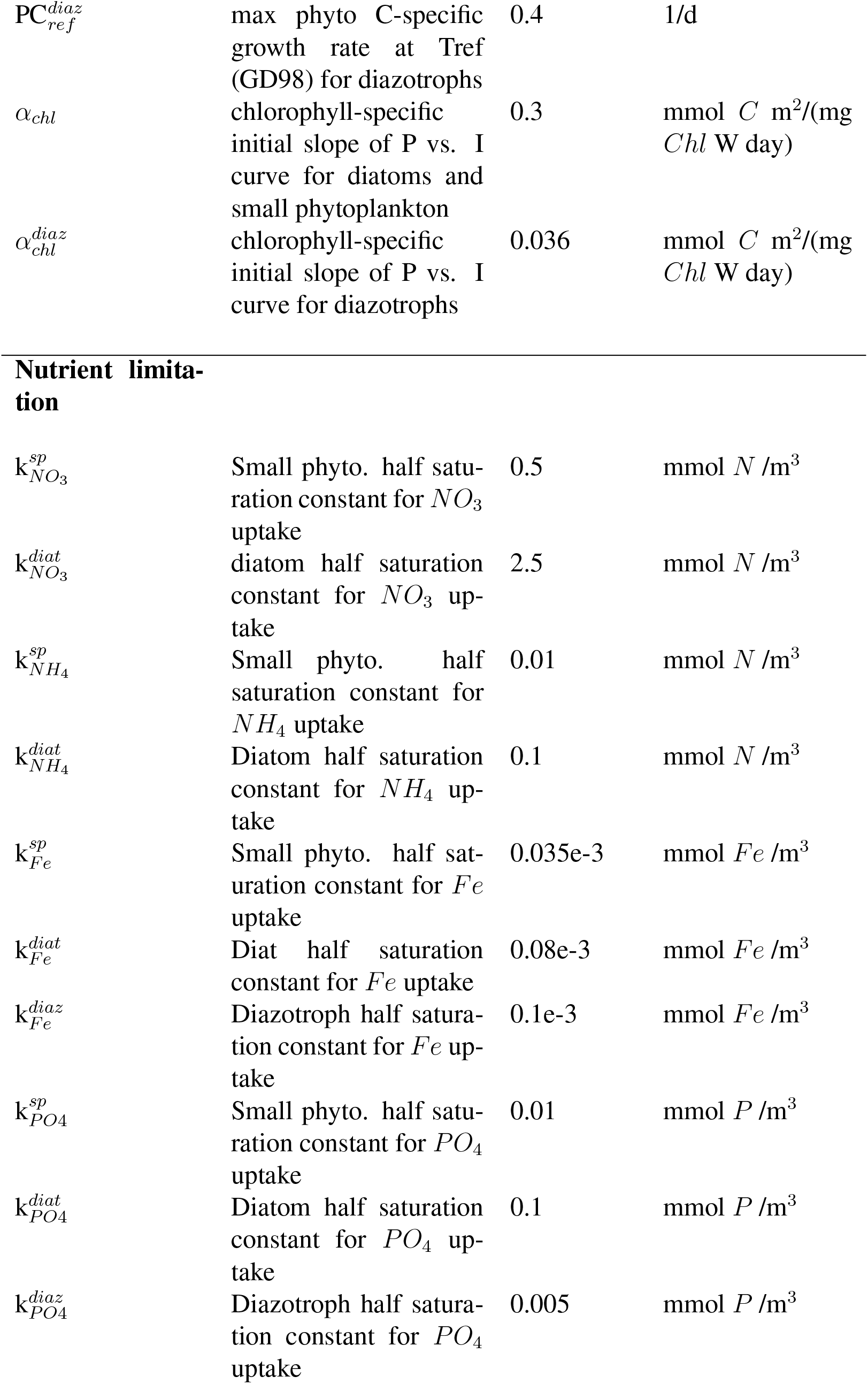
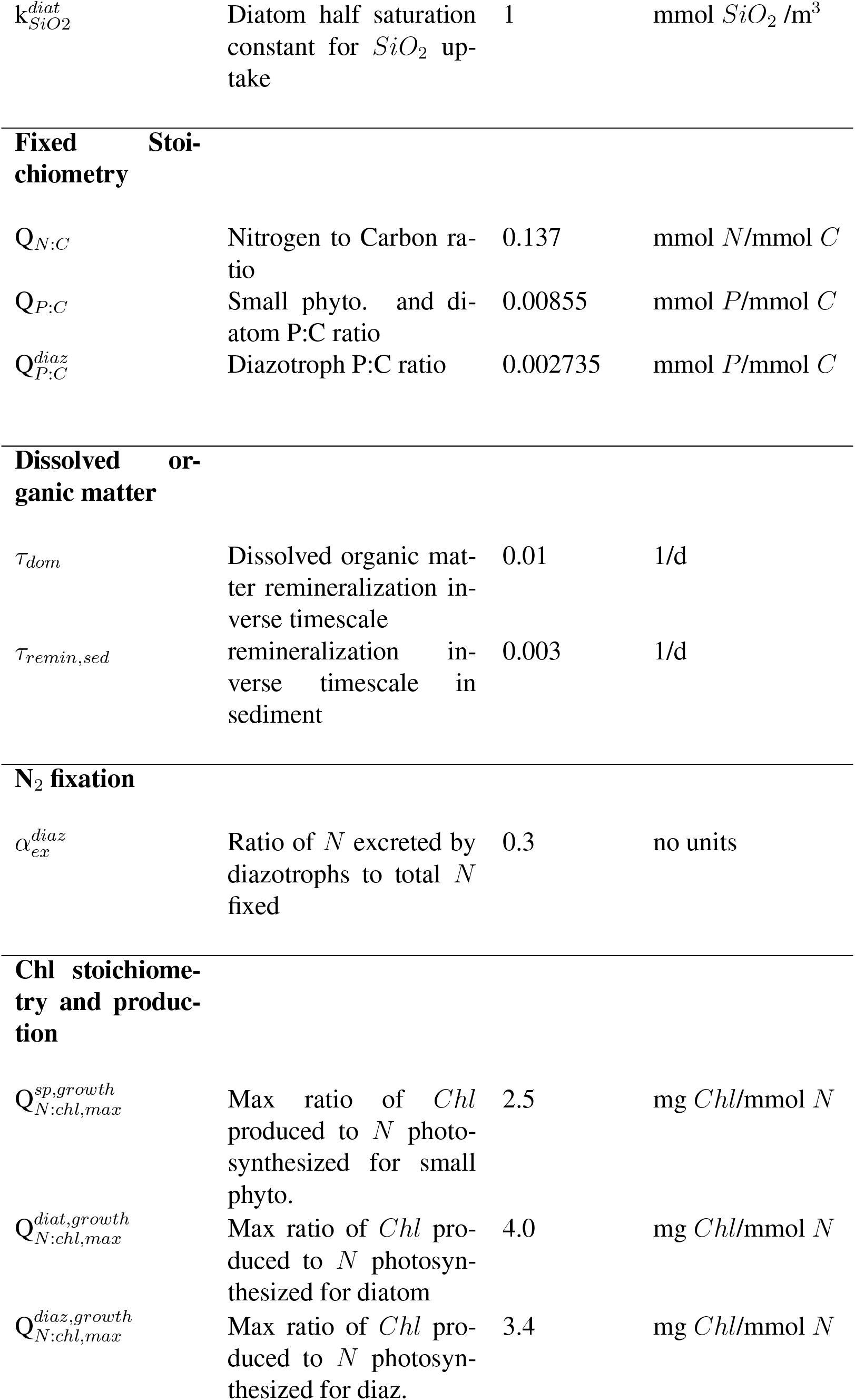
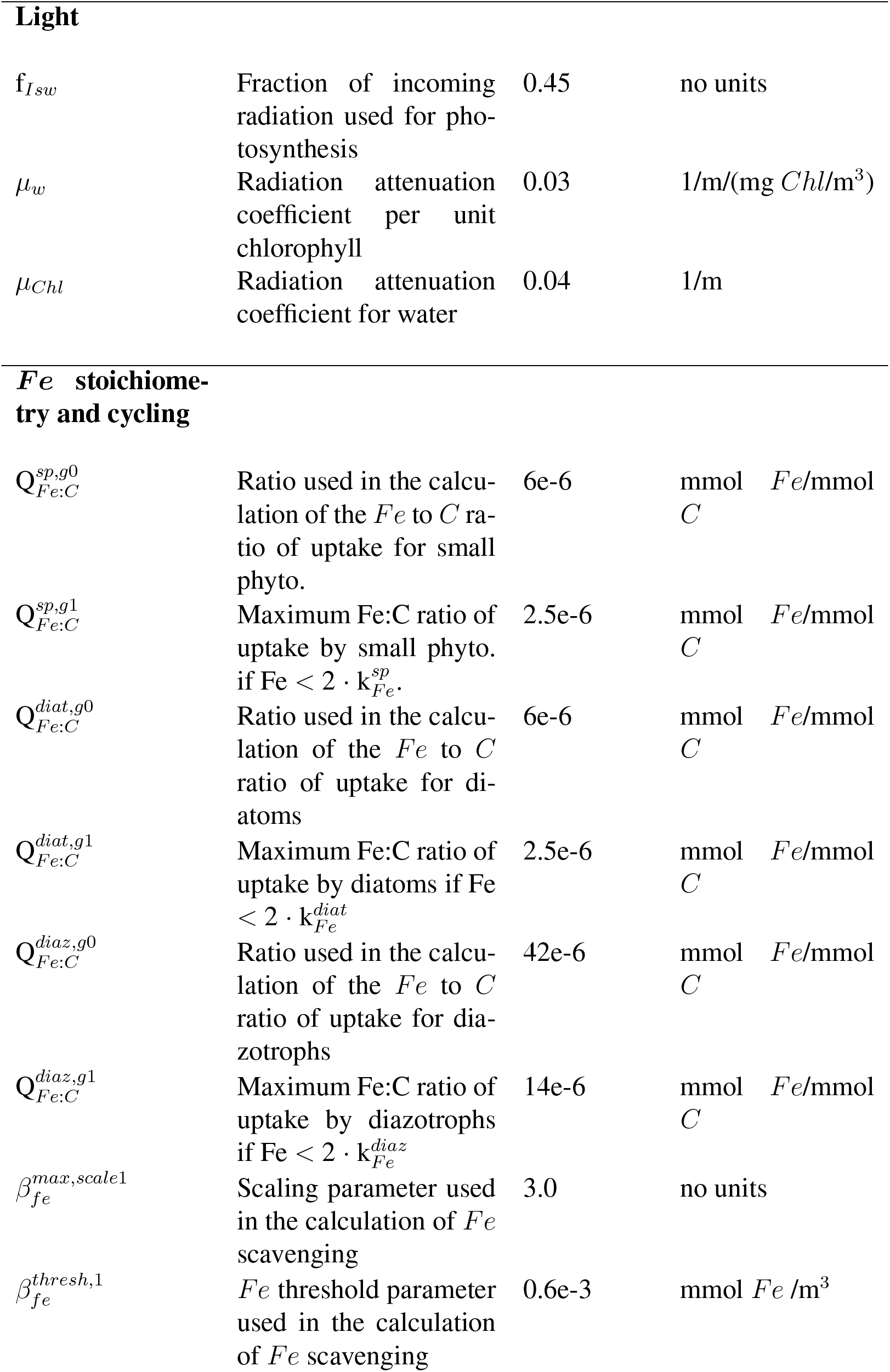
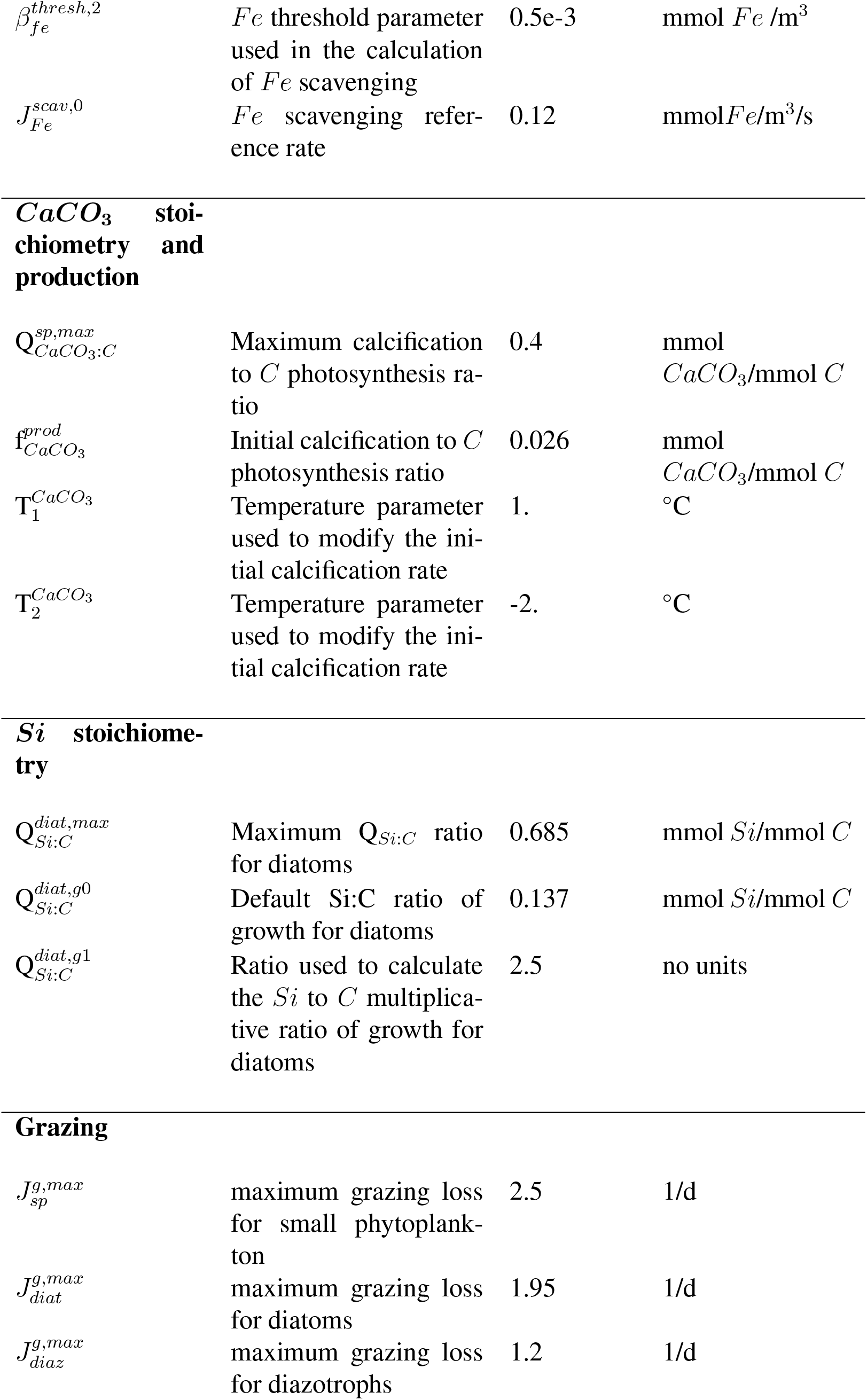
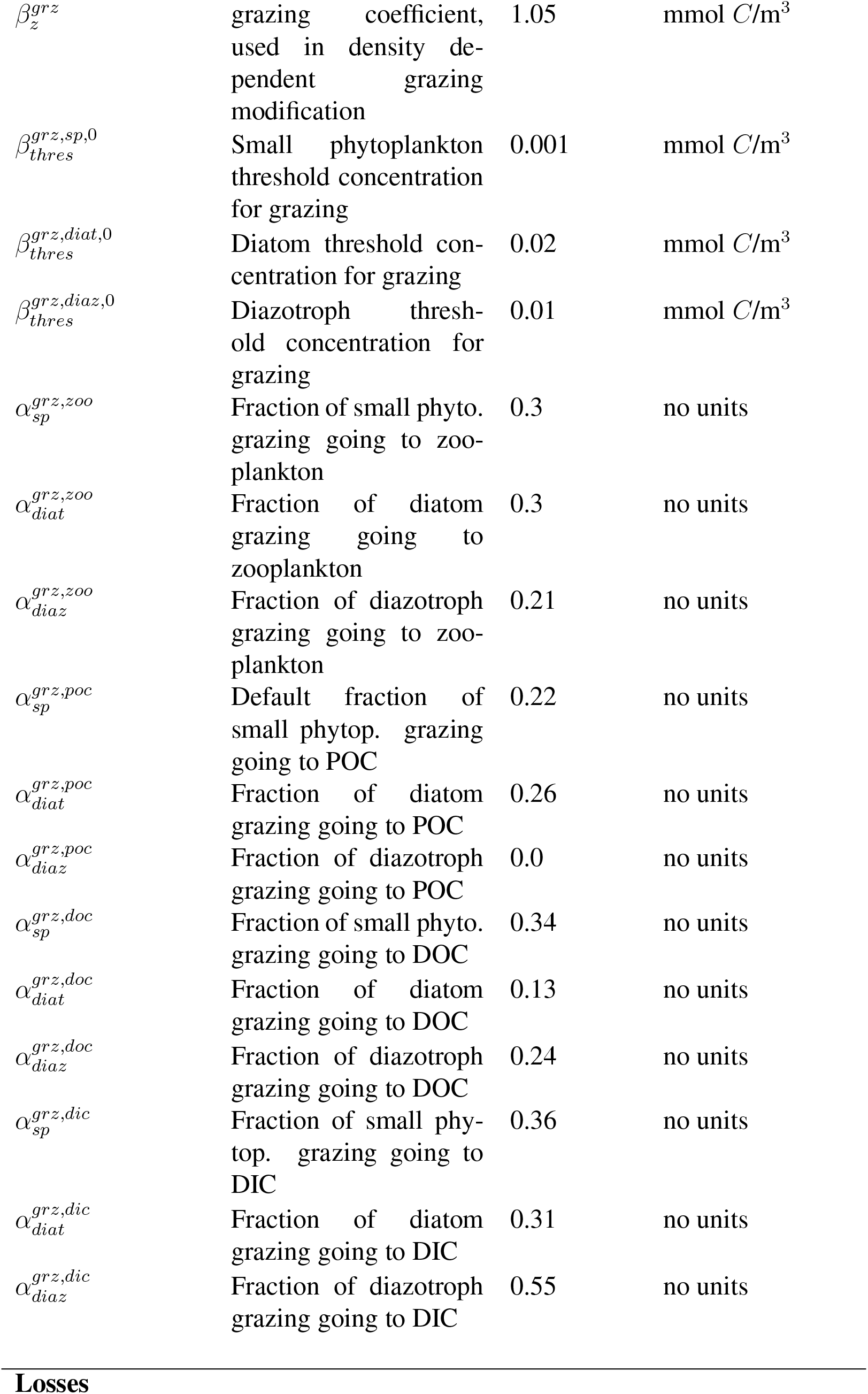
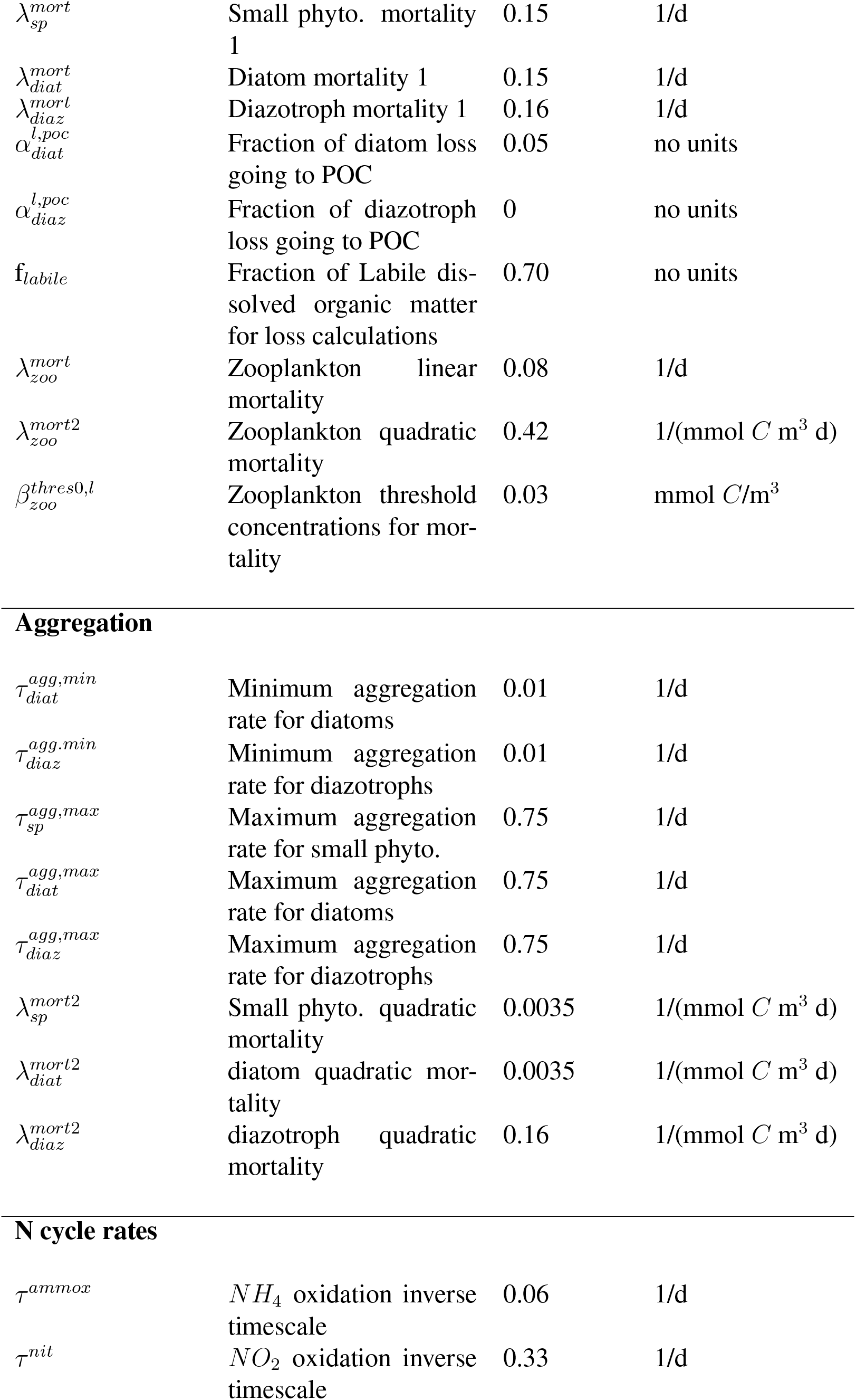
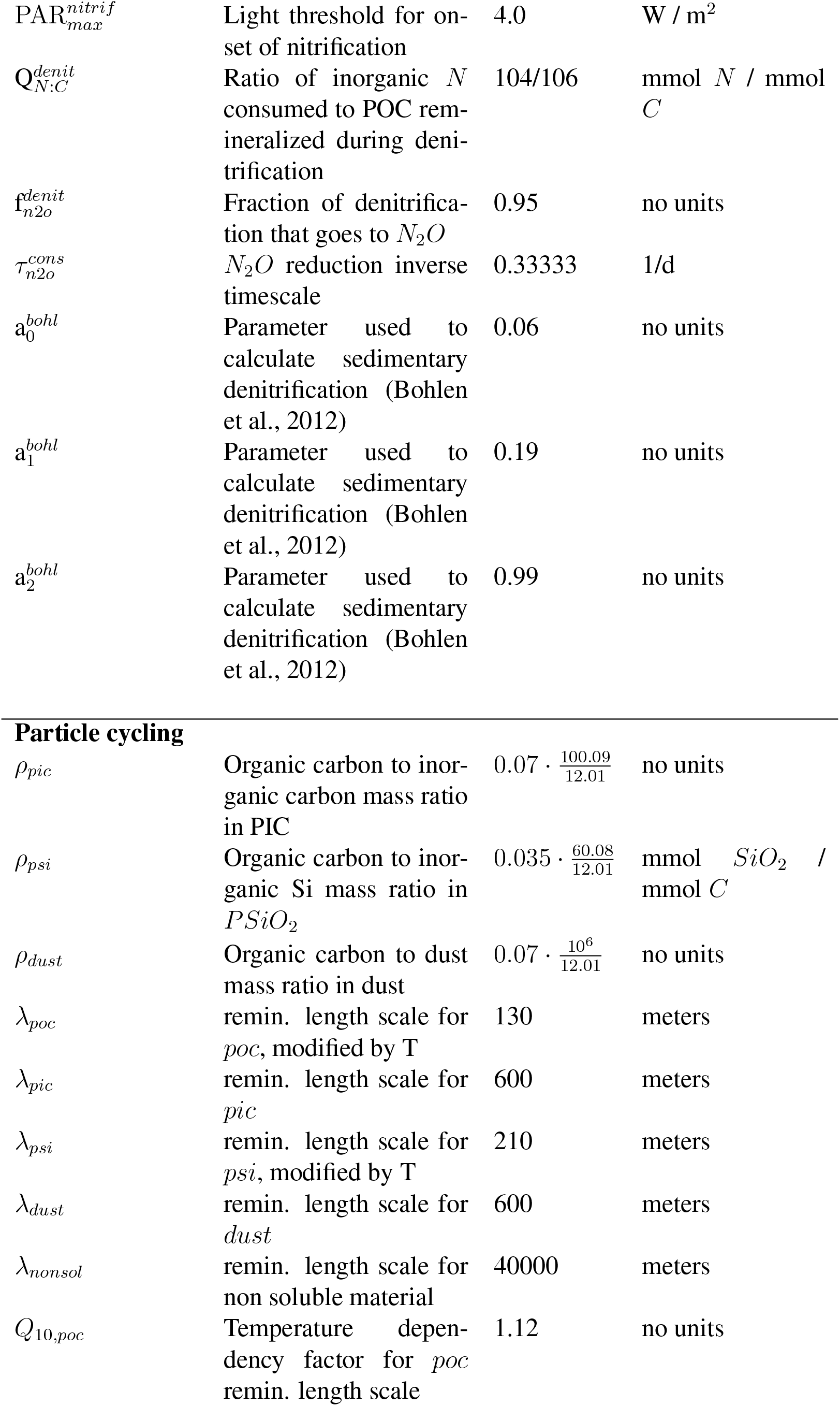
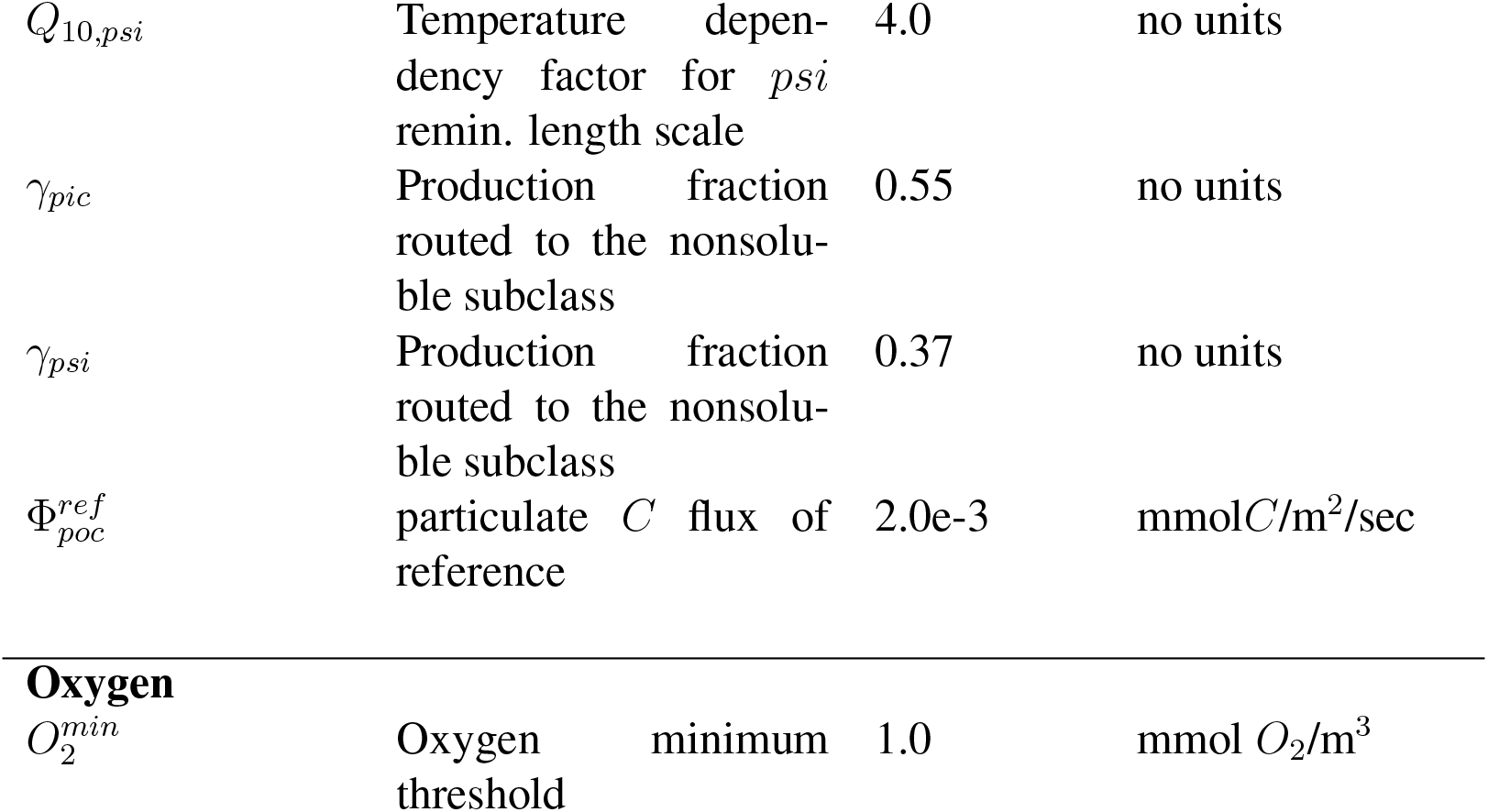

### 5.3 Model equations

#### 5.3.1 Tracer equations

Here the symbol d /dt denotes the sum of the local time derivative and the physical transport.

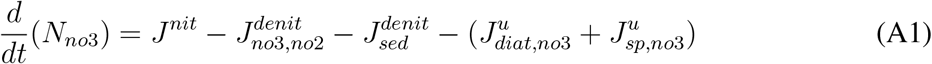

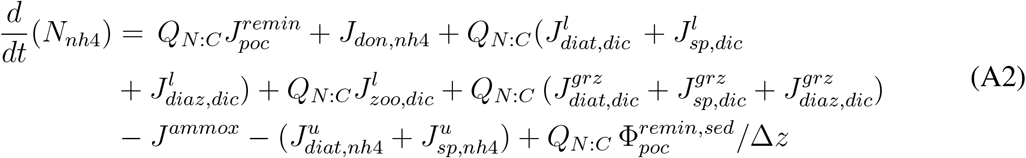

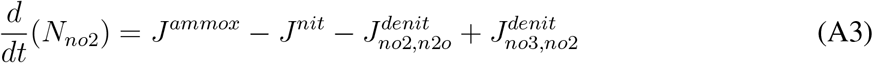

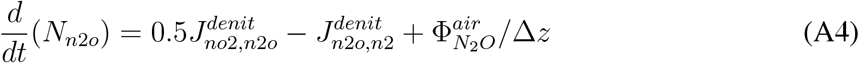

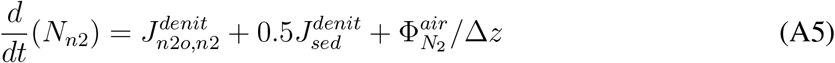

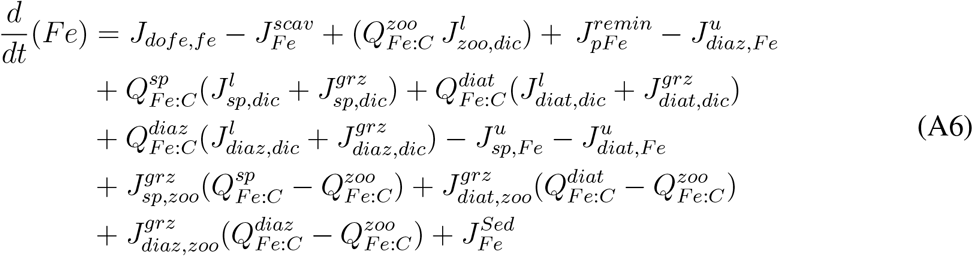

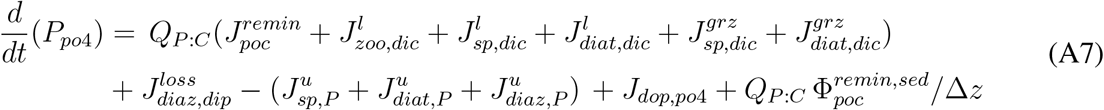

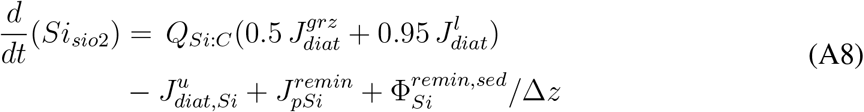

If (*O*_2_ > *O*_2_*min*)

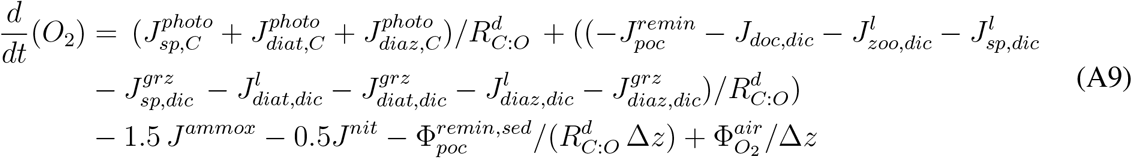

If (*O*_2_ ≤ *O*_2_*min*)

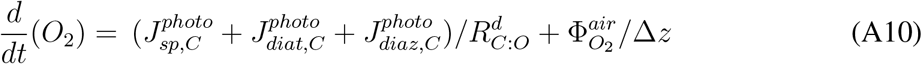

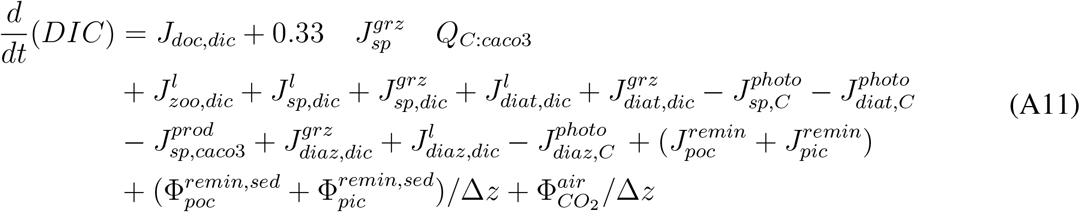

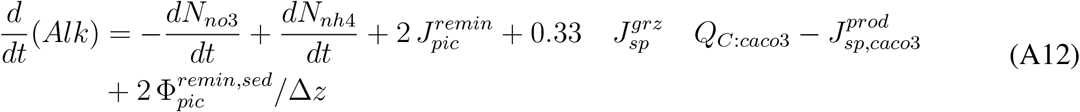

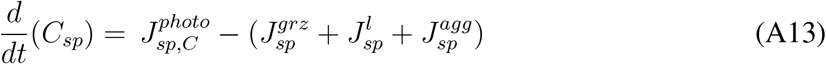

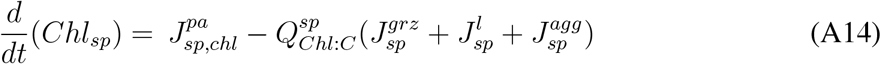

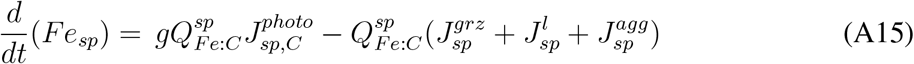

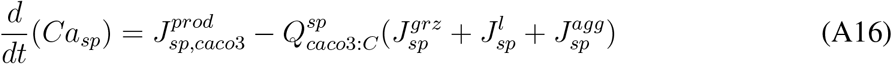

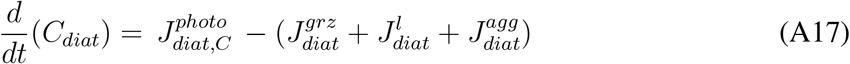

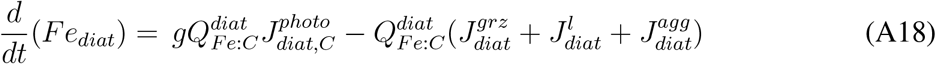

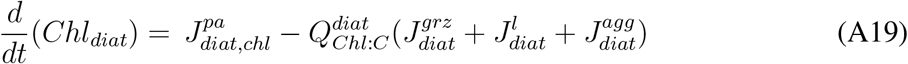

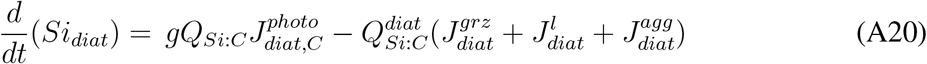

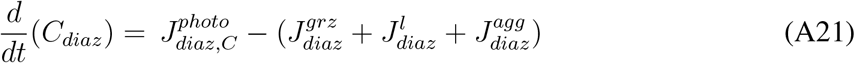

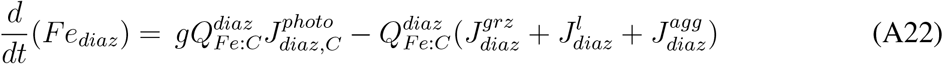

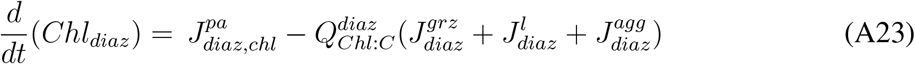

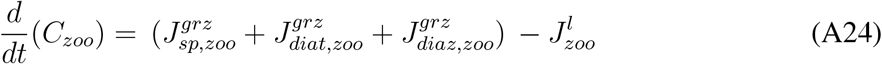

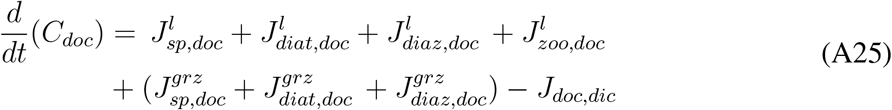

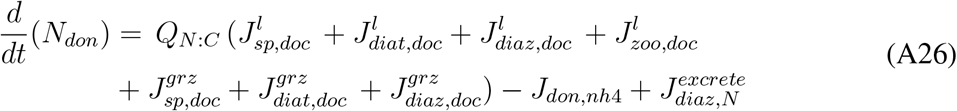

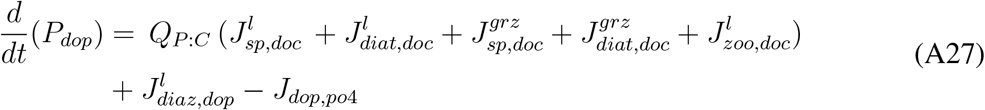

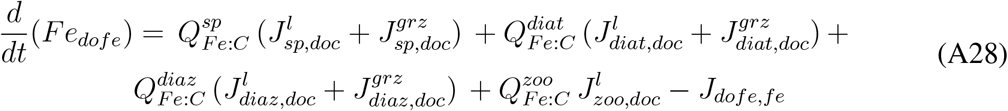

#### 5.3.2 Treatment of particulate organic matter

##### General model

Particulate organic matter is produced and instantaneously distributed over the depth of the water column following the exponential solution to the steady-state 1-dimensional production-remineralization equation:

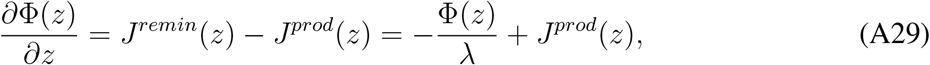

where Φ is a flux, λ is the remineralization length-scale, *J^prod^* is the known production rate within the layer, and *J^remm^* the remineralization rate within the layer, which needs to be determined. For a single layer, assuming the flux at the top of the layer Φ(k) is known, and the production *J^prod^*(*k*) is constant within the layer, the solution to equation A29, can be cast to determine the flux out of the layer, Φ(*k* — 1), for each element *i*:

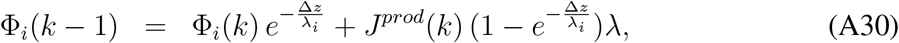

Particulate organic carbon (POC) is partitioned between a free and mineral associated component:

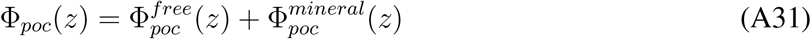

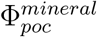 can be associated with CaCO_3_, SiO_2_ or dust. Each mineral-associated POC flux is further partitioned into a “soluble” component, which remineralizes with the length-scale of the associated mineral, and a “non-soluble” component which remineralizes with a length-scale of 40,000 m.

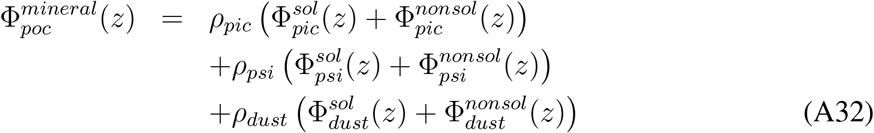

For all components of the fluxes except *Fe*, the flux out of the layer, Φ(k — 1), is computed first, with knowledge of the source within the layer, *J^prod^*(*k*), and of the remineralization length-scale. Remineralization in the layer is then calculated from conservation, *i.e.,* fromA29. Below, for each component, we list the equations used to determine the production terms *J^prod^*(*k*), followed by the fluxes out, Φ(k — 1), and finally the remineralization terms, *J^remm^*(*z*), which enter the tracer conservation equations.

##### Production

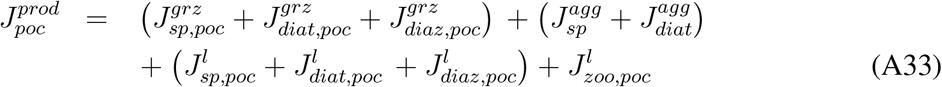

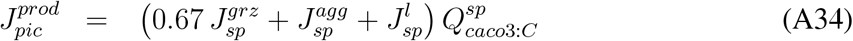

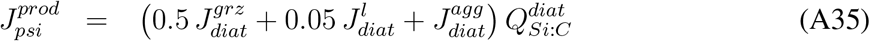

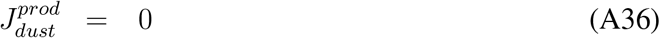

Available production for free POC is then:

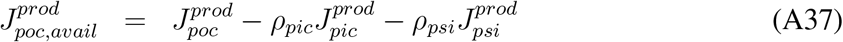

##### Fluxes out

Temperature dependency is used to modify the remineralization length scales of particulate organic carbon, POC, and opal, SiO_2_:

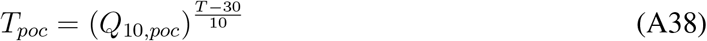

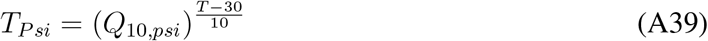

Free POC flux equation:

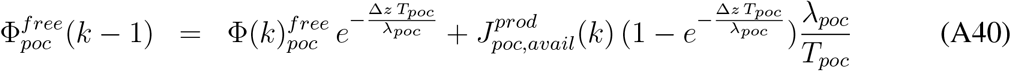

Soluble mineral-associated POC flux equation:

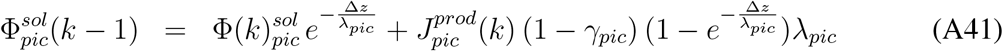

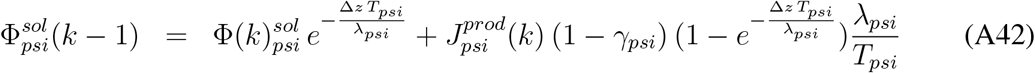

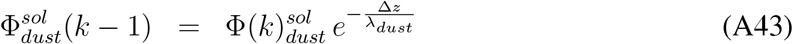

Non-soluble mineral-associated POC flux equation:

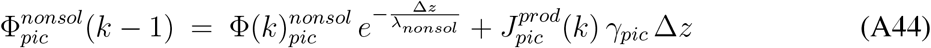

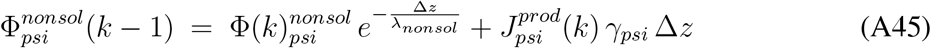

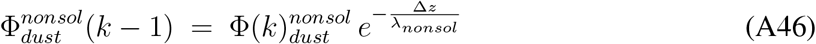

##### Remineralization

Remineralization is computed from conservation, i.e, A29:

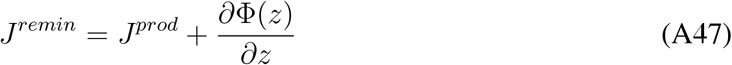

Numerically, for each individual layer, we have:

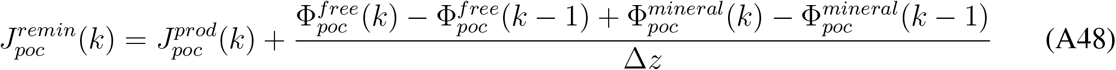

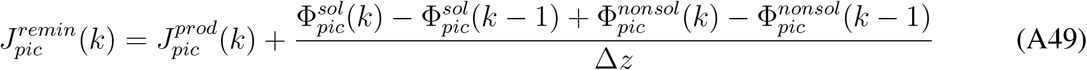

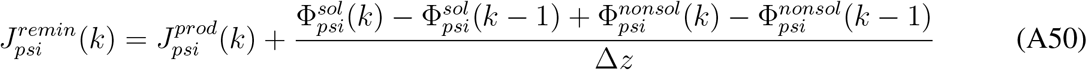

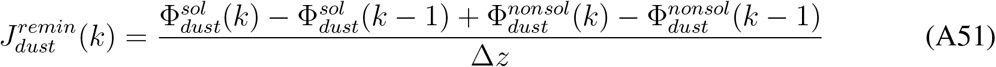

##### Particulate Fe

Production is as follows:

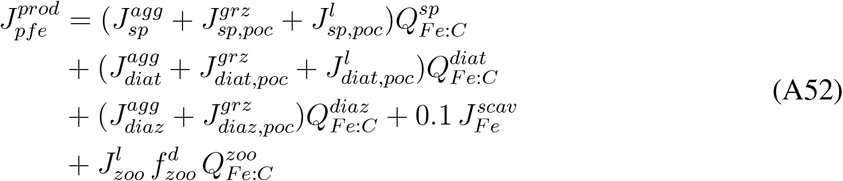

Particulate *Fe* remineralization is assumed to be proportional to POC remineralization plus a release from dust:

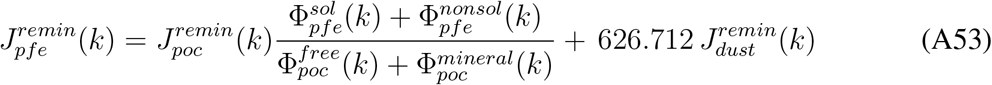

The flux out can then be computed from conservation:

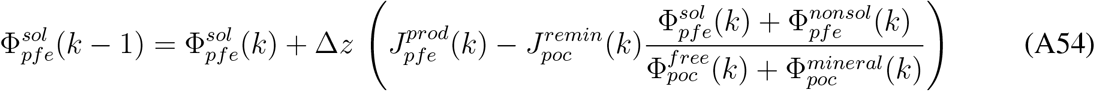

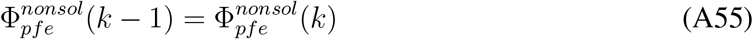

##### Bottom flux to sediment

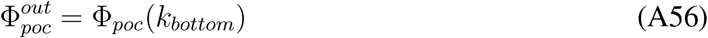

#### 5.3.3 Biogeochemical rates

##### Carbon

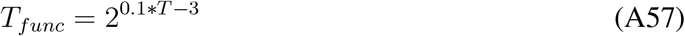

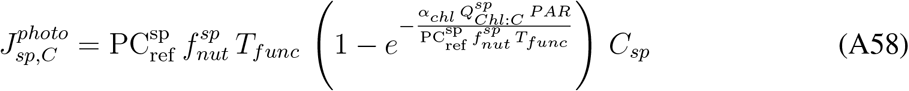

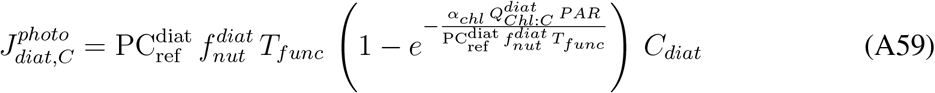

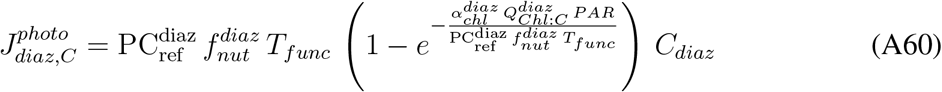

##### Remineralization

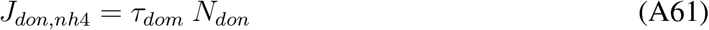

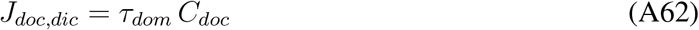

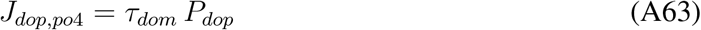

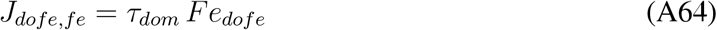

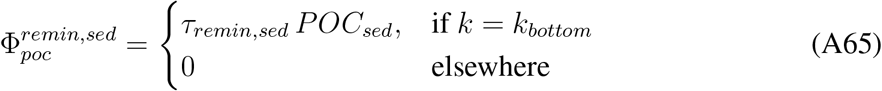

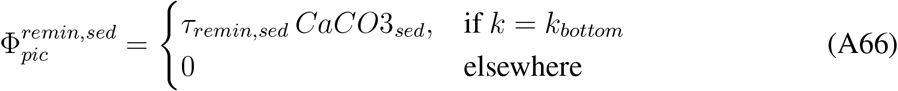

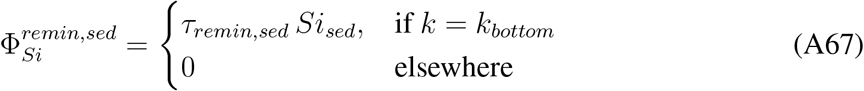

##### Nutrient limitation

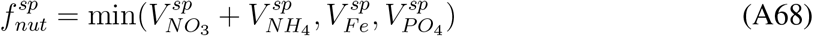

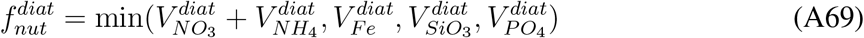

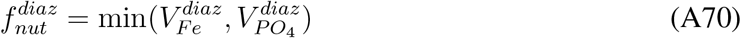

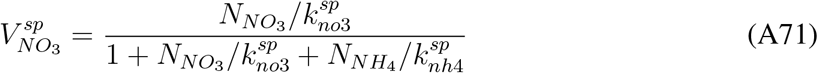

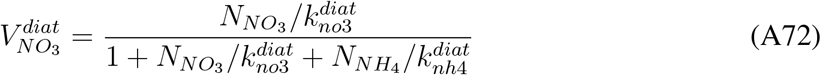

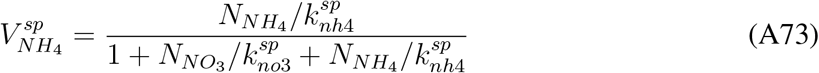

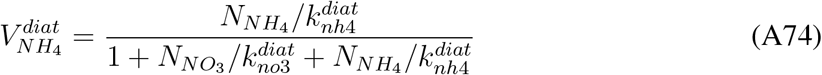

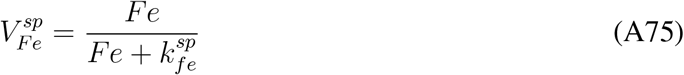

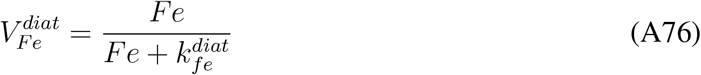

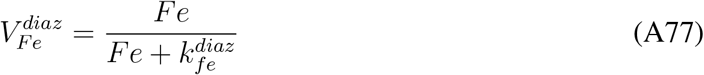

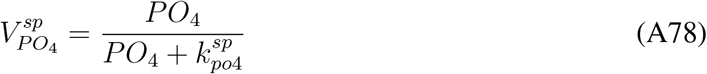

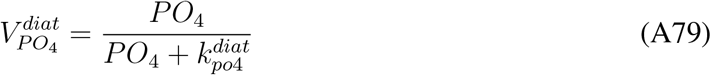

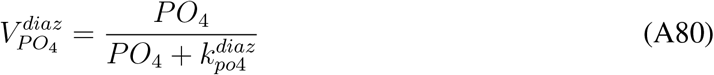

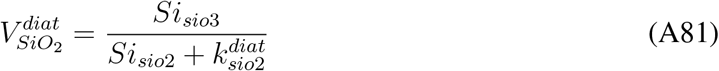

##### NO_3_ and NH_4_ uptake

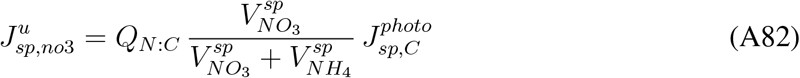

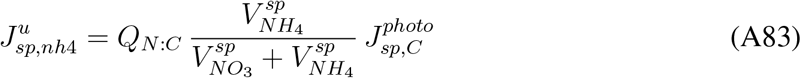

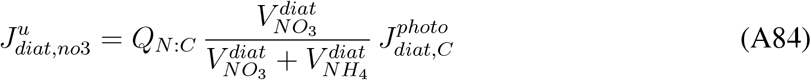

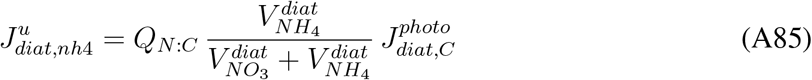

##### N_2_ fixation

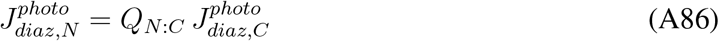

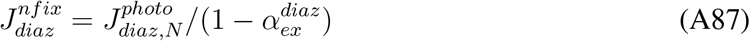

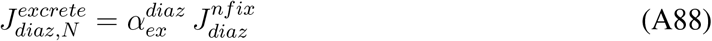

##### PO_4_ uptake

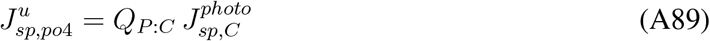

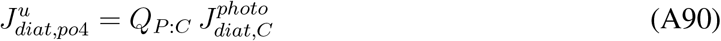

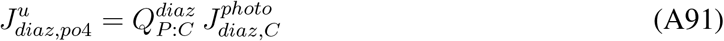

##### *Chl* stoichiometry and production

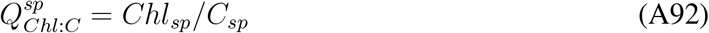

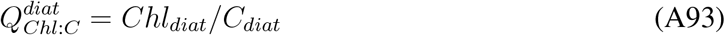

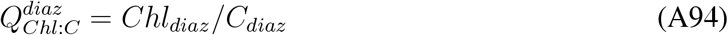

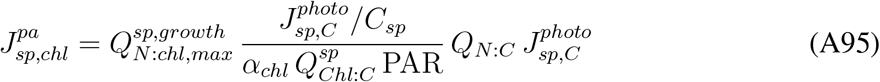

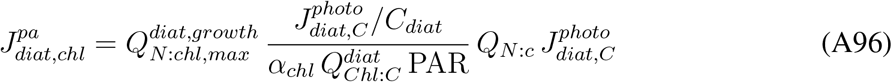

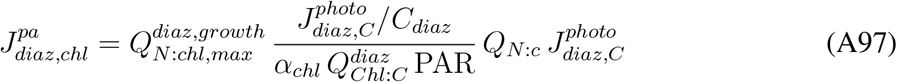

##### Light

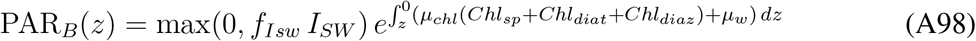

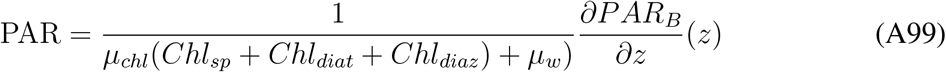

##### *Fe* stoichiometry and cycling

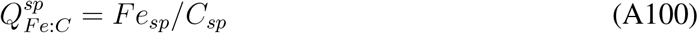

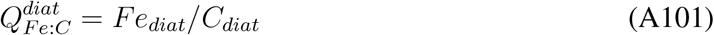

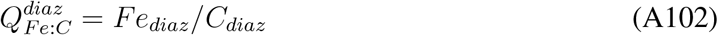

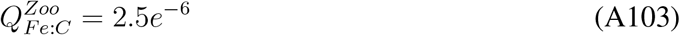

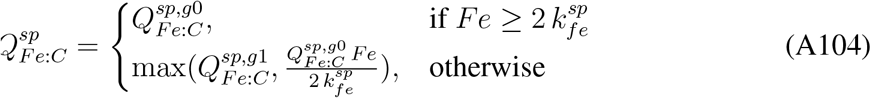

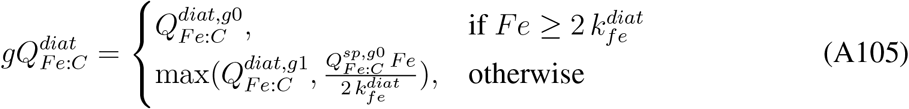

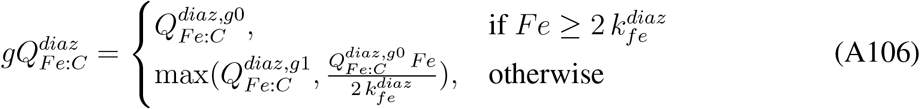

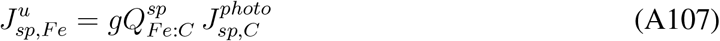

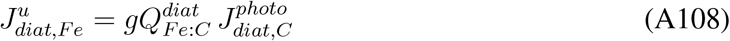

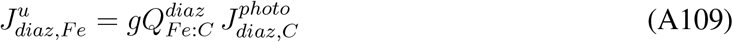

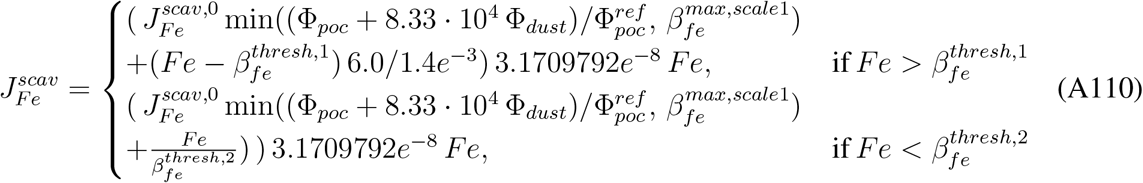

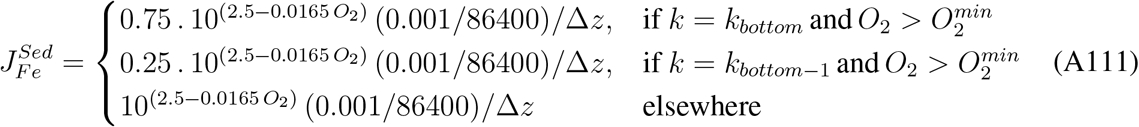

75% of 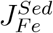 is released in the bottom layer and 25 % in the layer above.

##### *CaCO*_3_ stoichiometry production

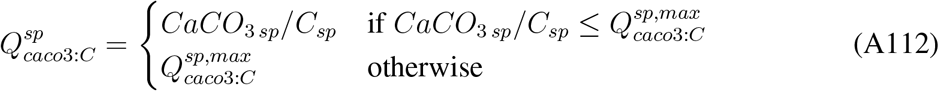

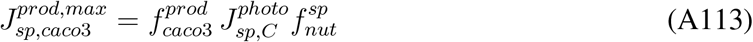

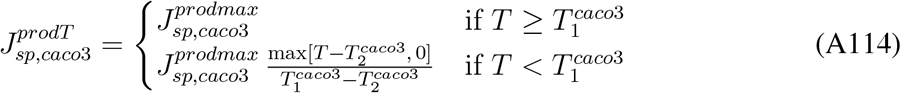

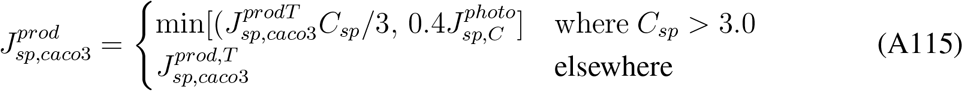

##### Si stoichiometry

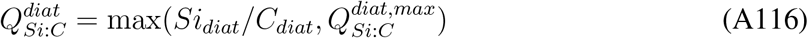

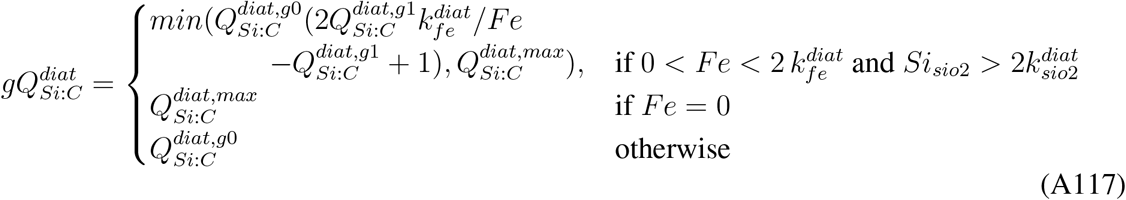

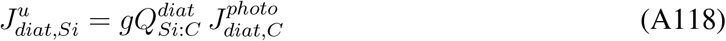

##### Grazing

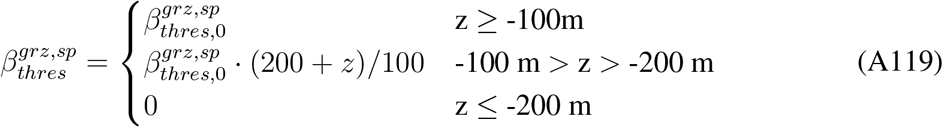

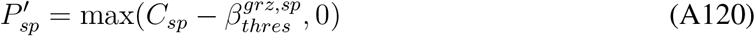

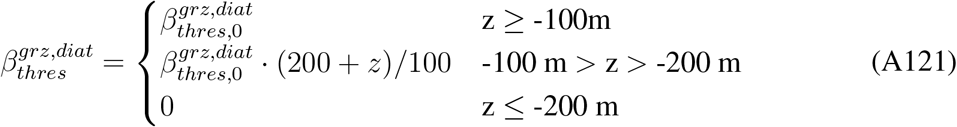

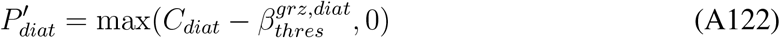

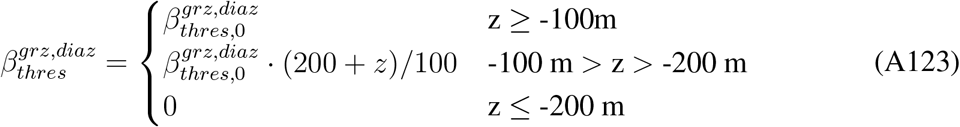

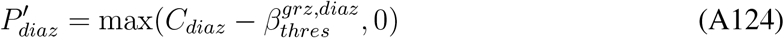

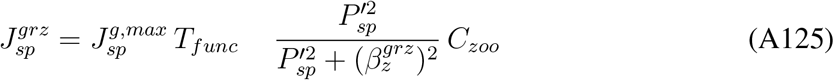

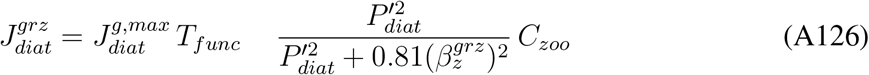

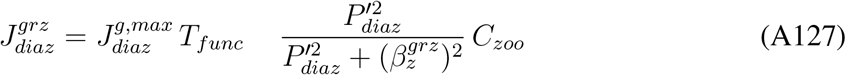

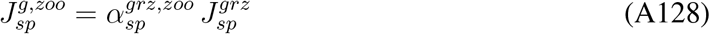

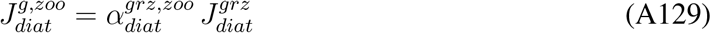

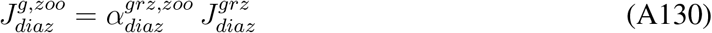

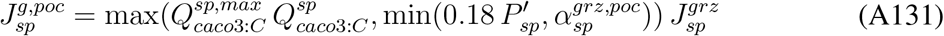

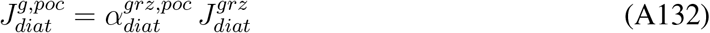

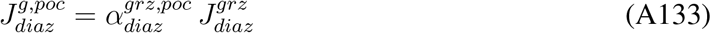

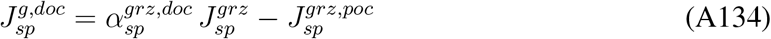

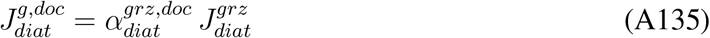

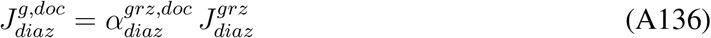

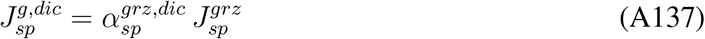

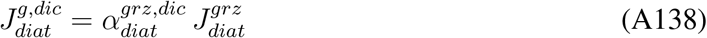

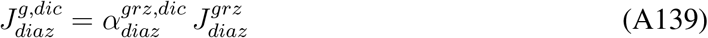

##### Losses

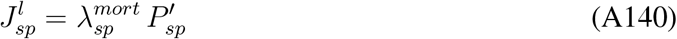

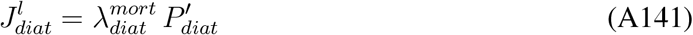

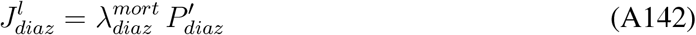

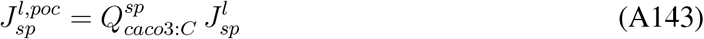

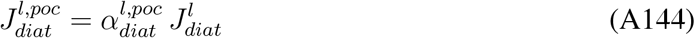

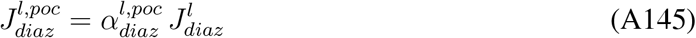

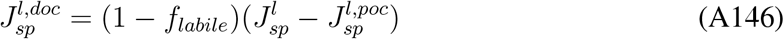

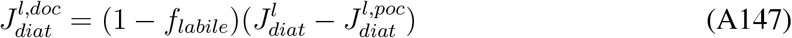

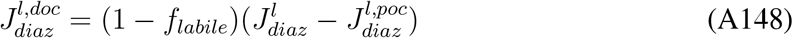

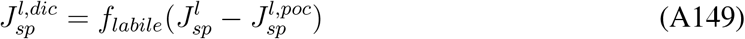

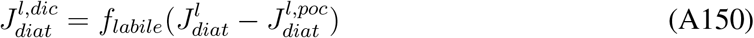

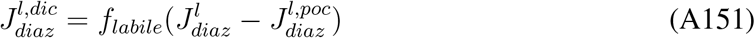

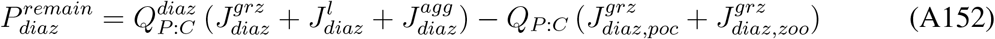

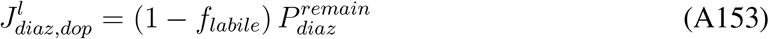

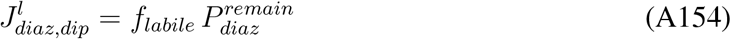

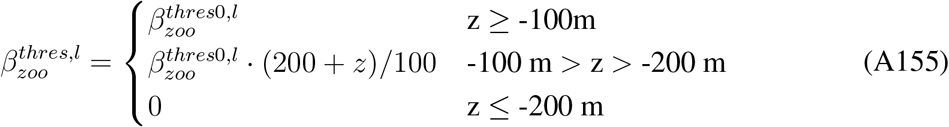

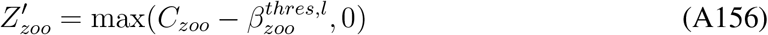

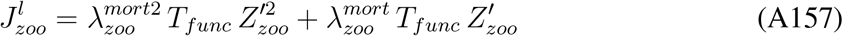

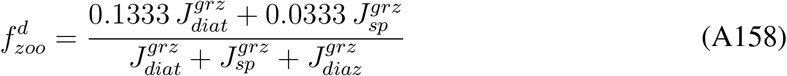

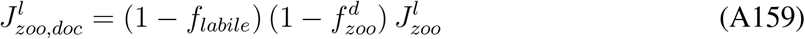

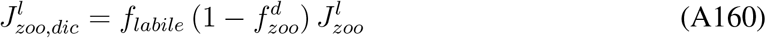

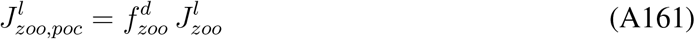

##### Aggregation

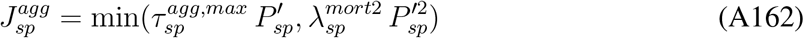

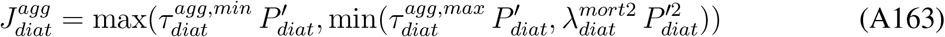

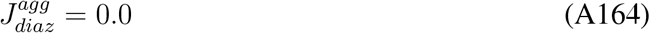

##### *N*-cycle rates

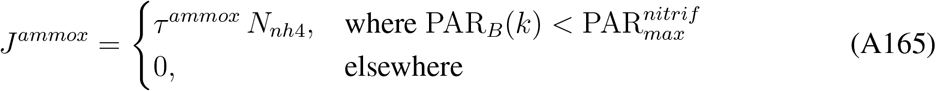

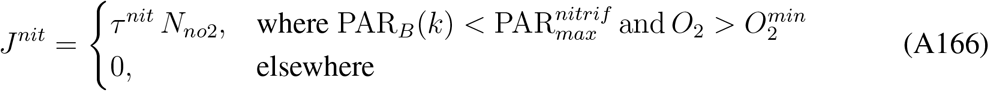

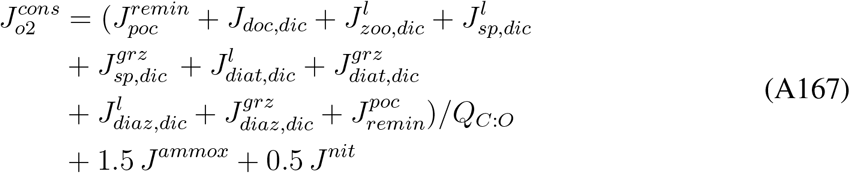

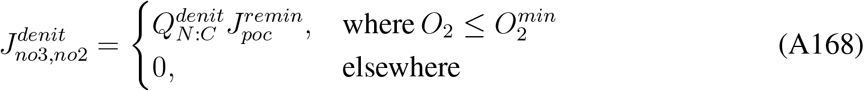

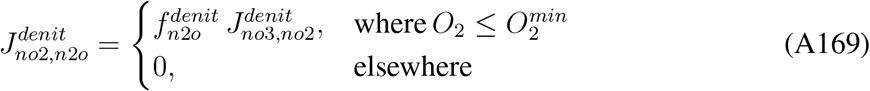

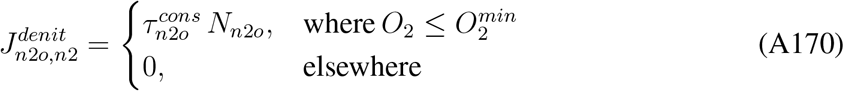

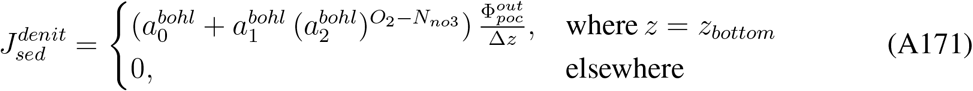

##### Air-sea fluxes

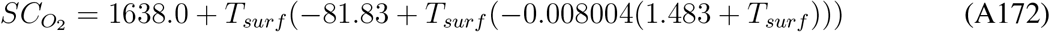

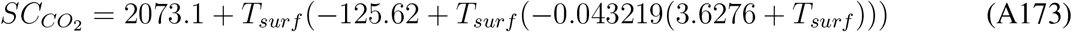

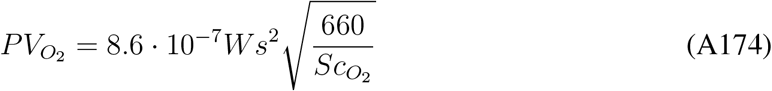

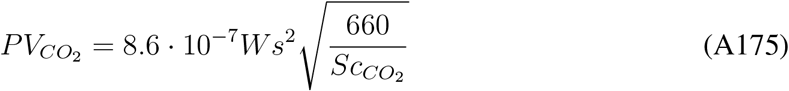

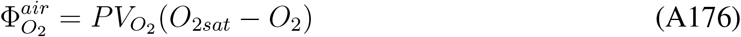

*O_2sat_* is the oxygen saturation concentration computed from Garcia and Gordon (1992), page 1310, eq. 8.

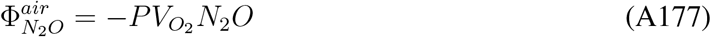

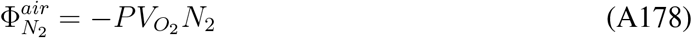

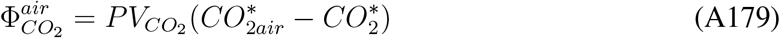

with 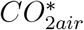 and 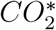 the concentrations corresponding to the partial pressure in air and water computed from Dickson and Goyet (1994) (SOP No. 3, p25-26).

##### Concentration in sediment

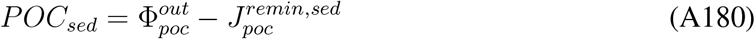

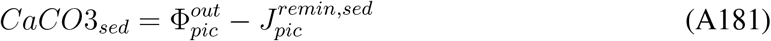

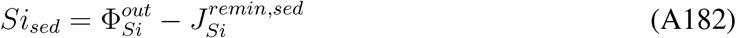

## References

Andersson, A. J., F. T Mackenzie, and N. R. Bates, 2008: Life on the margin: implications of ocean acidification on Mg-calcite, high latitude and cold-water marine calcifiers. Mar. Ecol. Progr. Ser., 373, 265–273.

Armstrong, R. A., C. Lee, J. I. Hedges, S. Honjo, and S. G. Wakeham, 2001: A new, mechanistic model for organic carbon fluxes in the ocean based on the quantitative association of POC with ballast minerals. Deep-Sea Res. II, 49, 219–236.

Bakker, D., and coauthors, 2016: A multi-decade record of high-quality fCO2 data in version 3 of the Surface Ocean CO2 Atlas (SOCAT). Earth System Science Data, 8, 383–413, doi:https://doi.org/10.5194/essd-8-383-2016.

Bakun, A., 1990: Global climate change and intensification of coastal ocean upwelling. Science, 247, 198–201.

Banas, N., and B. Hickey, 2008: Why is the northern end of the California Current System so productive? Oceanography, 21, 90–107.

Behrenfeld, M. J., and P. G. Falkowski, 1997: Photosynthetic rates derived from satellite-based chlorophyll concentration. Limnol. Oceanogr., 42, 1–20.

Bianchi, D., T. S. Weber, R. Kiko, and C. Deutsch, 2018: Global niche of marine anaerobic metabolisms expanded by particle microenvironments. Nature Geosci., 11, 263–268, doi.org/10.1038/s41561-018-0081-0.

Bograd, S. J., and R. J. Lynn, 2001: Physical-biological coupling in the California Current during the 1997-99 El Niño-La Niña cycle. Geophys. Res. Lett., 28, 275–278.

Bohlen, L., A. W. Dale, and K. Wallmann, 2012: Simple transfer functions for calculating benthic fixed nitrogen losses and C:N:P regeneration ratios in global biogeochemical models. Global Biogeochem. Cycles, 26, GB3029, 1–16.

Buil, M. P., and E. D. Lorenzo, 2017: Decadal dynamics and predictability of oxygen and subsurface tracers in the California Current System. Geophys. Res. Lett., 44, 4204–4213, doi:10.1002/2017GL072931.

Busch, D. S., and P. McElhany, 2016: Estimates of the direct effect of seawater pH on the survival rate of species groups in the California Current ecosystem. PLOS One, doi:10.1371/journal.pone.0160669.

Capet, X., J. McWilliams, M. Molemaker, and A. Shchepetkin, 2008: Mesoscale to submesoscale transition in the California Current System. Part I: Flow structure, eddy flux, and observational tests. Journal of Physical Oceanography, 38 (1), 29–43.

Carr, M.-E., and E. J. Kearns, 2003: Production regimes in four Eastern Boundary Current Systems. Deep Sea Res. II, 50, 3199–3221.

Chan, F., J. Barth, J. Lubchenco, A. Kirincich, H. Weeks, W. T. Peterson, and B. Menge, 2008: Emergence of anoxia in the California Current large marine ecosystem. Science, 319, 920–920.

Chavez, F., and Coauthors, 2002: Biological and chemical consequences of the 1997-1998 El Nino in central California waters. Progr. Oceanogr., 54, 205–232.

Chavez, F. P., and M. Messie, 2009: A comparison of eastern boundary upwelling ecosystems. Progr. Oceanogr, 83, 80–96.

Chavez, F. P., J. Ryan, S. E. Lluch-Cota, and M. Ñiquen, 2003: From anchovies to sardines and back: multidecadal change in the Pacific Ocean. Science, 299, 217–221.

Collins, W., and Coauthors, 2011: Development and evaluation of an Earth-System model-HadGEM2. Geosci. Model Devel., 4, 1051–1075.

Davis, K. A., N. S. Banas, S. N. Giddings, S. A. Siedlecki, P. MacCready, E. J. Lessard, R. M. Kudela, and B. M. Hickey, 2014: Estuary-enhanced upwelling of marine nutrients fuels coastal productivity in the U.S. Pacific Northwest. J. Geophys. Res. Oceans, 119, 8778–8799.

Deutsch, C., H. Brix, T. Ito, H. Frenzel, and L. Thompson, 2011: Climate-forced variability of ocean hypoxia. Science, 333, 336–339.

Deutsch, C., S. Emerson, and L. Thompson, 2006: Fingerprints of climate change in North Pacific oxygen. Geophys. Res. Lett., 32, 1–4.

Deutsch, C., A. Ferrel, B. Seibel, H.-O. Portner, and R. B. Huey, 2015: Climate change tightens a metabolic constraint on marine habitats. Science, 348, 1132–1135.

Di Lorenzo, E., and Coauthors, 2008: North Pacific Gyre Oscillation links ocean climate and ecosystem change. Geophys. Res. Lett., 35, doi:10.1029/2007GL032838.

Dickson, A. G., and C. Goyet, 1994: Handbook of methods for the analysis of the various parameters of the carbon dioxide system in sea water. version 2.

Doney, S. C., N. Mahowald, I. Lima, R. A. Feely, F. T. Mackenzie, J.-F. Lamarque, and P. J. Rasch, 2007: Impact of anthropogenic atmospheric nitrogen and sulfur deposition on ocean acidification and the inorganic carbon system. Proc. Nat. Acad. Sci., 104, 14580–14585.

Ducklow, H. W., S. C. Doney, and D. K. Steinberg, 2009: Contributions of long-term research and time-series observations to marine ecology and biogeochemistry. Annu. Rev. Marine. Sci., 1, 279–302.

Dugdale, R. C., F. P. Wilkerson, R. T. Barber, and F. P. Chavez, 1992: Estimating new production in the equatorial Pacific Ocean at 150 W. J. Geophys. Res. Oceans, 97, 681–686.

Dunne, J. P., R. A. Armstrong, A. Gnanadesikan, and J. L. Sarmiento, 2005: Empirical and mechanistic models for the particle export ratio. Global Biogeochem. Cycles, 19, doi:10.1029/2004GB002390.

Durski, S. M., J. A. Barth, J. C. McWilliams, H. Frenzel, and C. Deutsch, 2017: The influence of variable slope-water characteristics on dissolved oxygen levels in the northern California Current System. J. Geophys. Res. Oceans, 122, 7674–7697, doi:10.1002/2017JC013089.

Dussin, R., E. N. Curchitser, C. A. Stock, and N. V. Oostende, 2019: Biogeochemical drivers of changing hypoxia in the California Current Ecosystem. Deep-Sea Res. II, 169-170, doi:10.1016/j.dsr2.2019.05.013.

Eppley, R. W., F. P. Chavez, and R. T. Barber, 1992: Standing stocks of particulate carbon and nitrogen in the equatorial Pacific at 150° W. J. Geophys. Res. Oceans, 97, 655–661.

FAO, 2009: State of the World’s Fisheries and Aquaculture 2008. Food and Agriculture Organization of the United Nations.

Feely, R. A., C. L. Sabine, J. M. Hernandez-Ayon, D. Ianson, and B. Hales, 2008: Evidence for upwelling of corrosive “acidified” water onto the continental shelf. Science, 320, 1490–1492.

Fiechter, J., E. N. Curchitser, C. A. Edwards, F. Chai, N. L. Goebel, and F. P. Chavez, 2014: Airsea CO_2_fluxes in the California Current: Impacts of model resolution and coastal topography. Global Biogeochem. Cycles, 28, doi:10.1002/2013GB004683.

Firme, G. F., E. L. Rue, D. A. Weeks, K. W. Bruland, and D. A. Hutchins, 2003: Spatial and temporal variability in phytoplankton iron limitation along the California coast and consequences for Si, N, and C biogeochemistry. Global Biogeochem. Cycles, 17, doi:10.1029/2001GB001824.

Frenger, I., D. Bianchi, C. Stuhrenberg, A. Oschlies, J. Dunne, C. Deutsch, E. Galbraith, and F. Schutte, 2018: Biogeochemical role of subsurface coherent eddies in the ocean: Tracer cannonballs, hypoxic storms, and microbial stewpots? Global Biogeochem. Cycles, 32, doi:10.1002/2017GB005743.

Frischknecht, M., M. Münnich, and N. Gruber, 2018: Origin, transformation, and fate: The threedimensional biological pump in the California Current System. J. Geophys. Res. Oceans, 123, doi:10.1029/2018JC013934.

Garcia, H. E., and L. I. Gordon, 1992: Oxygen solubility in seawater: Better fitting equations. Limnol. Oceanogr., 37 (6), 1307–1312.

Garcia-Reyes, M., J. L. Largier, and W. J. Sydeman, 2014: Synoptic-scale upwelling indices and predictions of phyto- and zooplankton populations. Progr. Oceanogr., 120, 177–188.

Geider, R. J., H. L. Maclntyre, and T. M. Kana, 1998: A dynamic regulatory model of phytoplanktonic acclimation to light, nutrients, and temperature. Limnol. Oceanogr., 43, 679–694.

Gruber, N., C. Hauri, Z. Lachkar, D. Loher, T. L. Frölicher, and G.-K. Plattner, 2012: Rapid progression of ocean acidification in the California Current System. Science, 337, 220–223.

Gruber, N., Z. Lachkar, H. Frenzel, P. Marchesiello, M. Münnich, J. C. McWilliams, T. Nagai, and G.-K. Plattner, 2011: Eddy-induced reduction of biological production in eastern boundary upwelling systems. Nature Geosci., 4, 787–792.

Hales, B., P. G. Strutton, M. Saraceno, R. Letelier, T. Takahashi, R. Feely, C. Sabine, and F. Chavez, 2012: Satellite-based prediction of *pCO_2_* in coastal waters of the eastern North Pacific. Progr Oceanogr., doi:10.1016/j.pocean.2012.03.001.

Howard, E. M., and Coauthors, 2020: Climate-driven aerobic habitat loss in the California Current System. Science Adv., 6, doi:eaay3188.

Ito, T., and C. Deutsch, 2010: A conceptual model for the temporal spectrum of oceanic oxygen variability. Geophys. Res. Lett., 37, doi:10.1029/2009GL041595.

Ito, T., M. C. Long, C. Deutsch, S. Minobe, and D. Sun, 2019: Mechanisms of low-frequency oxygen variability in the North Pacific. Global Biogeochem. Cycles, 33, doi:10.1029/2018GB005987.

Jacox, M. G., C. A. Edwards, E. L. Hazen, and S. J. Bograd, 2018: Coastal Upwelling Revisited: Ekman, Bakun, and Improved Upwelling Indices for the U.S. West Coast. J. Geophys. Res. Oceans, 123, 7332–7350, doi:10.1029/2018JC014187.

Kahru, M., R. Kudela, M. Manzano-Sarabia, and B. G. Mitchell, 2009: Trends in primary production in the California Current detected with satellite data. J. Geophys. Res. Oceans, 114, doi:10.1029/2008JC004979.

Kessouri, F., D. Bianchi, L. Renault, J. C. McWilliams, H. Frenzel, and C. Deutsch, 2020: Submesoscale currents modulate the seasonal cycle of nutrients and productivity in the California Current System. Global Biogeochem. Cycles, 34, doi:10.1029/2020GB006578.

Key, R. M., and Coauthors, 2004: A global ocean carbon climatology: Results from Global Data Analysis Project (GLODAP). Global Biogeochem. Cycles, 18, doi:10.1029/GB002247.

Kudela, R. M., and Coauthors, 2008: New insights into the controls and mechanisms of plankton productivity in coastal upwelling waters of the Northern California Current System. Oceanography, 21, 46–59.

Kwon, E. Y., C. Deutsch, S.-P. Xie, S. Schmidtko, and Y.-K. Cho, 2016: The North Pacific oxygen uptake rates over the past half century. J. Climate, 29, doi:10.1175/JCLI-D-14-00157.1.

Large, W. B., 2006: Surface fluxes for practitioners of global ocean data assimilation. Ocean Weather Forecasting, Springer, 229–270.

Lauvset, S. K., and coauthors, 2016: A new global interior ocean mapped climatology: the 1°× 1°GLODAP version 2. Earth System Science Data, 8, 325–340, doi:10.5194/essd-8-325-2016.

Locarnini, R., and Coauthors, 2013: World Ocean Atlas 2013: Temperature. NOAA NESDIS, 1.

Long, M. C., C. Deutsch, and T. Ito, 2016: Finding forced trends in oceanic oxygen. Global Biogeochem. Cycles, 30, 381–397, doi:10.1002/2015gb005310.

Mahowald, N. M., M. Yoshioka, W. D. Collins, A. J. Conley, D. W. Fillmore, and D. B. Coleman, 2006: Climate response and radiative forcing from mineral aerosols during the last glacial maximum, pre-industrial, current and doubled-carbon dioxide climates. Geophys. Res. Lett., 33, doi:10.1029/2006GL026126.

Mantua, N. J., S. R. Hare, Y. Zhang, J. M. Wallace, and R. C. Francis, 1997: A Pacific interdecadal climate oscillation with impacts on salmon production. Bull. Am. Meteor. Soc., 78, 1069–1080.

McClatchie, S., R. Goericke, R. Cosgrove, G. Auad, and R. Vetter, 2010: Oxygen in the Southern California Bight: Multidecadal trends and implications for demersal fisheries. Geophys. Res. Lett., 37, doi:10.1029/2010GL044497.

Mecking, S., C. Langdon, R. A. Feely, C. L. Sabine, C. A. Deutsch, and D.-H. Min, 2008: Climate variability in the North Pacific thermocline diagnosed from oxygen measurements: An update based on the US CLIVAR/CO_2_ Repeat Hydrography cruises. Global Biogeochem. Cycles, 22, doi:10.1029/2007GB003101.

Meinvielle, M., and G. C. Johnson, 2013: Decadal water-property trends in the California Undercurrent, with implications for ocean acidification. J. Geophys. Res. Oceans, 118, 6687–6703.

Middelburg, J. J., K. Soetaert, P. M. Herman, and C. H. Heip, 1996: Denitrification in marine sediments: A model study. Global Biogeochem. Cycles, 10, 661–673.

Moore, J. K., S. C. Doney, J. A. Kleypas, D. M. Glover, and I. Y. Fung, 2002: An intermediate complexity marine ecosystem model for the global domain. Deep-Sea Res. II, 49, 403–462.

Moore, J. K., S. C. Doney, and K. Lindsay, 2004: Upper ocean ecosystem dynamics and iron cycling in a global three-dimensional model. Global Biogeochem. Cycles, 18, GB4028, doi:10.1029/2004GB002220.

Munro, D. R., P. D. Quay, L. W. Juranek, and R. Goericke, 2013: Biological production rates off the Southern California coast estimated from triple O_2_ isotopes and O_2_: Ar gas ratios. Limnol. Oceanogr., 58, 1312–1328.

Murray, J. W., J. Young, J. Newton, J. Dunne, T. Chapin, B. Paul, and J. J. McCarthy, 1996: Export flux of particulate organic carbon from the central equatorial Pacific determined using a combined drifting trap-^234^ Th approach. Deep-Sea Res. II, 43, 1095–1132.

Nagai, T., N. Gruber, H. Frenzel, Z. Lachkar, J. C. McWilliams, and G. K. Plattner, 2015: Dominant role of eddies and filaments in the offshore transport of carbon and nutrients in the California Current System. J. Geophys. Res. Oceans, 120, 5318–5341.

Palevsky, H. I., P. D. Quay, and D. P. Nicholson, 2016: Discrepant estimates of primary and export production from satellite algorithms, a biogeochemical model, and geochemical tracer measurements in the North Pacific Ocean. Geophys. Res. Lett., 43, 8645–8653.

Pierce, D. W., P. J. Gleckler, T. P. Barnett, B. D. Santer, and P. J. Durack, 2012: The fingerprint of human-induced changes in the ocean’s salinity and temperature fields. Geophys. Res. Lett., 39, L21704, doi:10.1029/2012GL053389.

Renault, L., C. Deutsch, J. C. McWilliams, F. Colas, H. Frenzel, and J. H. Liang, 2016a: Partial decoupling of primary productivity from upwelling in the California Current System. Nature Geosci., 9, 505–508.

Renault, L., J. C. McWilliams, A. Jousse, C. Deutsch, H. Frenzel, F. Kessouri, and R. Chen, 2021: The physical structure and behavior of the California Current System. Progr Oceanogr., submitted.

Renault, L., M. J. Molemaker, J. C. McWilliams, A. F. Shchepetkin, F. Lemarie, D. Chelton, S. Illig, and A. Hall, 2016b: Modulation of wind work by oceanic current interaction with the atmosphere. J. Phys. Ocean., 46, 1685–1704.

Roemmich, D., and T. McCallister, 1989: Large scale circulation of the North Pacific Ocean. Progr Oceanogr., 22, 171–204.

Rykaczewski, R. R., and D. M. Checkley, 2008: Influence of ocean winds on the pelagic ecosystem in upwelling regions. Proc. Nat. Acad. Sci., 105, 1965–1970.

Severmann, S., J. McManus, W. M. Berelson, and D. E. Hammond, 2010: The continental shelf benthic iron flux and its isotope composition. Geochimica et Cosmochimica Acta, 74, 39844004.

Shchepetkin, A. F., and J. C. McWilliams, 2005: The Regional Oceanic Modeling System (ROMS): A split-explicit, free-surface, topography-following-coordinate oceanic model. Ocean Modelling, 9, 347–404.

Stukel, M. R., M. R. Landry, C. R. Benitez-Nelson, and R. Goericke, 2011: Trophic cycling and carbon export relationships in the California Current Ecosystem. Limnol. Oceanogr., 56, 18661878.

Stukel, M. R., and Coauthors, 2017: Mesoscale ocean fronts enhance carbon export due to gravitational sinking and subduction. Proc. Nat. Acad. Sci., 114, 1252–1257.

Turi, G., Z. Lachkar, and N. Gruber, 2014: Spatiotemporal variability and drivers of pCO_2_ and air-sea CO_2_ fluxes in the California Current System: an eddy-resolving modeling study. Biogeosci., 11, 671–690, doi:https://doi.org/10.5194/bg-11-671-2014.

Turi, G., and Coauthors, 2018: Response of O_2_ and pH to ENSO in the California Current System in a high-resolution global climate model. Ocean Science, 14, 69–86, doi:10.5194/os-14-69-2018.

van Heuven, S., D. Pierrot, J. Rae, E. Lewis, and D. Wallace, 2011: MATLAB Program Developed for CO_2_System Calculations. ORNL/CDIAC-105b. Carbon Dioxide Information Analysis Center, Oak Ridge National Laboratory, U.S. Department of Energy, Oak Ridge, Tennessee, doi:10.3334/CDIAC/otg.CO2SYSATLAB_v1.1.

Wanninkhof, R., 1992: Relationship between wind speed and gas exchange over the ocean. J. Geophys. Res. Oceans, 97, 7373–7382, doi:10.1029/92JC00188.

Westberry, T., M. Behrenfeld, D. Siegel, and E. Boss, 2008: Carbon-based primary productivity modeling with vertically resolved photoacclimation. Global Biogeochem. Cycles, 22, GB2024, doi:10.1029/2007GB003078.

Zweng, M., and Coauthors, 2013: World Ocean Atlas 2013,: Salinity. NOAA NESDIS, 2.

